# Organ- and function-specific anatomical organization of the vagus nerve supports fascicular vagus nerve stimulation

**DOI:** 10.1101/2022.03.07.483266

**Authors:** Naveen Jayaprakash, Weiguo Song, Viktor Toth, Avantika Vardhan, Todd Levy, Jacquelyn Tomaio, Khaled Qanud, Ibrahim Mughrabi, Yao-Chuan Chang, Moontahinaz Rob, Anna Daytz, Adam Abbas, Zeinab Nassrallah, Bruce T. Volpe, Kevin J. Tracey, Yousef Al-Abed, Timir Datta-Chaudhuri, Larry Miller, Mary F. Barbe, Sunhee C. Lee, Theodoros P. Zanos, Stavros Zanos

**Affiliations:** Feinstein Institutes for Medical research, Manhasset, New York; Donald and Barbara Zucker School of Medicine at Hofstra/Northwell, Hempstead, New York; Temple University, Philadelphia, Pennsylvania; Elmezzi Graduate School of Molecular Medicine, Manhasset, NY

**Author notes:** These authors contributed equally to this work (Naveen Jayaprakash, Weiguo Song, Viktor Toth).

## Abstract

Vagal fibers travel inside fascicles and form branches to innervate organs and regulate organ functions. Vagus nerve stimulation (VNS) therapies activate fibers non-selectively, often resulting in reduced efficacy and side effects from non-targeted organs. Transverse and longitudinal arrangement of fibers according to functions they mediate and organs they innervate is unknown, however it is crucial for selective VNS. Using microcomputed tomography, we found that, in swine, fascicles are arranged in 2 overlapping axes, with sensory and motor fascicles separated cephalad and merging caudad, and larynx-, heart- and lung-specific fascicles separated caudad and progressively merging cephalad. Using immunohistochemistry, we found that the distribution of single fibers is highly nonuniform: myelinated afferents and efferents occupy separate fascicles, unmyelinated efferents co-localize with myelinated afferents, and small unmyelinated afferents are widely distributed. Using a multi-contact cuff electrode, we delivered fascicular cervical VNS in anesthetized and awake swine. Compound action potentials, from distinct fiber types, and organ responses, including laryngeal muscle, cough, breathing, heart rate and blood pressure responses are elicited in a radially asymmetric manner, with consistent angular separations. These results indicate that vagal fibers are anatomically organized according to functions they mediate and organs they innervate and can be asymmetrically activated by fascicular cervical VNS.

## Introduction

The autonomic nervous system maintains physiological homeostasis in organs and organ systems through neural reflexes. Autonomic reflexes are comprised of sensory neurons that detect changes in physiological parameters, and motor neurons that regulate organ functions in response to physiologic changes. With its thousands of afferent (sensory, directed from the periphery to the brain) and efferent fibers (motor, directed from the brain to the periphery), the vagus nerve is the main conduit for bidirectional communication between the brain and body organs and it participates in numerous autonomic reflexes regulating cardiovascular, respiratory, gastrointestinal, endocrine and immune functions. In the human vagus nerve, nerve fibers are arranged in fascicles, longitudinal bundles within the nerve separated by connective tissue [1, 2]. Along the cervical and thoracic vagus nerve, afferent and efferent fibers leave the fascicles and emerge from the nerve trunk to form branches, which in turn provide sensory and motor innervation to the heart, the airways and the lungs [3, 4].

Several aspects of the macroscopic anatomy of the vagus nerve have been described in detail, including perineural and epineural tissue [2], numbers and sizes of fascicles [5] and levels and patterns of emergence of organ-specific vagal branches [4, 6–9]. Despite this extensive body of work, the transverse (horizontal) and longitudinal (vertical) arrangement of fascicles in the vagal trunk, with respect to the organs to which they project and the functions they mediate, is very limited. It was recently shown that fascicles forming the superior and recurrent laryngeal branches in the swine are located on one side of the vagal trunk [5]. It is unclear whether this separate clustering of fascicles concerns other organs, in addition to the larynx, and whether it persists throughout the vagal trunk, away from the nerve levels at which organ branches emerge. Answering these questions requires tracking the trajectories of individual fascicles along many cm of nerve length. Using histological methods would necessitate staining, imaging and image analysis of thousands of nerve sections, a technically prohibitive undertaking. Microscopic 3-D imaging techniques could be used instead, e.g., microcomputed tomography (micro-CT) [10], but those have not been applied to long sections of nerves from humans or large animals.

Likewise, elements of the microscopic anatomy of the vagus nerve, most importantly the types and morphological characteristics of fibers of the vagal trunk, have been well characterized in humans and in other species [1, 6, 9, 11–23]. Fibers in the vagus nerve span a large range of sizes, from myelinated fibers with diameters >10μm, to unmyelinated fibers with diameters <0.5μm, with a variety of neurochemical phenotypes, including cholinergic, monoaminergic, glutamatergic, peptidergic etc [24, 25]. Past studies have arrived at estimates of fiber counts by analyzing spatially limited nerve areas and extrapolating to the entire nerve, assuming a uniform spatial distribution within and across nerve fascicles. However, it is unknown whether fibers are indeed uniformly distributed, or they show a preference for certain nerve areas, fascicles or even sub-fascicular sectors. Answering such questions requires identifying the majority, ideally the entirety, of many thousands of fibers in the vagal trunk, classifying them into one of several morphological types and assigning them to a location within specific fascicles. This task has not been feasible, as it requires processing of high-power microscopic images, able to resolve fibers with diameters ranging from <0.5 to >10μm, covering large nerve areas, with several stains needed to identify different morphological fiber types.

How fascicles and fibers are arranged inside the vagal trunk has implications for bioelectronic therapies based on vagus nerve stimulation (VNS). VNS relies on an electrode device implanted on the vagus nerve, that delivers electrical stimuli to alter activity of nerve fibers and modulate organ function in diseases in which the vagus nerve is implicated. VNS has been tested in heart failure [26], inflammatory bowel disease [27] and rheumatoid arthritis [28], among other diseases. However, current VNS devices deliver stimuli all around the vagus nerve, without regard to the underlying spatial arrangement of fascicles and fibers. This non-selective mode of stimulation often limits therapeutic efficacy and results in clinically significant side effects from non-targeted organs [5, 14, 29–31]. For instance, a VNS device tested in patients with heart failure, targeting vagal fibers projecting to the heart, produced side effects from the airways by activating laryngeal fibers; that prevented investigators from delivering the relatively high stimulus intensities required to activate cardioprotective fibers, thereby limiting therapeutic efficacy [32]. Fiber-selective VNS using optogenetic methods has been demonstrated in experimental animals, e.g., [33–35] [36]], however the clinical translation of optogenetic stimulation in humans is still unclear. Placement of electrodes on vagal branches to engage specific organ functions is an active area of research (e.g., [37]), however it poses significant surgical challenges. On the other hand, stimulation devices placed on the cervical vagus nerve that accommodate the anatomical of the vagal trunk could activate specific sub-populations of fibers and lead to increased efficacy with reduced side-effects [30, 38]. Such devices could expand the indications of nerve stimulation to organs not targeted by existing therapies [39, 40], and allow the mechanistic study of specific vagal reflexes in health and disease, in humans and large animals. VNS targeting specific fascicles has been demonstrated in rats [41] and in sheep [40]. However, both those species have vagus nerves with one or two fascicles, unlike the multi-fascicular human and swine vagus nerves. Therefore, the ability of fascicular VNS to deliver organ- and function-specific neuromodulation in the human vagus is unknown. Knowledge of the spatial arrangement of fascicles and fibers with regard to the organs to which they project and the functions they mediate is a prerequisite for the rational design of next generation, selective VNS devices. Currently, that knowledge is limited.

In this study, we developed a novel quantitative anatomy pipeline to characterize the transverse and longitudinal anatomical organization of the vagus, or any peripheral nerve, at several scales: branches, fascicles and single fibers. We applied the pipeline on the vagus nerve of the swine, which most closely resembles that of humans [2, 5] and discovered specific spatial organization in the vagal trunk, with fascicles and fibers grouped in distinct clusters, with respect to the physiological functions they mediate, the organs they innervate and the morphological classes of fibers. We applied the anatomy pipeline in a human cervical vagus nerve and found distinct similarities in the organization of fibers and fascicles to that of the swine. To test whether this specific fiber and fascicle arrangement can support fascicular VNS, we developed a multi-contact cuff electrode that accommodates the fascicular organization of the cervical vagus nerve and delivered fascicular VNS in anesthetized and awake swine. By targeting different sections of the nerve, we were able to evoke compound action potentials from nerve fibers of distinct functional types and elicit differential responses from organs innervated by the vagus nerve. Those electrical and physiological responses were asymmetrically distributed around the nerve in a manner consistent with the anatomical organization of the vagal trunk. These findings indicate that fascicular VNS is in principle feasible in the swine vagus nerve, a multi-fascicular nerve similar to the human vagus, and suggest that fascicular VNS should be tested further as a technique for selective VNS. At the same time, our findings have implications for defining anatomical constraints behind coding of interoceptive information in the central nervous system and the autonomic control of physiological homeostasis [42–44], and provide quantitative data for compiling realistic computational models of the vagus nerve [45]

## Results

### Fascicles form organ-specific clusters along the vagus nerve

The vagus nerve in swine is highly fascicular, with 40-50 fascicles of varying sizes in the cervical region (Suppl. Figure S1). Branches derived from the cervical and thoracic vagus nerve provide sensory and motor innervation to visceral and internal organs [4], including the larynx and pharynx, through the superior and recurrent laryngeal nerves [46], the heart, through the cardiac branch [4, 47], and the lower airways and lungs, through the bronchopulmonary branches [48]. To understand the spatial arrangement of fascicles with respect to the formation of vagal branches, we performed high resolution micro-computed tomography (micro-CT) imaging of the vagal trunk, from above the nodose ganglion, rostrally, to the lower thoracic portion of the nerve, caudally. Fascicles with any visible contribution to the formation of vagal branches were manually annotated at individual sections taken through the branching point (Suppl. Figures S2, S3, S4) and were considered to project to the corresponding organ (Figure 1A). The longitudinal trajectories of those fascicles were then tracked along the length of the vagus nerve rostrally with the use of an automated, deep learning-based, image segmentation algorithm.

**Figure 1.**
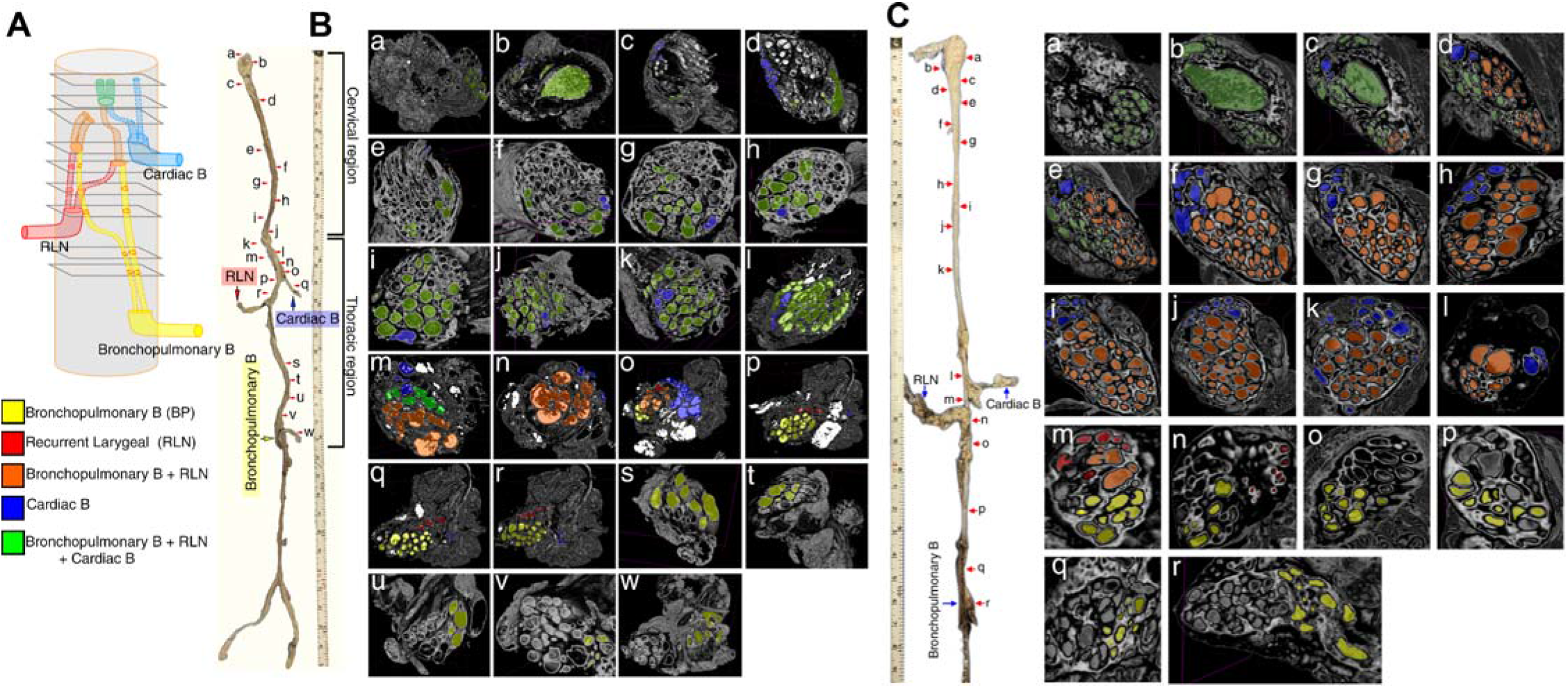
Fascicles form organ-specific clusters along the trunk of the swine vagus nerve. (A) Schematic showing a segment of the vagal trunk with 3 branches joining it (bottom to top): bronchopulmonary (BP, yellow), recurrent laryngeal nerve (RLN, red), cardiac branch (blue), (n =2). The trajectories of fascicles contributing to each branch were traced rostrally inside the trunk, even after they merged with other fascicles to form “mixed” fascicles: BP and RLN (orange), BP, RLN and cardiac (green). (B) Macroscopic structure of an intact right vagal trunk, from the nodose ganglion to the lower thoracic level (left). On the right, multiple micro-computed tomography sections through the trunk are shown (panels a-x), with fascicles contributing to each of the identified branches or combinations of branches shaded in the corresponding colors. (C) Macroscopic and fascicular structure of an intact left vagal trunk from a second animal.

Fascicles in the vagus nerve trunk contributing to the bronchopulmonary (BP) branch form a cluster close to its level of entry (yellow fascicles, Figure 1B, panels x and w; and Figure 1C, panels r and q); bronchopulmonary fascicles remain clustered for several centimeters rostrally (Figure 3B, panels v-s; Figure 3C, panels p-o). The recurrent laryngeal nerve (RNL) joins the trunk by contributing another cluster of fascicles (red fascicles; Figure 1B, panels r-q; Figure 1C, panels n-m). Some of those fascicles ascend in parallel to the BP cluster and some merge with it, in a “mixed” BP-RLN cluster (orange fascicles; Figure 1B, panels p-o; Figure 1C, panels m-l). Both BP and RLN fascicles disappear within a few centimeters and are replaced by a cluster of combined BP-RLN fascicles which runs rostrally along the upper thoracic (Figure 1B, panel n) and the cervical vagus nerve (Figure 1C, panel l). The cardiac branch joins the vagal trunk rostrally to the RLN entry and forms a separate cluster of fascicles (blue fascicles; Figure 1B, panel o; Figure 1C, panel l). Some of the cardiac fascicles maintain their separate trajectories all the way through the upper thoracic and cervical vagus nerve (Figure 1B, panels l-d; Figure 1C; panels k-d) and some merge with the mixed BP-RLN fascicles to form a larger cluster of mixed BP-RLN-cardiac (green fascicles; Figure 1B, panels m and above; Figure 1C, panels e and above). Many of the mixed BP-RLN-cardiac fascicles terminate in the nodose ganglion, and some of them bypass it (Figure 1B, panels b, a; Figure 1C, panels b, a). Fascicular branching, where one fascicle branches to form two fascicles, or fascicular merging, where two fascicles merged to form one fascicle is observed throughout the length of the nerve (Suppl. Figure S5, and Supplementary video 1: https://www.youtube.com/watch?v=gaNHijoVB_U).

**Figure 2.**
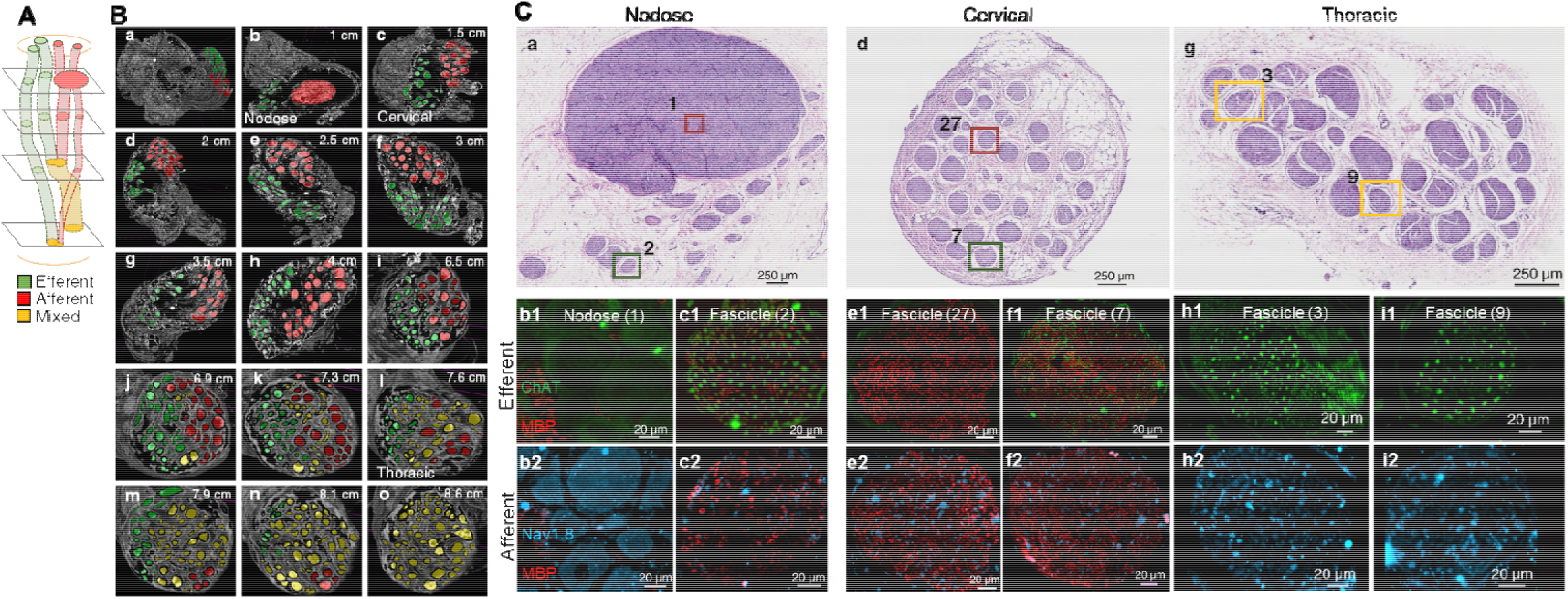
Sensory and motor fascicles form clusters along the trunk of the swine vagus nerve. (A) Schematic showing a segment of the vagal trunk with several fascicles along its path. At rostral levels, sensory fascicles (green) converge into the nodose ganglion, while motor fascicles (red) by-pass the nodose. At more caudal levels, sensory and motor fascicles merge to form mixed fascicles (yellow). (B) Micro-computed tomography imaging of an intact right vagus nerve trunk, from just above the nodose ganglion to the upper thoracic level (n=2). Shown here is results from one animal, similar results from a second nerve are shown in Suppl. Figure S6. Panels (a-i) show the trajectories of sensory and motor fascicles in consecutive sections through the cervical vagus, at different levels (shown in cm from the rostral end of the sample). (C) Imaging of afferent and efferent fibers inside sensory and motor fascicles. (a) H&E section at the level of the nodose ganglion. An area inside the sensory nodose ganglion (1) and a motor fascicle (2) are selected. (b1) Nodose ganglion stained with anti-myelin basic protein antibody (MBP, red) that labels myelin, and choline-acetyl-transferase (ChAT, green) that labels efferent, cholinergic, fibers. (b2) The same area in the nodose ganglion stained with MBP and anti-Nav1.8 antibody (cyan) that labels afferent fibers and sensory neurons. (c1) Motor fascicle (2) stained with ChAT/MBP. (c2) The same motor fascicle (2) stained with Nav1.8/MBP. (d) H&E section at the mid-cervical level. A sensory (27) and a motor fascicle (7) are selected and are stained with the same antibodies as before (e1-2 and f1-2 panels, respectively). (g) H&E section at the thoracic level. Two fascicles are selected, stained like before (h1-2 and i1-2 panels).

**Figure 3.**
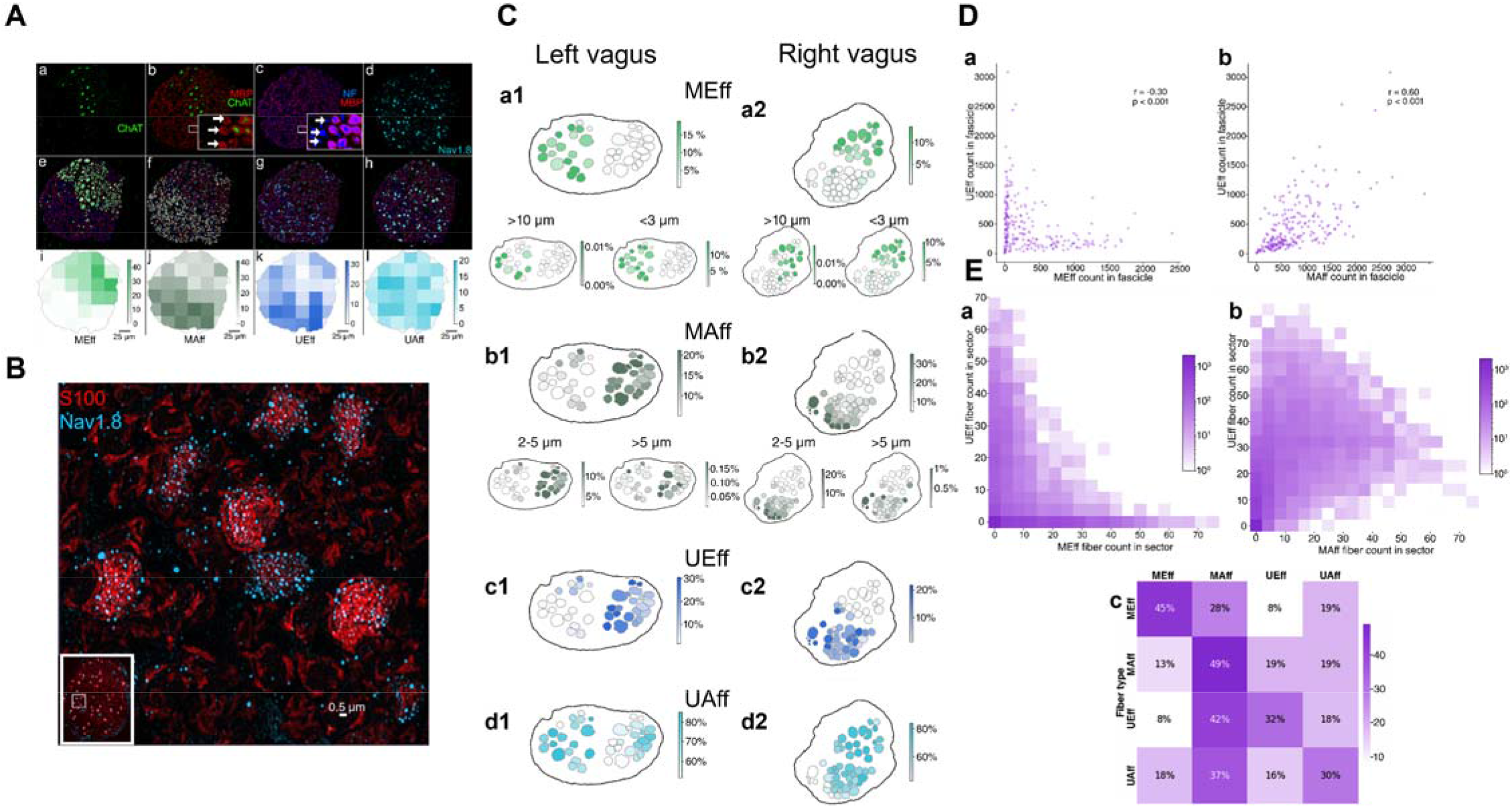
Morphologically distinct fiber types are organized across and within fascicles in the cervical vagus nerve of the swine in a specific, nonuniform pattern. (A) A large-scale immunohistochemistry (IHC) and analytical pipeline was used to image, identify, characterize and classify all single fibers, in every nerve fascicle. (a) ChAT+ fibers (Suppl. Figure S19) that were also NF+ and MBP+ (NF and MBP channels are not shown for clarity) were classified as myelinated efferents (MEff). (b) NF+ and MBP+ fibers that were ChAT-were classified as myelinated afferents (MAff); arrows point to single MAff fibers in inset. (c) NF+ and MBP-fibers were classified as unmyelinated efferents (UEff); arrows point to single MEff fibers in inset. (d) Nav1.8+ and MBP-fibers were classified as unmyelinated afferents (UAff). Unmyelinated afferent fibers did not stain for NF (Suppl. Fig. S12). (panels e-h) Individual fibers of each type were identified, and their features (e.g. size) and location in the fascicle were extracted. This allowed us to assess for different fiber types statistics of fiber counts, the area they cover and distributions across fascicles. (panels i-l) Local density values of each fiber type were extracted from fiber counts and locations, and projected on a fascicle using a color scale. This allowed us to assess within-fascicle arrangement of different fiber types. (B) IHC imaging of UAff fibers in a section of a nerve fascicle. Stains for Nav1.8, specific to afferent fibers, and S100, a marker of Schwann cells, are shown superimposed to reveal several Remak bundles, within which individual UAff fibers are surrounded by the cytoplasm of Schwann cells. Inset shows the entire nerve fascicle. (C) Counts of fibers of different types in the fascicles in a section from the left and the right cervical vagal trunk of the same animal. (a1) Counts of MEff fibers across fascicles of the left vagus nerve, as percentage of total number of fibers in each fascicle; percent counts represented by a color intensity scale. Top: Counts of all MEff fibers, independently of diameter. Bottom left: Counts of MEff fibers with diameter>10μm (Aα-type). Bottom right: Counts of MAff fibers with diameter<3μm (B-type). (a2) Same as a1, but for the right vagus nerve. (b1) Counts of MAff fibers across fascicles of the left vagus nerve. Top: for all MAff fibers; bottom left: for MAff fibers with diameters 2-5μm (Aß-type); bottom right: for MAff fibers with diameters >5μm (Aδ-type). (b2) Same as b1, but for the right vagus nerve. (c1) Counts of UEff fibers across fascicles of the left vagus, and (c2) of the right vagus nerve. (d1) Counts of UAff fibers across fascicles of the left vagus nerve, and (d2) of the right vagus nerve. (D) Distinct relationships in co-localization of different fiber types across fascicles. (a) Anti-correlated counts of MEff fibers in a fascicle (abscissa) vs. counts of UEff fibers in the same fascicle (ordinate). Each point represents a single fascicle. Shown are data from all fascicles in the cervical vagus nerve of 4 animals (8 nerves). (b) Correlated counts of MAff and UEff fibers. (E) Distinct relationships of co-localization of different fiber types within fascicles. (a) Counts of MEff and UEff fibers within a 30-μm-square fascicular sector in all analyzed fascicles. The color intensity of each pixel represents the frequency of the specific combination of MEff-UEff fiber counts: the lighter the color, the less frequent that combination is across sectors. Shown are data from all sectors within all fascicles, from all nerves. (b) Correlated counts of MAff and UEff fibers within a 30-μm-square sector. (c) Joint probability of co-localization of different fiber types within a 30-μm-square sector. In each row, color intensity and percentage values represent the probability of finding a fiber of each of the 4 types in the immediate vicinity of the fiber type represented by that row.

*These findings indicate that fascicles in the thoracic and cervical vagal trunk form spatially distinct organspecific clusters close to the levels of entry of branches to those organs; those clusters progressively merge into larger, mixed clusters in the rostral direction*.

### Fascicles form separate sensory and motor clusters along the vagus nerve

To understand the spatial arrangement of fascicles with respect to sensory or motor functions, we analyzed the micro-CT images in the opposite direction, from rostral to caudal: fascicles that originate in the nodose ganglion were considered primarily sensory (afferent) and those that bypass the nodose ganglion were considered primarily motor (efferent) (Figure 2A; Figure 2B, a, b). By tracking fascicular trajectories in the caudal direction (Suppl. Video 1: https://www.youtube.com/watch?v=gaNHijoVB_U), we found that sensory and motor fascicles form 2 distinct clusters throughout most of the cervical vagus nerve (Figure 2B, c-i); the 2 clusters begin to merge in the lower cervical region (Figure 2B, j, k), and the sensory-motor separation gradually disappears in the thoracic region (Figure 2A, l-o). Similar results from a second nerve are shown in Suppl. Figure S6.

To examine whether the separation of micro-CT-resolved sensory and motor fascicles persists when single fibers inside those fascicles are accounted for, we performed immunohistochemistry (IHC) in sections through the nodose ganglion, mid-cervical and thoracic region (Figure 2C, Suppl. Figure S7, S8, and S24). At the nodose ganglion level (Figure 2C, a), ChAT-positive (ChAT+) fibers, indicative of motor function, are almost absent (Figure 2C, b1), however Nav1.8-positive (Nav1.8+) sensory “pseudo-unipolar” cells are numerous (Figure 2C, b2; Suppl. Figure S9). In a motor fascicle, ChAT+ fibers are abundant (Figure 2C, c1); Nav1.8+ fibers are seen but are not as common (Figure 2C, c2). At the mid-cervical region (Figure 2C, d), fascicles poor in ChAT+ fibers (Figure 2C, e1) but rich in Nav1.8+ fibers (Figure 2C, e2) are separated from fascicles rich in ChAT+ (Figure 2C, f1, Suppl. Figure S10) and in Nav1.8+ fibers (Figure 2C, f2). Finally, at the thoracic region, all fascicles show intense staining for both ChAT and Nav1.8 (Figure 2C, g-i; Suppl. Figure S11).

*These findings indicate that sensory and motor fascicles form spatially distinct clusters in the upper and mid-cervical trunk of the vagus nerve; the fascicles progressively merge in the lower cervical region, and in the thoracic region all fascicles are mixed. When considered together with the organ-specific arrangement of fascicles, this suggests that physiological effects of vagus nerve stimulation will likely depend on the level at which stimulating electrodes are placed on the vagal trunk.*

### Morphologically distinct fiber types are organized across and within fascicles in a specific, nonuniform pattern

The human and swine vagus nerve contain fibers of several functional types, including myelinated and unmyelinated afferents and efferents (Suppl. Figure S28) [1, 14, 30, 47]. Myelinated efferents provide motor innervation to laryngeal muscles (Aα-type) or parasympathetic innervation to visceral organs (B-type); unmyelinated efferents are adrenergic hitchhiking postganglionic fibers from the sympathetic ganglion (C-type) [8, 47, 49], even though there is evidence for the presence of adrenergic and cholinergic afferents in the vagus nerve [50, 51]. Myelinated afferents convey sensory signals from the pharyngeal and laryngeal mucosa and laryngeal muscles (Aβ-type)[19, 30, 44, 52–54], and signals from mechanical and chemical receptors in visceral organs, including the heart [55, 56], lungs [19, 57, 58] and vessels [59] (Aδ-type); unmyelinated afferents convey chemical and possibly nociceptive signals from visceral organs (C-type) [16]. How these morphological fiber types are arranged in the vagal trunk, both across and within fascicles, is unknown. To understand the spatial organization of vagus nerve fibers of different types across fascicles in the cervical region, the most common anatomical location for placement of stimulation electrodes, we developed a quantitative IHC pipeline, consisting of IHC staining, imaging and automated fiber segmentation, feature extraction and analysis, at the single fiber level. Based on the expression pattern of several IHC markers, we classified each fiber in the cervical vagus nerve as myelinated efferent (MEff, Figure 3A, a), myelinated afferent (MAff, Figure 3A, b), unmyelinated efferent (UEff, Figure 3A, c) or unmyelinated afferent (UAff, Figure 3A, d and 3B; Suppl. figure S12). Applying additional fiber diameter criteria, we classified fiber types compatible with the Erlanger-Gasser scheme (Table I); for a more detailed discussion of criteria used in the classification of fiber types, see Suppl. Table 3, Suppl. Figure S28, and the corresponding Discussion section.

**Table I.**
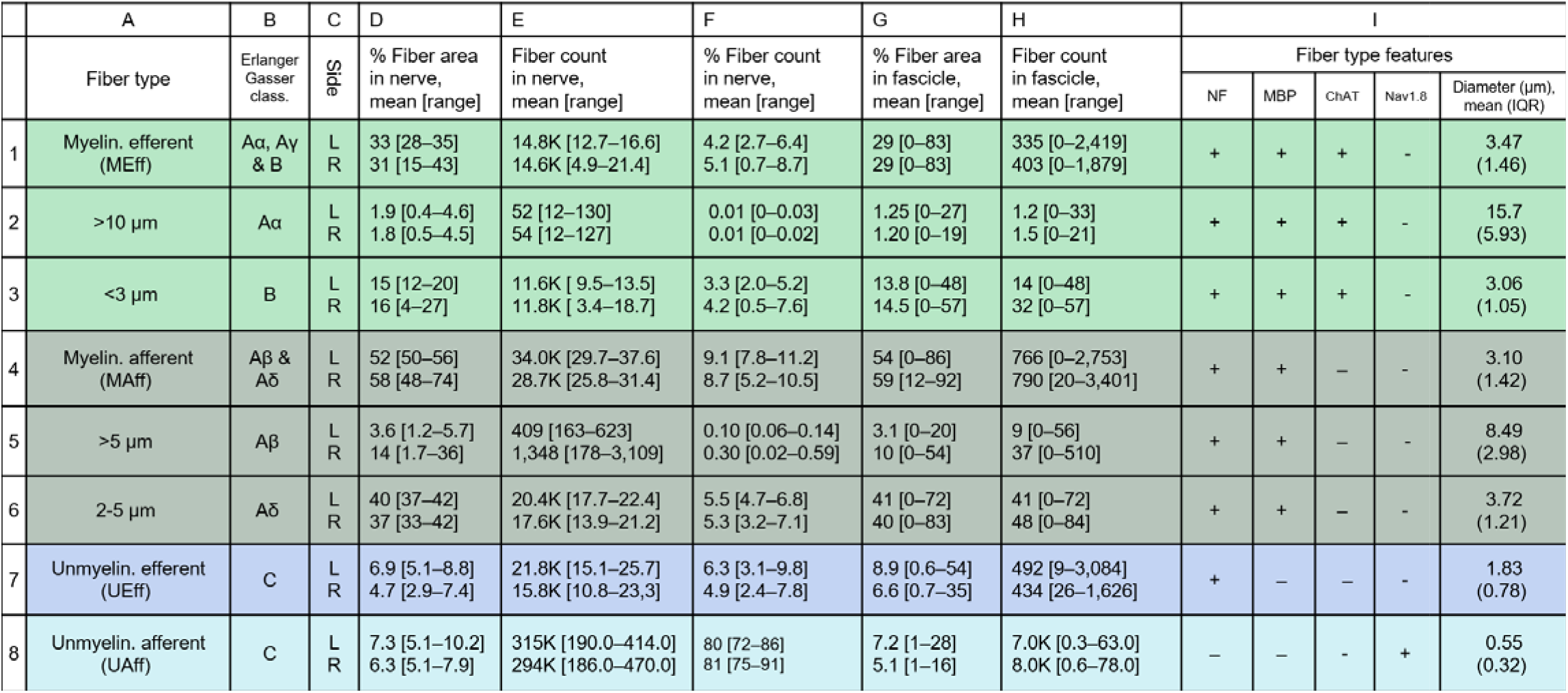
Fiber types in the cervical vagus nerve of the swine, fiber counts and areas they occupy.

Different fiber types (rows 1-8) according to criteria used in our pipeline (A) and corresponding fiber types according to Erlanger-Gasser classification (B). (C) Nerve side (left or right). (D) Percentage of fiber-covered area in the entire nerve corresponding to a given fiber type; shown is mean and range across 8 nerves. (E) Absolute fiber count of given fiber type in the entire nerve. (F) Percentage of given fiber type among all fibers in the whole nerve. (G) Percentage of area covered by given fiber type in single fascicles; mean [range] across all fascicles, of all nerves. (H) Absolute count of given fiber type in single fascicles. (I) Fiber type-specific features used in our pipeline to classify single fibers: positive (+) or negative (-) in NF stain, MBP (myelin basic protein) stain, ChAT (choline-acetyl-transferase) stain, and Nav1.8 stain. Measured diameters for each fiber type (in μm, mean and interquartile range).

Overall, we counted 200-450K individual fibers in each nerve. The 4 main fiber types have distinct fiber diameter distributions (Suppl. Fig. S13). Roughly 70-85% of them are of the UAff type; however, UAff fibers occupy less than 10% of the total fiber-covered area in the nerve (Table I). On the other end, myelinated fibers, MAff and MEff, despite representing less than 15% of total fiber count, occupy 60-80% of fiber-covered area (Table I; and Suppl. Figure S13). There is substantial variability in fiber counts and in respective occupied areas across fascicles, for all fiber types (Table I). More than 50% of all fibers in the nerve, independently of fiber type, lie within 500 μm from the epineurium (Suppl. Fig. S14A & S15). The area occupied by fibers of a specific type was not significantly different between smaller, intermediate-size and larger fascicles, and this was true for all fiber types (Suppl. Fig. S14B).

In both left and right cervical vagus, fascicles with large MEff fiber counts are found on one side of the nerve, forming an “motor cluster” (Figure 3C, a1 and a2) and fascicles with large MAff fibers are found on the opposite side, forming an “sensory cluster” (Figure 3C, b1 and b2), in agreement with our micro-CT and qualitative IHC findings (Figure 2). The very few, larger Aα-type efferents (diameter >10μm), are concentrated in specific fascicles (Figure 3C, bottom left in a1 and a2), whereas the smaller (2-5 μm in diameter), cholinergic (ChAT+) B-type fibers are more uniformly distributed within the motor cluster (Figure 3C, bottom right in a1 and a2). Importantly, the distribution of UEff fibers, a subset of which are tyrosine-hydroxylase positive (TH+) sympathetic fibers (Suppl. Fig. S16), has little overlap with that of cholinergic B-fibers as fascicles with numerous UEff fibers are commonly localized in the sensory cluster (Figure 3C, c1 and c2). Most large MAff fibers are found in specific fascicles of the sensory cluster, with some of them overlapping with fascicles with large MEff fibers (Figure 3C, bottom right in b1 and b2). Many more MAff fibers are intermediate size Aδ-type fibers, widely distributed within the sensory cluster (Figure 3C, bottom left in b1, b2). UAff fibers are present in fascicles of both sensory and motor clusters, even though fiber counts vary between 55% and over 80% (Figure 3C, d1 and d2). Maps with fiber distributions across fascicles from additional nerves are given in Suppl. Figure S17. Across all nerves, most fascicles with many UEff fibers have few MEff fibers and vice-versa (Figure 3D, a), in agreement with the non-overlapping fascicular distribution of cholinergic and non-cholinergic efferents. In contrast, many fascicles rich in UEff fibers are also rich in MAff fibers (Figure 3D, b). Interestingly, these relationships persist even at the sub-fascicular level: sectors within a fascicle with high UEff counts have low MEff counts (Figure 3E, a) and high MAff counts (Figure 3E, b). In general, within sub-fascicular sectors, fibers of the same type tend to cluster together (main diagonal of Figure 3E, c; Suppl. Figure S18).

*Taken together, these findings indicate that there are consistent, specific and highly nonuniform patterns in the way morphologically distinct fiber types are organized in the cervical vagus nerve. Afferent and efferent myelinated fibers mediating somatic and visceral sensory and motor functions are found in, largely nonoverlapping, sensory and motor fascicles. Cholinergic and noncholinergic efferent fibers are found in separate fascicles and separate sub-fascicular sectors. Unmyelinated afferent fibers are widely distributed across fascicles. These findings suggest that spatially selective, or fascicular, VNS at the cervical level could in principle be used to differentially activate distinct morphological types of vagal fibers.*

### Application of the anatomical pipeline in a vagus nerve sample from a human cadaver

The vagus nerve of humans closely resembles that of swine in size, fascicle diameter, fascicle distance from the epineurium, and branching pattern [1, 5]. The spatial arrangement of fascicles of the human vagus nerve, and their fiber composition, is unknown. To test whether the pipeline we developed can be used to characterize the anatomical organization of fascicles and fibers in the human vagus nerve, a 6-cm-long section of a vagal trunk, including the emergence of the RLN branch, was extracted from an embalmed human cadaver and was subjected to the same procedures as the swine nerves (Figure 4A, Supplementary figure S23).

**Figure 4.**
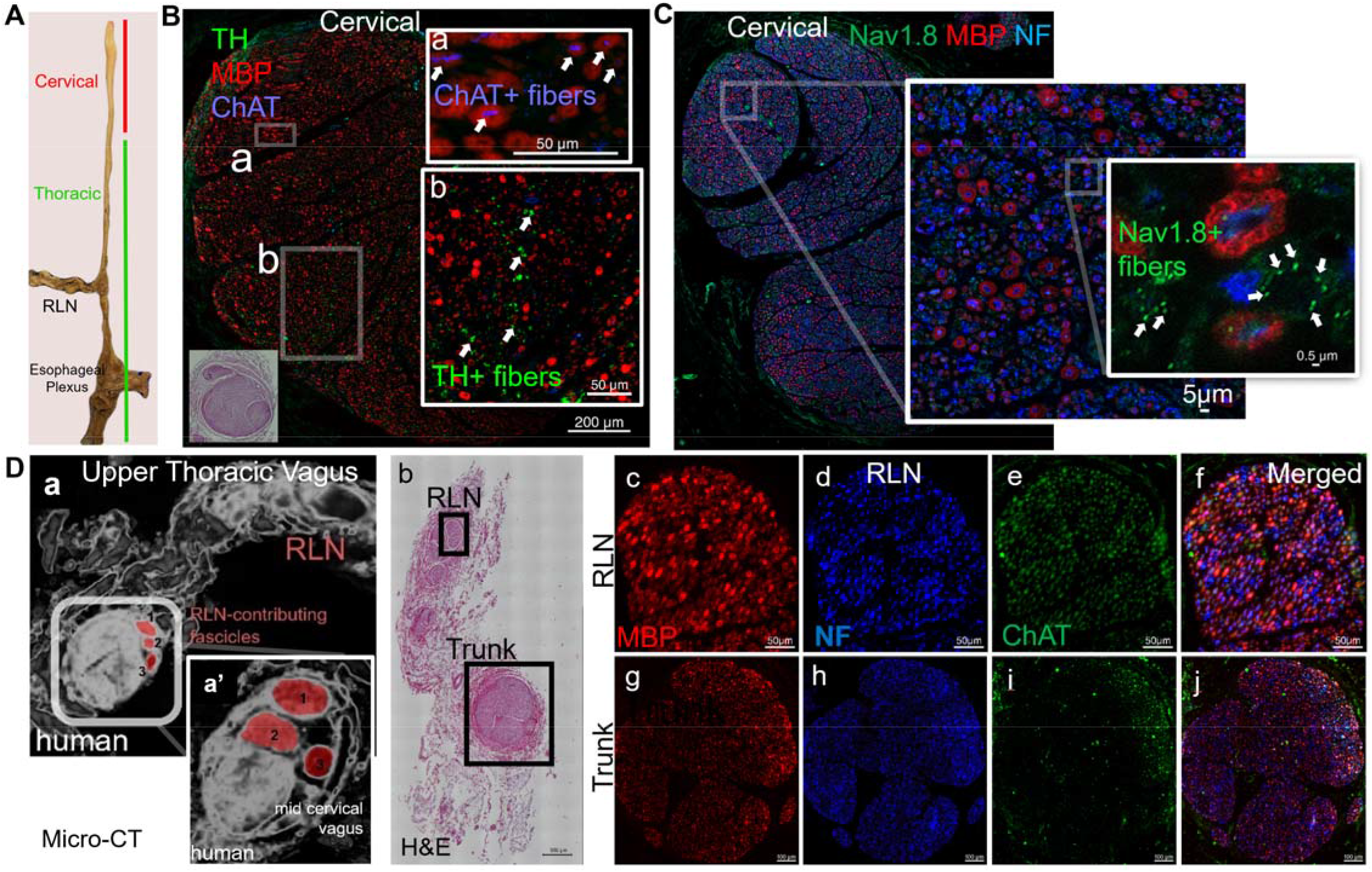
Application of the micro-CT and IHC pipelines in a human vagus nerve sample. (A) Left vagus nerve sample extracted from a human cadaver, including the recurrent laryngeal nerve (RLN) branch and part of the esophageal plexus. (B) IHC image of the sample stained with anti-tyrosine hydroxylase antibody (TH, green, catecholaminergic fibers), anti-myelin basic protein antibody (MBP, red, myelinated fibers), and anti-choline-acetyl-transferase (ChAT, green, cholinergic fibers). Insert image shows H&E image of the nerve at the same level. Panel (a), shows magnified a region of the nerve rich in ChAT+ (white arrows) but poor in TH+ fibers. Panel (b) shows a different region, rich in TH+ (white arrows), but poor in ChAT+ fibers. (C) IHC image of the same sample stained with anti-Nav1.8 antibody (green, unmyelinated afferents, UAff), MBP (red) and NF (blue). A selected area is shown at 2 levels of magnification. Highest power image shows super-resolution aryscan confocal image of individual UAff fibers (white arrows). Notice the lack of Nav1.8-NF co-localization, similar to the swine vagus nerve. (D) (a) Micro-CT image at the level of emergence of the recurrent laryngeal nerve (RLN, red fascicles) at the upper thoracic region. Inset panel (a’), shows a second micro-CT image of the nerve taken at the mid cervical level, where fascicles contributing to the RLN branch where traced. (b) H&E image from the level of RLN emergence. Two fascicles are selected for IHC imaging, one in the RLN branch and one in the main trunk. Panels (c-f), and (g – j) show the 2 fascicles stained with MBP, NF and ChAT.

A segment from the mid-cervical region of the extracted nerve was co-stained for either TH, MBP and ChAT or Nav1.8, MBP and NF (Figure 4B, C). Areas rich in ChAT+ fibers are poor in TH+ fibers (Figure 4B, a); conversely, areas rich in TH+ fibers are poor in ChAT+ fibers (Figure 4B, b). However, Nav1.8+ fibers are found to be distributed ubiquitously throughout the nerve with no spatial preference (Figure 4C). These findings are consistent with the distribution pattern of different fiber types observe in the swine vagus nerve (Figure 3). The same nerve sample was subjected to micro-CT scanning; fascicles giving rise to the RLN branch were manually identified at the level of branching (Figure 4D) and were traced to the mid-cervical region (Supplementary video 2: https://youtu.be/bVRJQn-3Fe4). We found that RLN fascicles are separately clustered from other fascicles, starting at the branching point and for several cm in the cephalad direction (Figure 4Da and a’), similar to the organization of RLN fascicles in the swine vagus nerve (Figure 1). IHC imaging of the RLN branch and the vagal trunk reveals arrangement of fiber types similar to those of the swine vagus nerve: the RLN fascicle is rich in large, myelinated ChAT+ fibers (Figure 4D, c-f), in agreement with a recent characterization of the emergence of the RLN from the vagal trunk in swine [5], whereas the fascicle in the vagal trunk is poor in ChAT+ fibers (Figure 4D, i) and shows non-uniform distribution of other, presumably afferent, large myelinated fibers (Figure 4D, g, h, j), similar to our findings in the swine vagal trunk (Figure 3). These results suggest that the anatomical pipeline used to resolve the microscopic structure of the swine vagus nerve at the fascicle and single fiber level, could also be used in the human vagus nerve as well; they also provide preliminary evidence for organ- and function-specific organization of fibers in the human vagus nerve, even though additional studies are needed.

### Fascicular vagus nerve stimulation asymmetrically elicits function- and organ-specific electrical and physiological responses

Activation by bioelectronic devices of different populations of efferent and afferent fibers innervating peripheral organs determines the effects of vagus neuromodulation therapies [30, 60, 61]. Having demonstrated that the vagal trunk has a specific, nonuniform organ- and function-specific anatomical organization, we next sought to test whether spatially selective, fascicular VNS can differentially activate fibers innervating specific organs, as well as fibers for controlling sensory or motor functions. Intraneural electrodes are likely best suited for fascicle-targeting neurostimulation, but they are not well characterized with respect to their elicited stimulation effects [62] and may be associated with an increased risk of nerve injury [63]. Given that >50% of fibers lie within 500 μm from the epineurium (Suppl. Figure S14), we developed a multi-contact cuff electrode device to deliver spatially selective nerve stimulation from the nerve’s surface. Based on the spatial overlap between nerve fascicles and circular representations of electrical fields generated by epineural contacts (Figure 5A, a), an 8-contact configuration was chosen because it attained higher overall efficacy compared to configurations with fewer contacts, with the same selectivity as those with more contacts (Figure 5A, b). The cuff has a helical design to accommodate nerves of different external diameters and includes 8 small, square-shaped contacts, made of platinum-iridium (Pt-Ir), evenly distributed around the circumference and 2 ring-shaped return contacts (Figure 5C). The electrodes were characterized after repeated use to determine durability, using electrochemical impedance spectroscopy and cyclic voltammetry (Figure 5D); no change in electrode impedance was found between the 1st and 6th use (Figure 5E). Charge storage capacity (CSC) measurements performed at a scan rate of 100 mV/s over the water window (after the 6th use) yielded 3.9 ± 0.63 mC/cm2, which is comparable to previously reported values for Pt-Ir electrodes [64, 65].

**Figure 5.**
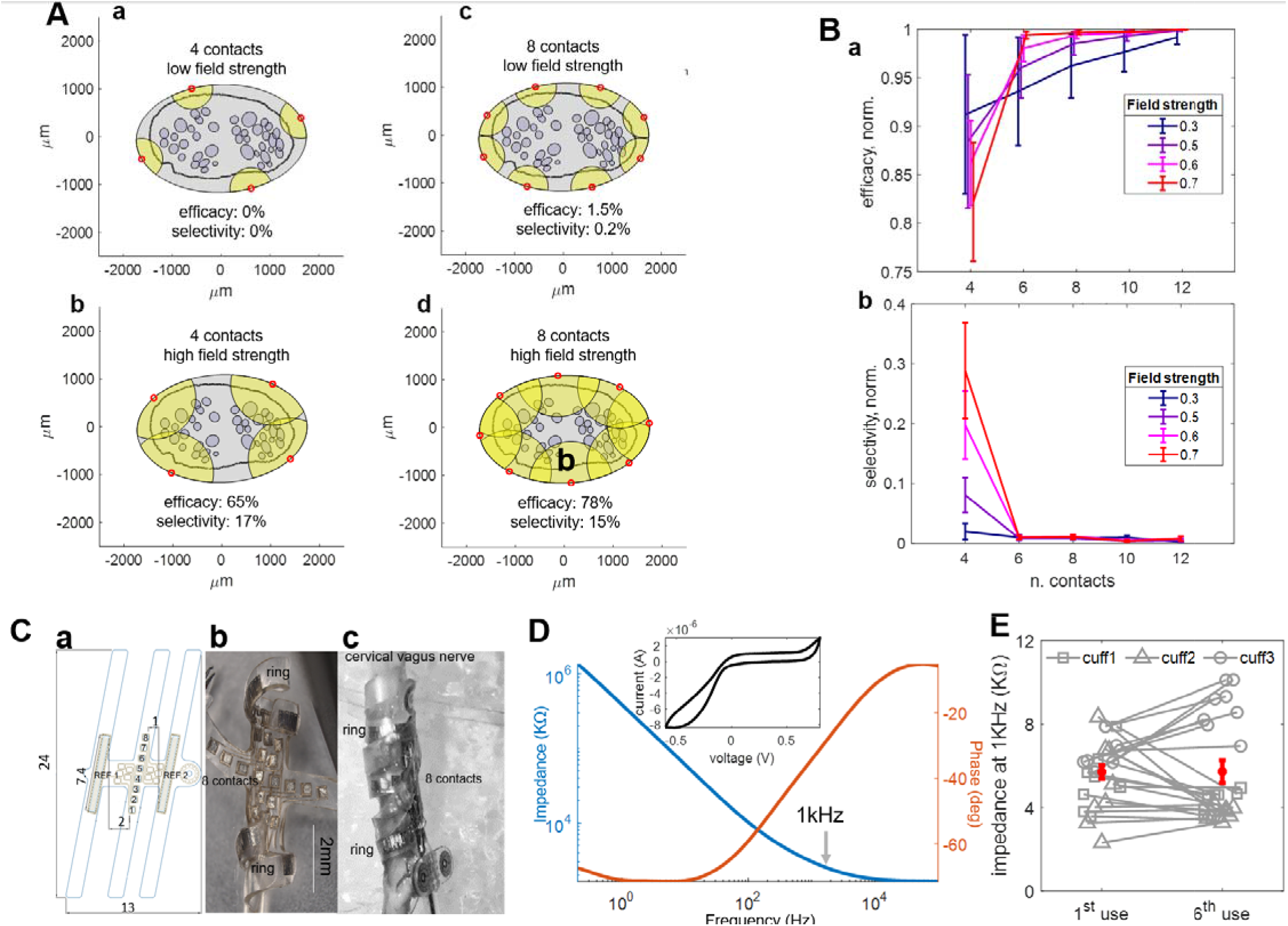
Development of a multi-contact cuff electrode for fascicular stimulation of the cervical vagus nerve. (A) Methodology to estimate efficacy and selectivity of different contact arrangements in a multi-contact cuff electrode. The outline of a swine cervical vagus nerve and of its fascicles is shown surrounded by an ellipse representing the cuff; red dots represent point-like contacts, and yellow circles around contacts represent electrical fields that activate all fibers in the fascicles that overlap with those circles. Efficacy of a given configuration is defined as the percentage of nerve fascicles activated by all contacts; selectivity as the percentage of fascicles activated, on average, by any one contact. Shown are examples of efficacy and selectivity for a 4-contact configuration, at low (a) and high field strength (b). Efficacy and selectivity for an 8-contact configuration (c, d). (B) Normalized selectivity and efficacy as a function of number of contacts in the cuff electrode design, for 4 different field strengths; means (+/-SEM) calculated across 8 nerves and 100 random orientations of each electrode configuration around each nerve. (C) (a) The final design of the helical cuff electrode. The cuff is comprised of 3 silicone loops that wrap around the nerve. The middle loop houses 8 small square contacts made of platinum iridium (500μm side, 500 μm apart from each other) and each of the 2 outer loops houses 1 ring-shaped return contact (500 x 7400 μm). (b) Close up of the actual helix cuff. (c) Cuff placed on a swine cervical vagus nerve. (D) Electrochemical impedance spectroscopy shows the impedance magnitude and phase of a representative electrode across a measurement frequency range from 0.2Hz to 100KHz; impedance measured at 1kHz is ~4 kΩ. The top inset shows a representative cyclic voltammetry trace used to characterize charge storage capacity (~4 mC/cm2), scan range from −0.6 to 0.8V at rate of 100 mV/s. (E) Individual impedance values at 1kHz (mean+/-SEM), for each of the 8 square contacts in 3 cuff electrodes, measured after the first and sixth use in acute in vivo VNS experiments.

The multi-contact cuff electrode was used in acute stimulation experiments in anesthetized swine. Stimulation through different contacts in the multi-contact electrode produces different physiological responses specific to organs receiving vagal innervation: changes in heart rate and in blood pressure (heart and vessels), changes in breathing rate (lungs), and contraction of laryngeal muscles (Figure 6A, a and b). Stimulation elicits evoked compound action potentials (eCAPs), recorded through a second cuff electrode placed at a distance from the stimulating device, reflecting direct activation of different fibers. Depending on which contact is used for stimulation, differential activation of fast fibers, with a conduction velocity compatible with that of Aα/β-type fibers, and of slow fibers, compatible with Aδ/B-type fibers, is observed (Figure 6A, c and d). Administration of a neuromuscular blocking agent (vecuronium), abolishes EMG activity recorded from laryngeal muscle (Figure 6A,f) but does not alter eCAPs from the vagus nerve (Figure 6A, e), confirming that no EMG activity contaminates eCAP recordings. Graded physiological organ responses and, accordingly, eCAP amplitudes of fibers mediating those responses are observed at a range of stimulus intensities, resulting in recruitment curves that are asymmetric across contacts (Figure 6B). Similar recruitment maps were also seen in another series of tests (Suppl. Figure S21).

**Figure 6.**
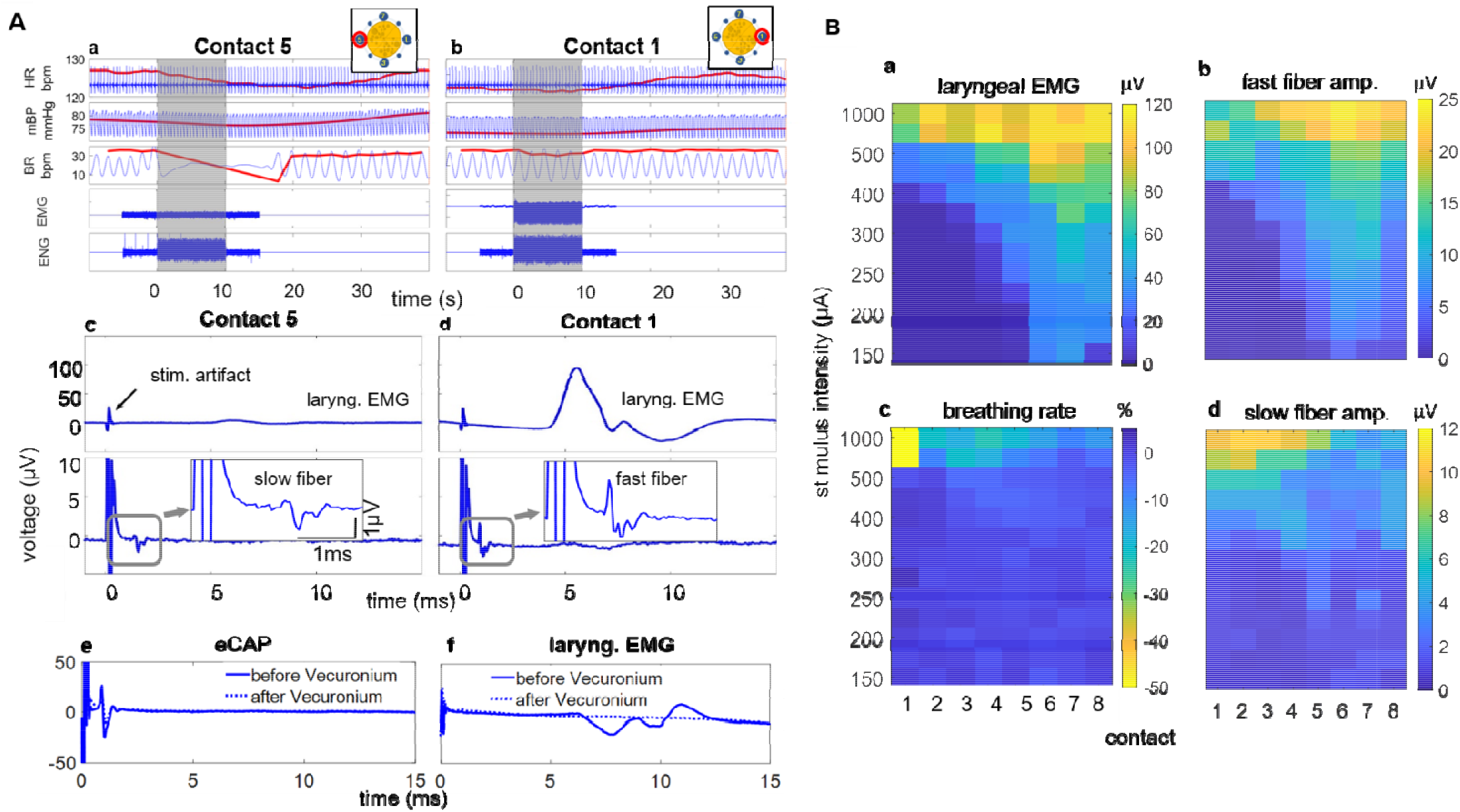
Physiological and evoked nerve potential responses to fascicular vagus nerve stimulation. (A) (a) Examples of cardiovascular, respiratory and laryngeal physiological responses to a stimulus train (shaded area) through one of the 8 contacts (contact 5) include a drop in heart rate (HR), a moderate drop in mean blood pressure (BP) and slowing of breathing rate (BR); at the same time, minimal evoked laryngeal muscle activity (EMG) and strong evoked responses in the nerve potential (ENG) were observed. (b) Example responses to the same stimulus train delivered through a different contact (contact 1) included minimal drop in HR, BP and BR but strong laryngeal EMG. The red line indicates the mean cycle rate from the cycling raw data. The two sketches in (a) and (b) indicate the relative position of the tested contacts around the nerve. (c) Evoked laryngeal EMG (top) and compound nerve action potential (eCAP, bottom) triggered by stimuli in the train delivered to contact 5. Small EMG is generated and the eCAP shows a response at a latency consistent with slow A-fiber activation. (d) EMG and eCAP triggered by stimuli delivered to contact 1; EMG response and a short latency eCAP, consistent with fast A-fiber activation, are observed. (e) nerve eCAP is not affected by neuromuscular blocker vecuronium. (f) EMG of laryngeal muscle is blocked by neuromuscular blocker. (B) Example physiological responses and eCAPs mediating them have different recruitment curves from different contacts. (a) Amplitude of laryngeal EMG is shown for different stimulation contacts (abscissa) and at different intensities (ordinate), represented by the color scale. (b) Same for amplitude of fast fiber response in the eCAP; fast fibers (e.g. Aα) mediate efferent signaling to laryngeal muscles. (c) Same for the magnitude of breathing response, quantified as % changein BR. (d) Same for amplitude of slow fiber response in the eCAP; slow fibers (e.g., Aδ) mediate afferent signaling to the brain that, through a reflex mechanism (Herring-Breuer reflex) slows down or arrests breathing.

To quantify and visualize the asymmetry in stimulation effects from different contacts and compare the radial organization of physiological and eCAP responses across animals, threshold intensities for eliciting organ responses [66] were determined for each contact and were plotted on a polar map representing the “directionality” of responses around the nerve (Figure 7A). Organ-specific physiological responses have different thresholds, and those thresholds depend on which contact is used for stimulation, resulting in asymmetric maps; also, maps have different shapes, depending on which organ’s response is considered (Figure 7A). In each animal, the radial location of the contact associated with the lowest heart rate threshold (heart-specific location) was arbitrarily placed at 12 o’clock (90° angle) in the polar plot. After aligning polar plots in each animal by that contact and normalizing responses between a minimum and maximum value, we then compiled average maps of the directionality of organ responses across animals (Figure 7B). The vector sum of average heart rate thresholds points towards the location with the greatest threshold and away from the “heart-specific” location, which lies at an angle of 134° (Figure 7B, a). The lung-specific location lies at an angle of 155° (Figure 7B, b) and the larynx-specific location lies at an angle of 65° (Figure 7B, c). The directionality map for the blood pressure response is similar to that for heart rate (Figure 7B, d). Stimulus-elicited CAPs also depend on the stimulated contact, which reflects in differential recruitment of fast and slow fibers (Figure 7C). After aligning data in individual animals by the contact with the maximum fast fiber amplitude (placed at 90° angle) and normalizing fiber amplitudes, we compiled maps of the directionality of fast and slow fiber responses in several animals. The fast fiber-specific location lies at 90° (Figure 7D, a), and the slow fiber-specific location at 154° (Figure 7D, b).

**Figure 7.**
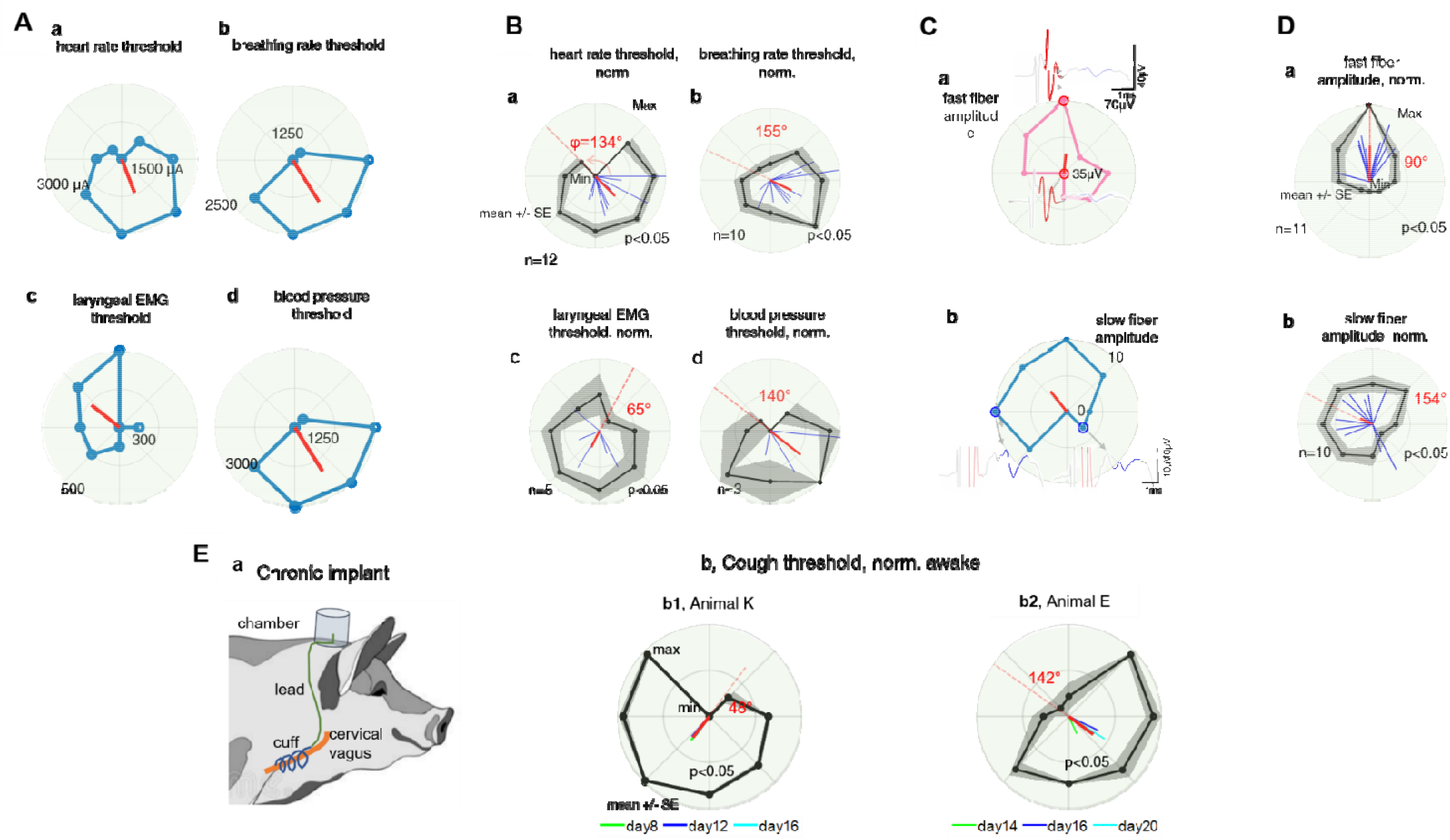
Fascicular vagus nerve stimulation differentially produces organ-specific physiological responses and fiber-specific evoked nerve potentials. (A) Example in a single animal of threshold intensities for different physiological responses, as determined for each of the 8 contacts in the multi-contact cuff electrode. (a) threshold for HR response, (b) BR threshold, (c) laryngeal EMG threshold, (d) BP threshold. Threshold was defined as the minimum intensity of a stimulus train (200 μs pulse width, 30 Hz) that produces a change of 5-10% in the corresponding physiological variable. The contact with minimum HR threshold is placed at 12 o’clock direction (origin of polar plot); the remaining plots are aligned to that orientation. In each polar plot, the origin represents the minimum threshold (e.g. 1500 μA for HR threshold, 1250 μA for BR threshold), and the outer circle the maximum threshold (e.g. 3000 μA for HR threshold, 2500 μA for BR threshold). Each point in the polar plots is the average determined from 3-5 stimulus trains delivered to each contact. The red line represents the resultant (vector sum) of thresholds across all eight contacts: its direction points towards the side of the nerve with the highest thresholds overall, i.e. away from the side that is most selective for that physiological response. (B) (a) Average (+/-SEM) normalized HR thresholds and resultant vectors in 12 individual animals. Each blue vector represents the resultant of HR thresholds around the nerve in a single animal. The red vector represents the sum of all individual resultant vectors, and it points towards the side of the nerve with the largest heart rate threshold; the dashed line opposite to that, and the associated angle, represents the most cardiac selective direction. p-value represents statistical significance of 1-way ANOVA (dependent variable: threshold, independent variable: contact angle). (b) Average BR thresholds and resultant vectors from 10 animals. (c) Average laryngeal EMG thresholds and resultant vectors from 5 animals. (d) Average blood pressure thresholds and resultant vectors from 3 animals. (C) Example in a single animal of eCAP responses and fiber amplitudes, elicited from different contacts. (a) Traces of eCAPs elicited by stimulation delivered to 2 different contacts located at the 12 and 6 o’clock directions, with the fast (red) fiber components highlighted. Polar plot of the amplitudes of fast fiber responses, ranging between minimum (origin) and maximum (outer circle). The contact eliciting the maximum fast fiber response is placed at 12 o’clock direction. The red vector represents the resultant vector of fast fiber amplitudes from all 8 contacts. (b) Same as (a), but for amplitudes of slow fiber responses. The 2 polar plots are aligned by the 12 o’clock contact. (D) (a) Average (+/-SEM) normalized eCAP response amplitudes for fast fibers (shaded trace) and resultant vectors (blue vectors) from 12 individual animals. Before averaging, individual polar plots were aligned to the contact associated with the maximum fast fiber response. The red vector represents the sum of all individual resultant vectors. p value represents statistical significance of 1-way ANOVA (dependent variable: fiber response amplitude, independent variable: contact angle). (b) Same as (a), but for slow fibers. Plot is aligned by the contact associated with the maximum fast fiber response (12 o’clock in panel (a)). (E) Physiological responses elicited by fascicular VNS delivered for up to 3 weeks in chronically implanted swine. (a) Schematic of the chronic vagus nerve implant in swine, involving the helix cuff. (b) Cough-reflex thresholds in two animals. (b1) Average (+/-SEM) normalized cough-reflex thresholds and the resultant vectors at 8, 12, and 16 days post-implantation (shown in respective colors) in animal K. Red vector represents the sum of all days, pointing towards the side of the nerve with the largest cough-reflex threshold; the dashed line opposite to that, and the associated angle, represents the most coughing selective direction. (b2) Same as (b1), but for Animal E.

To test whether selectivity of nerve stimulation remains consistent across days, we performed chronic vagus nerve implants with the multi-contact cuff electrode device in 2 animals (Figure 7E). By stimulating through different contacts during awake experimental sessions, we found that threshold intensities for eliciting the cough reflex, whose sensory arc is mediated by myelinated afferents innervating the laryngeal and pharyngeal mucosa [19, 44, 53, 54], are asymmetrically distributed around the nerve and, despite the gradual increase in thresholds, the overall shape of the spatial distribution is maintained for up to 3 weeks post-implantation (Figure 7E, b). The chronic implants were associated with minimal damage to the nerve fibers (Suppl. Figure S25).

*These findings indicate that the highly nonuniform spatial organization of fascicles and fibers in the vagus nerve can in principle be leveraged with a multi-contact cuff electrode to elicit differential nerve responses from vagus-innervated organs and functional fiber types*.

## Discussion

In this study, we demonstrate rich, nonuniform structure in the anatomical organization of the cervical and thoracic vagus nerve of the swine, with specific organ- and function-specific arrangement of fascicles and fibers in the transverse and longitudinal directions. We also show that a multi-contact cuff electrode device that delivers fascicular vagus nerve stimulation (VNS), can in principle differentially activate vagal fibers of different morphological types that are mediating different organ-specific functions. These findings have implications for the development and testing of selective vagus neuromodulation therapies targeting specific organs and functions mediated by the cervical vagus nerve in humans.

### Choice of swine as an animal model of the human vagus nerve

In our study, we chose the swine as animal model because the swine vagus nerve more closely resembles the human in its macroscopic fascicular structure, effective diameter and relative thickness of the endoneurium, perineurium and epineurium, compared to rodents and non-human primates [2, 5]. These macroscopic features are particularly relevant to the translational testing of neurostimulation therapies based on implanted devices [30]. In a proof-of-principle study of a single human vagus nerve sample, we document additional similarities between the swine and the human vagus nerves, that extend to nerve fibers themselves and their spatial arrangement in the nerve trunk. For example, we found for the first time that myelinated efferent (MEff), ChAT(+), fibers and many of the unmyelinated efferent (UEff), TH(+), fibers occupy separate fascicles and areas within the human vagus nerve (Figure 4A), similar to the distribution of MEff and UEff fibers in the swine (Figure 3C, a vs. c). We found that unmyelinated afferent (UAff), Nav1.8+, fibers are widely distributed throughout fascicles in the human vagus nerve (Figure 4C), similarly to their distribution in the swine (Figure 3C, d). Finally, we describe for the first time that fascicles contributing to the recurrent laryngeal branch in humans are clustered together in the nerve trunk several cm rostral to the emergence of the branch (Figure 4D), similarly to the organ-specific clustering of fascicles in the swine (Figure 1B, 1C). A major difference between the human and the swine vagus nerve is the number of fascicles in the cervical vagus nerve: the swine vagus nerve has 5-10 times more fascicles than the human vagus nerve and accordingly smaller fascicle diameters (Suppl. Figure 1), offering a more “granular” anatomical substrate for testing methods and technologies for selective vagus neuromodulation [1, 30, 39, 67, 68].

### Anatomical organization of fascicles in the vagal trunk

A first level of organization of the vagus nerve concerns the organs that are innervated by branches emerging along its course: cervical branches, including the superior and recurrent laryngeal nerves for the larynx and pharynx [5, 7], thoracic branches, including the cardiac and bronchopulmonary nerves for the heart and the respiratory system, and numerous lower thoracic and abdominal branches, innervating organs of the gastrointestinal system [1]. By imaging fascicles contributing to individual organ-specific branches with micro-CT [3] and tracking the 3D trajectories of individual fascicles along the nerve trunk, we discovered that organ-specific fascicles form clusters close to the level at which those branches emerge and progressively merge with clusters from other organs, at more rostral levels (Figure 1). Cardiac fascicles are an exception to this rule, as some of them do not merge with other fascicles (Figure 1), an attribute that may facilitate selective VNS for cardiac diseases [69]. Recently, it was reported that fascicles contributing to the formation of laryngeal nerves in swine are grouped separately from other, non-laryngeal, fascicles in the vagal trunk [5], consistent with our findings in swine (Figure 1) and in human vagus nerves (Figure 4). We suggest that this separation is a manifestation of an organotopic anatomical organization of the vagal trunk, in which clusters of fascicles form organ-specific branches, at least along parts of the thoracic and cervical regions. The presence of an organotopic organization of fibers in the trunk of the vagus nerve is reminiscent of the topographic organization of motor vagal neurons in the dorsal motor nucleus of the brainstem [70, 71] and of sensory vagal neurons in the nodose ganglion [71] and the nucleus tractus solitarius [72]. The progressive merging of organ-specific fascicles in the cephalad direction suggests that fascicles of the cervical vagus nerve are organized in a “fanning-out” manner with respect to their organ connectivity (Figure 1A).

A second level of organization of the vagus nerve concerns the sensory and motor innervation of organs, provided by afferent and efferent fibers, respectively. By tracking the 3D trajectories of fascicles emerging from the sensory nodose ganglion (primarily sensory fascicles) and of fascicles by-passing the nodose ganglion (primarily motor fascicles), we document for the first time spatial separation between the 2 groups, that persists throughout most of the cervical region; fascicles forming the motor and the sensory clusters start merging in the lower cervical and are completely merged in the upper thoracic region (Figure 2B; Suppl. Figure S6). This sensory-motor separation involves large myelinated afferent and efferent fibers that provide somatic sensory and motor innervation to the larynx and pharynx (Figure 3C, a and b), in agreement with a recent finding that motor laryngeal fascicles from a cluster separate from the nodose ganglion [5]. This suggests that fascicular VNS could in principle suppress the off-target effects of laryngeal muscle contraction, by avoiding the motor cluster, or of coughing reflex, by avoiding the sensory cluster. We additionally find for the first time that this separation includes smaller afferent and efferent fibers that provide visceral sensory and motor innervation to visceral organs (Figure 3C, panels a, fibers <3μm, panel b, fibers 2-5μm, and panels c, d). Therefore, the afferent-efferent separation of fascicles may represent an organizing principle for both somatic and visceral fibers in the vagus nerve throughout most of the cervical region, similar to the distinct topographic arrangement of some somatic peripheral nerves [6]. This finding suggests that sensory-motor functions in the cervical vagus nerve are organized, at a large scale, in a “cord-like” manner: the 2 functions are mediated by separate fascicles that descend in parallel to each other, similar to cords running down a cable, only to merge at the lower cervical and upper thoracic vagal trunk (Figure 2A).

Taken together, the increasing separation of sensory-motor fascicles in the cephalad direction and of organ-related fascicles in the caudad direction, suggests that the level of the cervical vagus nerve at which an electrode is implanted is likely to influence the organ- and function-related selectivity of VNS. Placing an electrode close to the emergence of organ-specific branches, e.g., on the lower cervical and thoracic vagal trunk, is likely to maximize selectivity related to organ functions; in contrast, placing an electrode closer to the nodose ganglion, e.g., on the mid and upper cervical vagal trunk, is likely to maximize sensory vs. motor selectivity. For those reasons, mid-cervical placement of a stimulation device may be the preferred option for selective VNS, given the ease of surgical access of that location.

### Anatomical organization of morphological fiber types across and within fascicles

A third layer of neural organization in the vagus nerve concerns the several morphologically-distinct fiber types that comprise it [73]. Historically, to obtain high resolution mapping of fibers in the vagus nerve, electron microscopy (EM) imaging has been used to visualize, classify and quantify single fibers based on their ultrastructural characteristics. To discriminate between afferent and efferent fibers, selective vagotomies are performed at different levels, causing selective loss of injured afferent or efferent axons, whose distributions are then compared to those in intact nerves. In those studies, fiber quantification is performed by manual annotation of EM images from randomly selected regions of the nerve; the number of fibers in the sampled area is then extrapolated to estimate the total number of fibers in the entire nerve [20]. This procedure assumes uniform distribution of fibers across and within fascicles, something that our data does not support. Because it involves EM imaging and selective vagotomies, this process is time-consuming and expensive and cannot realistically be used to fully resolve fiber arrangement in many fascicles and/or in many nerves.

To address these issues, we developed a quantitative pipeline for staining, imaging and automated annotation and quantification of single fibers. In our study we used an IHC panel of antibodies against choline acetyltransferase (ChAT) (Suppl. Figure S19), neurofilament (NF) and myelin basic protein (MBP), and the colocalized expression of all three stains together was used to identify large myelinated efferent (MEff) fibers (Figure 3A). Fibers that lack ChAT staining but are positive for NF and MBP were classified as myelinated afferents (MAff) and fibers that lack ChAT and MBP are classified as unmyelinated efferents (MEff). In our study we imaged the entire nerve and all fascicles at a resolution of 0.150 microns/pixel. At that resolution, individual axons as small as 0.3 μm diameter can be visualized and annotated. To the best of our knowledge, this is the first methodology using light microscopy that allows complete characterization of all the main morphological fiber types in a peripheral nerve, including small, unmyelinated fibers. ChAT+ fibers in previous reports were quantified using a DAB chromogen technique, at a relatively low power e.g. 20×, which does not allow accurate estimation of fiber diameter [5]. In this work, we employed immunofluorescence (IF) to detect ChAT expression. The advantage of the IF technique is the ability to colocalize the ChAT expression with NF and MBP. Also, IF enabled us to visualize individual fibers at 100× magnification, accessing the subcellular localization of the ChAT, NF, and MBP in the same axon with high selectivity and sensitivity. Small afferent fibers do not express NF in our stains (Suppl. Figure S12) and therefore could not be identified using that marker. We instead relied on an antibody against Nav1.8, a tetrodotoxin-resistant voltage-gated channel encoded by the SCN10A gene specifically expressed by small diameter sensory neurons in spinal and cranial nerves [74, 75], to identify clusters of unmyelinated afferents (Figure 3B; Suppl. Figure S11; Suppl. Figure S20). Since non-myelinating Schwann cells in Remak bundles are known to encompass small unmyelinated fiber clusters [13, 76], we confirmed detection of C-afferent clusters by testing for co-localization with the S100 antibody expressed by Schwann cells (Figure 3B). In our study, Nav1.8+ fibers do not stain for MBP or NF (Suppl. Figure S12) which is consistent with their expression in unmyelinated, nociceptor fibers but not myelinated, mechanoreceptor afferent fibers in the vagus nerve [75]. Sizes and counts of Nav1.8+ fibers in our study are consistent with those of small, unmyelinated afferents in EM studies of the cervical vagus nerve in humans [12]. However, additional investigations directly comparing IHC to EM imaging will be needed for unequivocal confirmation that Nav1.8 is an accurate and precise marker of UAff fibers in the vagus nerve.

Until now, analysis of IHC data in nerves to segment single fibers has relied on inherently slow manual or semi-automated processes, e.g. [77]. Likewise, analysis of micro-CT data in nerves has used software-aided manual segmentation methods [10, 78]. In this study, we used a mask-RCNN deep-learning architecture to segment both micro-CT and IHC images [79, 80]. Deep-learning based algorithms have recently received attention in anatomically guided, medical image segmentation [81]. With regard to segmentation of anatomical features of nerves, deep-learning algorithms have been used on EM images [82], to resolve single fibers, and on ultrasound [83] and histological images [84], to resolve fascicles. To the best of our knowledge, this is the first use of convolutional neural networks on micro-CT and IHC data, to automatically segment and extract anatomical features from micro-CT or IHC data. After additional validation, similar pipelines to the ones described here may be useful in quantitative anatomical studies of peripheral nerves in health and disease, by improving efficiency and by minimizing sampling errors [85].

Using the pipeline, for the first time, we were able to count, rather than estimate, the entirety of fibers in several vagus nerves (4 left, 4 right), at the cervical level, and provide their detailed spatial distributions across all fascicles. The fiber counts we documented in the swine vagus nerve are comparable to those reported in the, much less fascicular, ferret [11] and cat nerves [20]: MEff ~4% of all fibers in swine (4% in ferret), MAff ~8% (9% in cats and ferrets), UEff ~5% (8% in ferrets, 26% in cats), and UAff >80% (79% in ferrets, 54% in cats). Ours is the first quantification of the distribution of UEff fibers in the swine vagus nerve. Many of the UEff fibers are small, TH+, adrenergic fibers (Suppl. Figure S16). The remaining UEff fibers (TH-) stain for NF but not for Nav1.8, consistent non-adrenergic non-cholinergic (NANC) efferents; for example, the vagus nerve contains nitrinergic [86, 87] and peptidergic efferent fibers [87–89]. Our study is also the first to describe the complete arrangement of vagal UAff fibers across the entire vagus—or any large size nerve—using the Nav1.8 marker, a product of the SCN10A gene, which is highly expressed in sensory neurons in the nodose ganglion [90]. UAff fibers occupy 5-10% of the nerve area on average and 1-28% of area in fascicles (Table I); accordingly, a recent EM study in the human cervical vagus nerve estimated that UAff fibers occupy up to 18% of the nerve area [13]. The fascicular distribution of UAff fibers in the vagus nerve is described here for the first time. We find significant numbers of UAff fibers in every fascicle; this means that even motor fascicles have significant numbers of UAff fibers; given that motor fascicles bypass the nodose ganglion, a percentage of identified UAff fibers may represent peripheral axons of sensory cells outside of the nodose ganglion (e.g. jugular ganglion) or hitchhiking fibers from sensory fascicles. For all morphological fiber types, the variation of fiber counts and occupied areas between fascicles is far greater than the variation between nerves or subjects (Table I), indicating that sampling from a subset of fascicles in a nerve may introduce significant estimation errors.

We show for the first time that most morphological fiber types in the vagus nerve have highly nonuniform distributions across and within fascicles. Fascicles with high UEff fiber counts tend to have low MEff fiber counts, and vice versa (Figure 3D, a); because MAff and MEff fibers are found in separate fascicles, that also means UEff and MAff fibers tend to co-localize in the same fascicles (Figure 3D, b). Many of the MEff fibers are cholinergic, B-type efferents and are the “canonical” vagal fibers targeted by VNS therapies for heart failure and inflammatory disorders. On the other hand, a significant portion of UEff fibers is TH+ (Figure 4B & Suppl. Figure S16), while others could be dopaminergic or non-adrenergic-non-cholinergic fibers. The spatial separation of UEff fibers from MEff fibers suggests that a portion of catecholaminergic (UEff) and cholinergic (MEff) efferent fibers in the cervical vagus nerve occupy different nerve sections, and could in principle be differentially modulated by fascicular VNS. For the first time, we document nonuniform distribution of vagal fibers within single fascicles, as well as sub-fascicular co-localization of different fiber types. For instance, myelinated fibers are frequently located in close proximity with other myelinated fibers (Figure 3D, c-e); UAff fibers also form clusters (Figure 3E, c), in agreement with what has been found in other peripheral nerves with C-type fibers [76].

Our choice of IHC criteria for classifying morphological fiber types has limitations. First, we classified fibers that are positive for NF, MBP and ChAT as myelinated efferents (MEff) (Table 1; Suppl. Table 3; Suppl. Figure S12). However, studies have shown ChAT activity in the nodose (sensory) ganglion in various species [91, 92], suggesting the presence of ChAT^+^ sensory fibers in the vagus. In those studies, chemoluminescence methods were used in the entire ganglia, whose output reflects ChAT activity in both the cell bodies of sensory neurons and the efferent bundles that pass around the nodose ganglion. Using IHC, only few ChAT^+^ neurons were observed in the nodose ganglion in rats and humans [50, 91, 93]. Recently, quantitative IHC studies in transgenic mice (ChAT-EGF) found that sensory neurons in the nodose ganglion are completely devoid of ChAT; instead, ChAT expression is limited to the efferent fibers passing around the nodose ganglion [44]. Despite their potentially small numbers, ChAT^+^ afferent fibers in the vagus nerve will need to be quantified in humans and large animals in future studies.

Second, we classified fibers that are positive for NF but negative for BMP and Nav1.8 as unmyelinated efferents (UEff); because only myelinated fibers are ChAT+, those fibers are also negative for ChAT (Table 1; Suppl. Table 3; Suppl. Figure 12). We found that the vast majority, if not all, of those fibers are positive for TH (Suppl. Figure 16), something that suggests that a significant portion of UEff fibers may represent sympathetic efferents. However, we did not perform a formal quantitative assessment of TH+ fibers in all our samples. Importantly, vagal fibers that are positive for both ChAT and TH have been described [94]. Therefore, the exact count of TH+ fibers in our samples is unknown, as is what fraction of ChAT-unmyelinated efferents they represent.

Third, a portion of UEff-classified fibers may in reality be afferent. Retrograde neuronal tracing studies have shown that some TH+ fibers distributed to the esophagus originate from cells in the nodose ganglion [51, 95] with central axons projecting to the (sensory) nucleus tractus solitarius [95, 96], suggesting that some TH+ fibers are afferent. In fact, it has been estimated that up to 17% of TH+ fibers in the vagus originate in the nodose ganglion, with most of the remaining from cells in the DMV [23]. A study found that as many as 8% of (sensory) cells in the nodose ganglion are TH+ [95], even though more recent studies showed sparse TH staining in the nodose ganglion, corroborated by low overall TH gene expression in nodose neurons [90]. Single-cell RNA sequencing in nodose and jugular ganglion cells showed that all but one cell type express Vglut2 (Slc17a6), a marker for peripheral sensory neurons. The one cell type that does not express Vglut2-displays a “sympathetic” expression profile (expressing Hand2, Tbx20, and Ecel1 genes) and its axons could therefore be considered “efferent” [97].

The role of TH+ fibers in the vagus has been a matter of debate. TH+ fibers in the cervical and sub-diaphragmatic vagus nerve have been well-documented in rats [98–101], cats [102] and dogs [103]. In humans, TH+ fibers are present throughout the vagal trunk at the cranial level, jugular and nodose ganglion, cervical and thoracic trunk, and many vagal branches [15]. In some studies, many TH+ fibers are postganglionic sympathetic fibers with the cell bodies located in the sympathetic chain, entering the vagus nerve through the jugular ganglion [15]. In cats and dogs, the sympathetic trunk and the vagal trunk are fused together, to form the vagosympathetic trunk. In pigs, the sympathetic trunk and the vagus nerve are separate and they run parallel to each other [5]. The presence of the postganglionic sympathetic fibers in the vagus nerve could be due to projection of hitchhiking branch from the sympathetic trunk entering the cervical vagus nerve [8, 22, 103] and the presence of these fibers have been suggested as possible sources of variability in the clinical responsiveness to VNS [22].

Our findings on the distribution of morphological fiber types across and within fascicles have implications for how vagal projections between the brain and peripheral organs may contribute to the strategies followed by the central nervous system for coding interoceptive stimuli [42] and for the autonomic control of organ homeostasis [104]. The co-localization of certain afferent and efferent fiber types in the same fascicles, and in the same sub-fascicular sectors, suggests that the microscopic structure of the vagus nerve may follow a “multiplexing” architecture, in which vagal signals, from/to the same or different organs, are conveyed through projections that lie in close proximity; this organization may pose physical constraints in the spatial arrangement of vagal sensory and motor neurons in the ganglia and the brainstem [42]. Our quantitative data on how fascicles and fibers are arranged in the vagal trunk will be applicable to building anatomically realistic computational models of the vagus nerve, commonly used to optimize design of bioelectronic devices and explain functional outcomes of neuromodulation therapies on the basis of variability in the underlying anatomy and in the electrode-tissue interface [98–101].

### Fascicular vagus nerve stimulation using a multi-contact cuff electrode

Following the example of classic studies that investigated how structure of autonomic nerves explains their function (e.g. [105]) we sought to determine whether the anatomical information gained in our studies could be leveraged for more selective stimulation of the vagus. Selective VNS may provide a means to personalize therapy to account for variability in individual nerve anatomy and engage the innervation of desired organs with fewer off-target effects [3]. The use of multi-contact electrodes for steering of electrical fields during nerve stimulation is an established method to deliver spatially selective stimulation, as it permits activation of subsets of fascicles or fibers. Such an approach has been previously followed in a rat model of VNS, in which reduction of blood pressure was attained without affecting heart rate and breathing rate [106]; it has also been used in a sheep model of VNS, in which selective stimulation attained changes in breathing rate without affecting the heart rate [40]. For the first time in this study, we tested fascicular stimulation effect in a multi-fascicular vagus nerve, with direct relevance to the, also multi-fascicular, human vagus nerve.

Because a large percentage of vagal fibers lie within less than 500 μm from the epineurium (Figure 5A,), a cuff electrode may be a viable selective interface with the vagus nerve. Cuff electrodes for VNS, like the one used in this study, are less invasive than penetrating probes, and have an established safety record, e.g. [107] [https://www.accessdata.fda.gov/cdrh_docs/pdf/p970003s207b.pdf]. By calculating the spatial overlap between fascicles and circular areas centered on individual contacts, representing electric fields that are strong enough to activate nerve fibers, we found that a cuff with 8 contacts along the nerve circumference, provides close to maximal efficacy, with similar selectivity as configurations with more contacts (Figure 5 C). Even though the cuff has not undergone formal preclinical device testing, in principle, it could be used in future clinical tests: the 8-contact configuration is compatible with the fascicular anatomy of the human vagus nerve (Suppl. Figure S26), the helical structure can accommodate a range of nerve diameters (Figure 5C), and the silicone substrate is biocompatible [108] [https://www.accessdata.fda.gov/scripts/cdrh/cfdocs/cfPMN/pmn.cfm?ID=K183437].

Delivering stimulation through single contacts results in differential engagement of fibers innervating the larynx, the lungs and the heart, as demonstrated by the different patterns of physiological and evoked potential responses (Figure 6A). Correlations between physiological and corresponding nerve fiber responses are seen in the recruitment curves, which are also different among different contacts (Figure 6B). This indicates that the differences among contacts in the stimulation intensity thresholds arise because of differential activation of separate fiber populations. The different radial distributions of thresholds for e.g., cardiac vs. respiratory responses indicates that between-contact differences arise because of differences in the underlying fiber distribution, rather than other factors that affect one side of the nerve or the interface (e.g. thicker epineurium or looser placement of the cuff), in which case all 3 registered physiological responses would have similar radial distributions.

The angular separation between the cardiac-, lung- and larynx-selective contacts (Figure 7B) is likely to reflect the underlying anatomical arrangement of fascicles contributing to cardiac, bronchopulmonary and laryngeal branches (Figure 1); however, we did not co-register the exact contact locations with the specific functional anatomy of the stimulated nerves, that being a limitation of our study. The angular separation between the fastfiber-selective contacts and slow-fiber-selective contacts (90° counter-clockwise, Figure 7D) agrees with the separation between contacts eliciting corresponding physiological functions, i.e., fast-fiber-mediated laryngeal EMG and slow-fiber-mediated changes in breathing rate (also 90° counter-clockwise, Figure 7B). Even though physiological and eCAP responses were registered in most animals, we did not align eCAP responses by the contact with the minimum heart rate threshold, but to the contact with the maximum fast fiber amplitude. eCAPs are recorded through a second multi-contact cuff (Suppl. Figure S22), placed at a distance of at least 5 cm from the stimulating cuff and, over that distance, fascicles merge or split and their radial location in the trunk changes significantly (Suppl. Video 1: https://www.youtube.com/watch?v=gaNHijoVB_U). For that reason, the spatial arrangement of fascicles at the stimulating cuff, assessed through physiological thresholds, is different than the arrangement at the recording cuff, and aligning the 2 response patterns would be unlikely to produce interpretable results. eCAP traces from each of the 8 recording contacts reflect the fascicular structure of the nerve at the recording site, when stimulation is delivered to the same site (e.g., Suppl. Figure S22). When comparing eCAPs elicited from different stimulation contacts, we wanted to minimize that source of eCAP variability, hence we averaged the 8 individual eCAP traces elicited by stimuli delivered to each of the stimulation contacts. That way, differences between eCAPs primarily reflect the fascicular structure of the nerve at the stimulating site.

To test whether fascicular VNS is feasible in awake conditions, and to establish its stability across experimental sessions, we administered fascicular VNS from the same cuffs for up to 3 weeks, in 2 chronically implanted swine (Figure 7E, a). In awake conditions, activation of the cough reflex is the physiological response observed at lowest stimulus intensities; indeed, cough and throat pain are common side effects in patients receiving cervical VNS [109]. Intensity thresholds for eliciting the cough reflex are asymmetrically distributed across stimulating contacts in both animals (Figure 7E, b1 and c1), suggesting that the severity of this off-target effect can in principle be reduced by fascicular VNS. To our knowledge, this is first report of a chronic VNS implant in swine, and the first report of cough reflex thresholds from VNS in awake swine. Cough reflex thresholds progressively increase post-implantation (Suppl. Figure S27), similar to what we have observed in chronic VNS implants in mice [110] and rats [66]. Progressive threshold increases likely reflect the progressive development of fibrotic tissue around and inside the implanted cuff rather than major loss of nerve fibers, for which we found no histological evidence (Suppl. Figure S25). Despite the increase in thresholds, the shape of the threshold distribution around the contacts remains consistent across time in both animals (Figure 7E, b1 and c1; Suppl. Figure S27), suggesting that fascicular VNS may be a feasible strategy to suppress this off-target effect in chronically implanted patients.

During fascicular VNS, we observed eCAPs comprised of only fast and slow A-fiber components. This may be related to the large areas covered by large myelinated fibers (Table I). At the same time, we did not activate small, unmyelinated fibers (e.g., C-type), as evidenced by the lack of slow fiber eCAP responses. Small, unmyelinated fibers constitute >80% of the total vagal fiber count, however, they occupy <10% of the entire area the nerve (Table I). Small, unmyelinated fibers have high activation thresholds [111, 112]. In our experiments, we used smaller current intensities; by using intensities high enough to activate C-fibers, larger fibers would have been activated as well, and hence the spatial selectivity of other physiological effects would have disappeared. At the same time, most unmyelinated fibers have a more uniform distribution across fascicles (Figure 3). For that reason, it is less likely that fascicular VNS would result in asymmetric C-fiber responses in eCAPs—however our study has not tested that possibility.

Our physiology studies demonstrate that spatially selective, fascicular stimulation using a multi-contact cuff electrode is a meaningful strategy for selective VNS in the multi-fascicular vagus nerve of the swine, and it may be feasible in the human cervical vagus nerve, which is also multi-fascicular. However, the degree of organ- and function-selectivity of our approach study is limited: no one contact is 100% selective for a single organ or for a single fiber type. This is expected, given the many, small fascicles in the swine nerve, the epineural placement of the stimulating electrode, and the non-selective nature of stimulus waveforms used. Because the human cervical vagus nerve has fewer and larger fascicles it may be an even better fit for selective, fascicular VNS with cuffs with relatively few contacts. To translate these findings into clinically-relevant selective VNS therapies, anatomy-guided electrode contact configurations used with fiber-selective stimulus parameters will need to be tested [113]. To deliver precision VNS therapies, the functional anatomy of the vagus nerve will need to be mapped in individual subjects. Towards that end, the large-scale fascicular structure of the swine vagus nerve has been visualized with noninvasive, high resolution ultrasound imaging [68] or with electrical impedance tomography of nerve activity [114, 115]. Our results show that functional nerve mapping can be also performed by quantifying physiological and/or nerve fiber responses delivered through different contacts and visualizing their spatial distributions around the nerve (Figure 6B, Figure 7A). Physiological responses and eCAPs elicited by stimulation can be registered and processed in almost real-time in humans or experimental animals to generate functional nerve maps and calibrate and optimize VNS parameters [60].

## Methods

### Animal surgeries

The effects of VNS on physiological and neural response were examined in male swine (Yucatan; n=13, ~50kg; Yorkshire, n=1, ~30 kg). All of the animal protocols and surgical procedures were reviewed and approved by the animal care and use committee of Feinstein Institutes for Medical Research and New York Medical College.

Anesthesia: For a 12-hour period before surgery, the animals were maintained on a no-food and no-fluid regimen (NPO). On the day of surgery, the animals were weighed and sedated with mixture of Ketamine (10-20 mg/kg) and Xylazine (2.2 mg/kg) or Telazol (2-4 mg/kg). Propofol (4-6 mg/kg, i.v.) was used to induce anesthesia, and following intubation, the anesthesia was maintained with isoflurane (1.5-3%, ventilation). Mechanical ventilation was provided and was turned off only at time breathing rate changes were measured during VNS. Animals were placed on a table in a supine position with normal body temperature between maintained between 38°C and 39°C using a heated blanket and/or a hot air warmer and monitored with a rectal probe thermometer. Blood pressure and blood oxygen level were monitored with a BP cuff and a pulse oximeter sensor. The depth of anesthesia was monitored by assessing heart rate, blood pressure, respiration, and mandibular jaw tone. All surgeries were performed using sterile techniques.

For survival surgeries, a fentanyl patch (25-50 mcg/hr) and buprenorphine (0.01 mg/kg s.q.) were provided to the animal at the end of surgery to alleviate post-operative pain. After gradually weaning off the ventilator and extubated and when physiological signs returned to normal, animals were returned to the home cage and were monitored 3 times/day for 72 hours. Cefazolin (25 mg/kg p.o.) was given for 5 days. Carprofen (2-5 mg/kg, s.q.) was administered when the animal showed signs of pain. Animals were allowed 10 days for full recovery before any tests were performed.

### Cervical vagus nerve and laryngeal muscle implants

Under the surgical level of anesthesia, a 4-5 cm long incision was cut down to the subcutaneous tissue on the right or left cervical region of the neck, 2 cm lateral to the midline. Subcutaneous tissues were dissected and retracted to expose the underlying muscle layers. The sternomastoid muscle was bluntly separated and retracted laterally away from the surgical field. The carotid sheath was exposed and the vagus nerve was identified and isolated away from the carotid artery. One or two helical cuff electrodes (CorTec, Germany) were placed on the rostral (around 2cm to nodose) and/or caudal sites with a distance of 4 to 5 cm after isolating the underlying 2 cm long section for each electrode. The helical cuff included 8 contacts (0.5mmX0.5mm) evenly distributed (1mm between two contacts) around the circumference and 2 ring-shaped (0.5mmX7.4mm) return contacts. The middle portion, between the two isolated sites was left intact within the carotid bundle to minimize the extent of surgical manipulation and trauma to the nerve. For laryngeal muscle recording, after the hyoid bone was blunt dissected, Telfon-insulated single or multi-stranded stainless-steel wires were deinsulated at the tip about 1 mm, and inserted in thyroarytenoid (TA) of laryngeal muscle with a needle. For survival surgery, the incision was closed, the leads and connectors of the implanted cuff electrodes and EMG electrodes are tunneled from the surgical field in the neck area to the back of the neck subcutaneously and then to the top of the skull, where they are externalized. Several stainless-steel skull screws are implanted in the exposed area of the skull by drilling small burr holes. The connectors and the anchoring screws are secured with dental acrylic, and then a titanium casing was attached around and secured to the skull around the connector for protection of the implants (Figure 7 E,a).

### Physiology experiments

Breathing rate: Breathing was monitored by using a respiratory belt transducer (TN1132/ST), which was placed around the belly or chest and connected to a bridge amplifier (FE221, ADI). The change of the belt pressure with breathing represented the breathing cycle and was used to derive the breathing rate. The mechanical ventilator was turned off when measuring the effects of vagus nerve stimulation on breathing.

ECG: The skin around the wrist was shaved and conductive gel was applied to patch electrodes which were subsequently adhered to the skin, ECG was recorded using a 3-lead patch electrode configuration and amplified using a commercial octal bio-amplifier (FE238, ADI).

Heart blood pressure: Ultrasonic guidance (H8018QE and Vivid IQ v203 4D) was used to pass a 19 G angiographic needle through the skin at a 60-degree angle to the target vessel. Once blood backflow was detected, a soft guidewire (0.035 inches) was introduced into the vessel. A 5-Fr micro manometer-tipped catheter (SPR-751, Millar Inc) was advanced into the aorta or right ventricle, and pressure was recorded continuously after amplification (FE228 ADInstruments). The catheter locations were confirmed by verifying the characteristic phasic blood pressure waveforms corresponding to different placements.

All physiological signals were first digitized and then acquired at 1 kHz (PowerLab 16/35, ADI) and then visualized using LabChart (ADInstruments Inc).

Neural signal and EMGs were sent to the data acquisition system through a headstage (RHS 32, Intan Tech). Signals were digitized at a sampling frequency of 30 ks/s with an acquisition system (Intan 128 recording/stimulating, Intan Tech).

### Multi-contact cuff electrode device

#### Methodology for cuff design specification

To determine an “optimal” number of contacts in the cuff electrode for targeting the fascicular anatomy of the cervical vagus nerve of swine, we estimated the expected efficacy and selectivity of different contact configurations. Outlines of the epineurium of the entire nerve and of the perineurium of each of its fascicles were extracted from H&E images of sections taken through the mid cervical region of 8 extracted vagus nerves. Each nerve or fascicle outline was converted to a matrix of x-y coordinates and fitted with an equivalent ellipse centered at the center of mass of the outline. The equivalent ellipse was calculated by iteratively determining the length of short and long axis and angular rotation that minimized the difference between the area of the ellipse and the area of the fascicle outline. The ellipse surrounding the nerve outline represents the epineural cuff. Points representing the electrode contacts were placed on the nerve-equivalent ellipse at regular intervals (e.g. for 4 contacts, the angular separation between contacts was 90 degrees) (Figure 5A). For each nerve, that placement was repeated 100 times, with a random angular placement of the configuration on the ellipse. Circles centered on each of the contacts were drawn, representing the electrical fields that activate all fibers in the area of fascicles that overlaps with those circles. The strength of electrical fields is represented by the radius of the circle, and is expressed in units of the nerve radius (length of the short axis of the nerve-equivalent ellipse). The degree of overlap between a fascicle area and a field area is calculated for each fascicle, as % of the fascicle area; if the overall is >75% of the fascicle area, that fascicle is considered “activated”; at <15%, the field is considered “spared”. Efficacy of a configuration for a given field strength is defined as the percentage of fascicles that are activated by all contacts in a configuration, averaged across all 100 random placements on a nerve. Selectivity of a configuration for a given field strength is defined as the percentage of nerve fascicles that are activated by each 1 contact, averaged across all contacts in a configuration and across all 100 random placements on the nerve.

#### Electrode fabrication

The helical cuff is based on the self-spiraling cuff with laser-machining method [116, 117]]. At first, a polymeric release layer was made on a ceramic substrate (Al2O3, 100 x 100 mm^2^, thickness: 0.635 mm) through a mechanical carrier. Then, silicone rubber 1:1 n-heptane-diluted (Carl Roth GmbH + Co. KG, Karlsruhe, Germany) and MED-1000 (NuSil Technology LLC, Carpinteria, CA, US) is spin-coated on top, and followed by laser-structured using a picosecond Nd:YAG laser (HyperRapid NX, λ= 355 nm, Coherent, Inc., Santa Clara, CA, US). Platinum/Iridium (90/10) foil (thickness:25 μm, Goodfellow GmbH, Bad Nauheim, Germany) is laminated, and laser-cut and glued to the pre-stretched MED-1000 silicone rubber foil, which was laser cut. Helical cuff electrode array is composed of 8 square contact pads (0.5 × 0.5 mm), which are arranged evenly with 1mm pitch to wrap around nerve in helix shape (Figure 5C). 8 contacts were chosen based on the optimization of efficacy and selectivity for vagus nerve stimulation (figure 5A,B). Two reference electrodes (0.5X7.4mm) are parallel to the array on proximal (distance to array 2mm) and distal side (distance to array 1mm). All electrodes are made from Pt/Ir (90/10) material and are embedded in medical grade silicon rubber and laser cut. The impedance for each contact of the array is less than 10 KQ, and the impedance of reference electrode is less than 1K) at 1Khz. The charge storage capacity for each electrode is from 3000 to 5000 μC/cm^2. Each cuff electrode can be used multiple times without performance degradation (figure 5D and 5E).

#### Electrode characterization

##### Cleaning

Prior to characterization, the electrodes were soaked in a 1% tergazyme solution and placed in an ultrasonic cleaner for 5 minutes. They were then allowed to soak in the tergazyme solution (room temperature) for at least 1 hour, following which the solution was decanted. The electrodes were rinsed with DI water, then placed back in the ultrasonic cleaner for 5 additional minutes in DI water. The water was then decanted and the electrodes were rinsed once more before performing measurements.

##### Electrochemical impedance spectroscopy (EIS)

The EIS measurements were done using a 3-electrode set up. A graphite rod was used as a counter electrode along with an Ag/AgCl reference electrode. The electrode under test along with the reference electrode were placed in a 7.2 pH 1x PBS solution, then sonicated for 5 minutes to remove any air bubbles from around the cuff structure. They were then placed in a Faraday Cage, where the counter electrode was introduced into the solution. The EIS measurements of each electrode were taken separately, from a range of 100,000 Hz to 0.2 Hz.

##### Charge storage capacity (CSC)

The cyclic voltammetry (CV) plots were taken in a similar manner as the EIS measurements. The cuff electrode and reference electrode were again placed in the PBS solution and sonicated for 5 minutes to remove air bubbles, after which the counter electrode was introduced. The CV measurements were performed over 10 cycles and swept from −0.6 - 0.8V at a scan rate of 100 mV/s. In calculating the charge storage capacity, the first and last sweep were disregarded, and the remaining sweeps were averaged and integrated below zero to yield the cathodic CSC.

### Vagus nerve stimulation and physiological data analysis

#### Physiological thresholds

Biphasic stimulus pulses were delivered using an isolated constant current stimulator (STG4008, Multichannel Systems) through each of the 8 contacts of the cuff electrode device. Physiological threshold (PT) was detected from stimulus trains of 10 s durations (30 Hz, pulse width 200 μs). There was at least 30 s long duration between successive trains to ensure that physiological measurements had reached a steady state before a new train was delivered. We define as threshold the lowest stimulation intensity to induce 5-10% drop in heart rate (HR threshold), ~50% drop in breathing rate (BR threshold), and a visible response in laryngeal EMG (EMG threshold). The 50% drop in BR was chosen because of the sharp, sigmoid relationship between intensity and drop in BR, documented in [60]. In awake experiments we measured the threshold for evoking cough reflex, as the lowest stimulation intensity to induce coughing. In those experiments, we measured changes in heart rate at cough reflex threshold intensities. The thresholds for each of the 8 contacts were tested in a random order.

#### Evoked compound action potentials (eCAPs)

After removing the DC component (<1 Hz) from recordings, evoked compound action potentials (eCAP) from the vagus nerve were averaged across all of the individual stimulus pulses within each train. The responses (peak-to-peak) from eCAP within predefined time window were classified into different fibers based on the conduction speed. Given the distance between recording and stimulation electrode, ~45-60 mm, the first peak latency within 1.5 ms (conduction speed, >33 m/s) was classified as a fast fiber response, consistent with activation of large, myelinated A-type fibers; the late peak after 2 ms (conduction speed, <20 m/s) was classified as slow fiber response, consistent with activation of smaller, myelinated fibers (Aδ- and B-type). In most of the recordings, EMG contamination observed in eCAP recordings could be blocked by Vecuronium. Long latency C-fiber response was not observed in our tests. The eCAP responses (peak-to-peak amplitude) from eight contacts on the second cuff were recorded simultaneously while stimulating each of the eight contacts; however, because of the high degree of similarity between individual eCAP traces elicited from the same stimulated contact (Suppl. Figure S22), eCAP traces were analyzed after averaging across the recording contacts.

#### Recruitment map and population responses

The responses from laryngeal muscle (peak-to-peak EMG amplitude), breathing rate change and mean eCAPs responses (of 8 contacts) of different fibers were characterized to inform recruitment maps during increasing stimulation intensity across 8 stimulating contacts.

To characterize the spatial selectivity of the 8-contact cuff, PTs and eCAPs from each contact were plotted on a polar-plot with mean direction and resultant vector length. To show the spatial selectivity across animals and compare difference between physiological responses, we aligned the lowest thresholds of HR on the 12 o’clock position from all animals, and then PTs of BR and EMG was aligned to the HR. Then the PTs were normalized between the lowest (to 0) and highest (to 1) to give the population (mean+/-SEM) across animals. Similarly, we first aligned the highest responses (eCAP of peak-to-peak amplitude) of fast fiber to 12 o’clock position, and then we get the population eCAPs (mean+/-SEM) across all animals, and finally the population response from slow fiber eCAPs were alighted to the fast response for each swine and averaged across animals (mean+/-SEM). ANOVA was used to test the spatial selectivity across contacts (anova1, *p<0.05).

### Euthanasia and surgical dissection nerve samples for micro-computed tomography

At the end of the study, the swine was euthanized with a solution (Euthasol) (A dosage of 1 ml/10 pounds BW, i.v.). Euthanasia solution (Euthasol) was injected intravenously. Death was confirmed using the ECG signal and the arterial blood pressure signal. Absence of a pulse in both signals confirmed cardiac death. After euthanasia, both the left and right cervical vagus nerve were dissected from above the nodose ganglion to the end of the thoracic vagus; during that time, nerve branches, still attached to the nerve trunk, were isolated using blunt dissection up to the respective end organ (heart, lung or larynx). A fine suture loop (6-0) was placed on the epineurium of each branch, close to its emergence from the trunk, as a label. Detail record of every branch and it’s innervating organ was maintained during dissection with identifiable suture labels placed in each branch. The entire nerve along with branch specific sutures label was photographed with the ruler before being fixed. The sutures used were radiopaque and hence can be see clearly in the micro-CT scanned images. The samples were fixed in 10% formalin for 24 hours, then transferred to Lugol’s solution (Sigma, L6146) for five days to achieve optimal fascicular contrast for the micro-CT. Nerve samples were wrapped in a less dense material such as saran wrap and mounted in position on a vertical sample holder tube. The samples were scanned using Bruker micro-CT skyscan 1172 with a voxel size of 6 μm resolution. Volume rendering was done using Bruker CTvox v3.3.1 to obtain a 3D reconstructed view of the nerve (Suppl. Figure S2-S4).

### Segmentation and tracking of fascicles in micro-CT data

The automated fascicle tracing approach consists of two stages: segmentation and tracking. Instance segmentation on micro-CT images was performed using the Matlab Deep Learning toolbox, Mask R-CNN model implementation (Mathworks, Inc.). The mask R-CNN model comes pretrained based on the COCO image dataset and transfer learning was applied by freezing the Resnet50-coco backbone during training which forces the model to utilize the existing COCO features from the Resnet50 subnetwork while updating the weights of the region proposal network and mask head. Since micro-CT images are significantly different than COCO images (grayscale intensity images as opposed to color photographs, content is at a much smaller spatial scale), to achieve adequate performance, 80% of images that were annotated from the initial nerve needed to be included in the training data. Once the fascicle detection mask R-CNN model was trained (stage 1), it was updated for each new nerve sample with fewer annotated slices (stage 2); we annotated 1 out of every 50 micro-CT slices on subsequent nerves. The annotated images included three categories: fascicle, false alarm on nerve, and false alarm off nerve. The false alarm classes were created by storing the segmentation results that didn’t overlap with the ground truth annotations and then subcategorizing them based on the average pixel intensity within the detection masks. We chose an initial learning rate during training for each stage, *λ_0_*, to 1.2 * 10^-5^, that was decreasing with each epoch, based on the formula *λ=λ_0_/(1 + τ(epoch-1))*, where *τ* is a decay rate of 0.01. The model was trained for 18 epochs resulting in training iterations which is on par with the number of training iterations used to train the original Mask R-CNN model [118]; stochastic gradient descent with momentum was used to update the network weights with a momentum of 0.9 and a minibatch size of 1. We chose a mask RCNN model as our detector because it performs well on the detection task and can readily be updated in individual animals if necessary. Shape-based, pixel-based, and texture-based features were extracted for each detected fascicular cross-section. Shape-based features consist of the centroid, circularity, eccentricity, irregularity, area, orientation, major and minor axes lengths, solidity, and the weighted centroid. Pixel-based features consisted of the mean pixel intensity and central moments like variance, skewness, and kurtosis. Texture-based features were derived from the gray-scale cooccurrence matrix and include contrast, correlation, energy, and homogeneity. A minimum area constraint of 120 pixels was imposed based on the smallest area in the annotated data.

After the detected fascicular cross-sections for each slice of the micro-CT data were determined, the structure of the nerve was estimated by tracking the fascicular cross-sections longitudinally. The tracker filters and structures the detections to construct a graph that captures properties of the fascicular cross-sections (shape and texture metrics) at each node and encodes branching and merging via the graph edges. We used a windowed, density-based-clustering approach. The distances between the detected fascicles within the micro-CT slice window were computed as *1 - J*, where *J* is the Jaccard similarity (a.k.a. intersection over union). DBSCAN was sued to cluster the detected fascicular cross-sections at each window position, with a minimum of 3 samples per cluster and an epsilon neighborhood of 0.6. A track was extended if more than 2/3 of the detections are the same between the adjacent window positions. Any detections that are not included in a track was discarded, except for detections that correspond to manual annotations. Once this step was completed, a post-processing algorithm was run on the tracks to connect broken segments that result from missed detections and detect branching and merging events. Individual fascicles contributing to the different vagal branches were manually identified at the branching points (Suppl. Figure S2-S4). The tracked fascicle trajectories were then color coded from the caudal end of the branching point to the rostral end of the nodose ganglion (Figure 1).

### Immunohistochemistry

Cross-section of the vagus nerve at 5 μm thickness was obtained through paraffin sectioning. Sections were stained for myelin basic protein (MBP), neurofilament (NF), choline acetyltransferase (ChAT) and Nav1.8 using standard IHC protocol [119] to identify the number and distribution of the myelinated and non-myelinated fibers inside different fascicles. Briefly, sections were subjected to deparaffinization procedure using xylene and ethanol rinse and then washed with distilled water for 10 min. Antigen retrieval was performed by briefly subjecting the samples to 1 x citrate buffer to unmask the antigens. Sections were rinsed with 1x Tris-buffered saline and subjected to 1 hour of blocking step using one normal goat serum. Sections were incubated with anti-Neurofilament (1:500, Abcam AB8135), anti-Myelin basic protein (1:500, Abcam AB7349) and either anti-ChAT (Millipore Sigma, AB144P) or anti-Tyrosine Hydroxylase (abcam, ab112 (“efferent IHC panel”) or SCN10A monoclonal antibody for swine (Nav1.8, MA5-27663 Fisher) and SCN10A polyclonal antibody for humans, alomone labs, ASC-016 (“afferent IHC panel”), overnight at 4 °C in a shaking humidifier. The following day, sections were rinsed and incubated with the secondary antibody for two hours at room temperature. Slides were then rinsed thoroughly with TBS buffer three times, and cover glass was mounted on the sides with the fluoromount (Vector labs, H-1700). The slides were then imaged using ZEISS LSM 900, confocal laser scanning microscope, and BZ-X800 all-in-one fluorescence microscope at 100x magnification.

To ensure the quality of IHC and validate the specificity of the antibodies, we undertook the following studies.

a. Transgenic experiments using pigs are not feasible, hence we used transgenic mice to validate antibody specificity. In our study, we used transgenic mice that express enhanced fluorescent protein (EYFP) with ChAT expression as a positive control. Vagus nerves from these transgenic animals were used to validate binding specificity of the ChAT antibody with the same specific antibody used in the pigs (Supplementary Figure S19) and also to define the ChAT staining pattern of single fibers, information that was used to train the segmentation and classification algorithm.
b. By omitting the primary (ChAT) antibody run and using only the secondary antibody in tandem sections as negative control, we observed no nonspecific binding in tissue.
c. We studied the localization of antigens to well-characterized specific locations in the cell to confirm the validity of binding. Antibodies against NF, MBP, ChAT and Nav 1.8 are well characterized with regard to their binding location in axons; our results are consistent with those studies ([1],[5],[120]). We found that NAV1.8+ fibers have a punctate expression pattern. To confirm that that staining pattern represents C-fibers and not merely precipitation of the Nav1.8 antibodies, we double-labeled the sections with Schwann cells marker (S100) and Nav1.8. We found that Schwann cells engulf Nav1.8 positive fiber clusters (Figure 3B). Unmyelinated small diameter C-fibers clusters are known to be ensheathed by nonmyelinating Schwann cells called Remak Schwann cells [121],[122]; Nav1.8 is known to have a clustered distribution along the length of the axons [123] both consistent with our current findings. Together our results prove anti Nav1.8 antibody used in this study target C fiber selectively.
d. Higher or lower dilution factors of the antibody will result in false positive or false negative results. To address this, we diluted the antibody to the optimal point just enough to detect the antigen of interest leading to minimal background staining. Further during imaging, exposure levels were kept at the optimal levels to obtain higher signal to noise ratio.
e. BLAST homology analysis: To identify unmyelinated afferent fibers in swine tissue, we used a monoclonal anti-Nav1.8 antibody, which, compared with polyclonal antibodies, binds with higher affinity and specificity to its antigen. However, this antibody failed to identify afferent fibers in human vagus nerve sections. Therefore, we tested a polyclonal anti-Nav1.8 antibody and our preliminary results show comparable staining of afferent fibers as seen in pig nerve sections. To further validate the new polyclonal antibody, we ran a BLAST analysis of the amino acid sequence of the rat antigen against which it was raised (amino acids 1724-1956: ENFNVATEESTEPLSEDDFDMFYETWEKFDPEA) and found that it had 100% identity match with human Nav1.8 (several isoforms).

### Fiber segmentation and feature extraction

We designed a set of algorithms and a data pipeline to segment and extract nerve, fascicle and single-axon level features from IHC images of the vagus nerve. First, we developed a fiber segmentation and feature extraction algorithm by combining standard computer vision methods with statistical modeling and geometrical manipulations, including Gaussian mixture models and Voronoi tessellation, and achieved a single-axon detection accuracy of 82% and myelination classification accuracy of 89% [79].

To improve the detection accuracy, we further trained a Mask R-CNN deep convolutional neural network on axon instance segmentation. The model was pretrained on the COCO Instance Segmentation task [124]. We generated training images by first annotating fibers using the more standard algorithm, followed by manual correction of 80 fascicles, done in an ordinary image editing software. We estimated that pre-annotating using the original algorithm reduced the time of correction by four times compared to manual annotation from scratch.

#### Fiber identification

Neurofilament: To detect neurofilament positive pixels, we empirically selected a threshold brightness in the neurofilament color channel (154 out of 256) above which pixels were classified as positive. We further discovered connected components of such pixels to arrive at blobs of neurofilament. Blobs containing less than 10 pixels were excluded as noise. The remaining blobs were broken divided into individual fibers by fitting Gaussian mixture model on them of different number of Gaussian distributions (i.e. number of fibers) and then selecting the best fit determining the amount and location of enclosed axons.

Myelin: Myelin positive pixels were detected using the same technique as described previously [79]. Briefly, myelin channel pixels surrounding each detected neurofilament positive axon were analyzed. Pixels were classified as myelin positive when their brightness reached 128 out of 256. First, the outside pixel shell of neurofilament positive fibers were taken and neighboring pixels were recursively checked (for a maximum of 5 recursive steps) for myelin positive pixels. If more than 30% of fiber neighboring pixels were found to be myelin positive, the said fiber was classified as myelinated.

ChAT: Similar to Neurofilament and myelin pixel classification, we empirically selected a brightness threshold, applied to the chat color channel, of 131 out of 256. ChAT positive pixels overlapping myelinated axons were counted separately for each fiber and if more than 5% of pixels were found to be chat positive, we regarded the fiber as chat positive.

The final set of segmentations was performed by the combination of the standard computer vision and deep learning algorithms: we let Mask R-CNN detect axons first, then opted for traditional methods to detect and divide overlapping neurofilament blobs that were ignored by the deep learning model. This combination achieved a fiber detection accuracy of 96% [125]

After the instance segmentation of neurofilament and the assignment of myelin to detected fibers, we extracted a set of features for each axon. These features include the longer and shorter diameters, the perimeter, and area of the fiber, in addition to the thickness of the myelin, and the polar coordinates of the fiber relative to the center of mass of the enclosing fascicle.

We manually annotated nerve cross-sections on the fascicle level to derive polar coordinates of fascicles and their distance from the epineurium. Moreover, we extracted fascicle-level features including the perimeter of the perineurium, the area of the fascicle enclosed by the perineurium, and the total count and area of each fiber type within a fascicle.

Nav1.8: To generate ground truth data for the deep learning instance segmentation network, we performed manual annotation of NAV1.8 stained images to determine the density of C fibers within bundles given the area of the enclosing bundle. Due to the low resolution of light microscopy at the range of 1 μm, we used Airyscan equipped super high resolution confocal microscope captured laser microscopy images of parts of fascicles, from which single NAV1.8 positive C fibers can be distinguished (Suppl. Figure S20, a). These high resolution images were manually annotated bundle-by-bundle to amass counts of fibers within more than 300 bundles extracted from two animals and four different fascicles. To predict number of NAV1.8 positive C fibers from lower resolution light microscopy images (Suppl. Figure S20, c) a linear regression model was fit on the C fiber counts given the area of the bundle (r^2^ = XY) (Suppl. Figure S20, d). We subsequently employed the linear regression model to estimate the number of NAV1.8 positive C fibers in clouds of NAV1.8 stains, detected as bundles, in light microscopy images (Suppl. Figure S20, e). As the linear regression model estimates are real numbers, we round those numbers to the closest integer for each detected bundle to get actual counts.

All images are preprocessed before the segmentation to enhance contrast and to remove noise. Myelin-neurofilament-ChAT stained images were first adjusted by adaptive histogram equalization (dividing each fascicle image into a 32-by-32 grid), then color clipped by defining upper and lower threshold of the clip for separately each cross section manually. The thresholds for each cross-section and channel were selected to remove background noise and to render the stains close to binary (distinct foreground and background), which removed the brightness variance between cross-sections allowing us to use the same parameters for all cross-sections in the subsequent segmentation process. Myelin-neurofilament-NAV1.8 images were not processed by adaptive histogram equalization, but were color clipped as described above. In the subsequent segmentation process, we ignore all individual neurofilament and NAV1.8 blobs that are smaller than 0.225 μm^2^, which we found to be background noise rather than actual fibers.

#### Fiber population data analysis

Once we detected and localized fibers and fascicles, and further extracted features from them, we performed subsequent analysis to map the distribution of different fiber types within-fascicle, and within the whole nerve.

We calculated within-fascicle counts and areas of the four fiber types, namely, myelinated efferents (MEff), myelinated afferents (MAff), unmyelinated efferents (UEff) and unmyelinated afferents (UAff). The per-fascicle counts are either presented in their absolute values (Figure 3A, I-L) or as percentages normalized by the total fiber count per fascicle (Figure 3C).

For each fiber type we estimated the ratio of its neighboring fiber types in a 15 μm radius (Figure 3E, c). For each fiber in every fascicle at the mid-cervical level, we captured its neighboring fibers using k-nearest neighbors, then filtered out the ones outside of the 15 μm radius of the fiber. We accumulated the counts of neighboring fiber types for each center fiber type; normalized by the total count of neighbors we calculated the overall ratio of a fibers a given type being in the vicinity of a given fiber type. To further describe the tendency of different fiber types to share space, we compared their total numbers in each fascicle (Figure 3D, a & b). Within-fascicle mixing was analyzed by splitting each mid-cervical fascicle into a grid of 30-by-30 μm nonoverlapping spatial sectors, establish the fiber counts by type in the sectors, then finally count the number of sectors of a certain composition and represent the propensity of two fiber types to share a space of 30-μm-width fascicle area by showing a 2D histogram of the distribution of compositions of the two fiber types (Figure 3E, a & b)

To examine distribution of fiber types within the nerve, we measured the percentage of a fiber type being positioned in fascicles inside rings of 100-μm width at different distances from the epineurium. We show the overall distribution of fiber types as the function of epineurium distance combining data extracted from nerves of different animals at the mid-cervical level (Suppl. Figure S14, A). We also present the same information for individual nerves as cumulative distributions (Suppl. Figure S15). We grouped fascicles according to their effective diameter into 50 -μm bins and measured the percent area occupied by each fiber type within each fascicle normalized by the area of the fascicle. We then computed the average and standard deviation of such fiber type percentages per effective diameter bin (Suppl. Figure S14, B).

As we manually annotated single fibers of MEff, MAff, UEff and UAff to generate ground truth data for our instance segmentation and NAV1.8 positive C-fiber counting model, we had access to accurate single-fiber features. To gauge the size of the different fiber types, we measured the effective diameter of the manually annotated single fibers of each type and presented their distribution (Suppl. Figure S13).

## Acknowledgments

This work was supported by a grant from United Therapeutics Corp. to SZ and by an NIH-SPARC grant to LM (1OT2OD026539). The authors would like to acknowledge Drs. Betty Diamond, Loren Rieth, Thomas Coleman, Valentin Pavlov, Sangeeta Chavan and Eric Chang for helpful discussions.

## Author contributions

NJ, VT and WS designed experiments, performed experiments and collected data, analyzed data, interpreted data and wrote the paper. AV and TL analyzed data, interpreted data and wrote the paper. JNT and KQ performed experiments and collected data. IM, YCC, MR, AD and AA collected and analyzed data. ZN, BTV, KJT, YA, TDC interpreted data and wrote the paper. LM, MFB and SCL performed experiments and collected data, interpreted data and wrote the paper. TPZ designed experiments, analyzed data, interpreted data and wrote the paper. SZ designed experiments, performed experiments and collected data, analyzed data, interpreted data, wrote the paper and managed the project. SZ and LM acquired funding.

