## Supplemental Figures for "Organ- and function-specific anatomical organization of the vagus nerve supports fascicular vagus nerve stimulation" (from another version of this preprint: version 7, 2022-11-01)

**Supplementary Information**
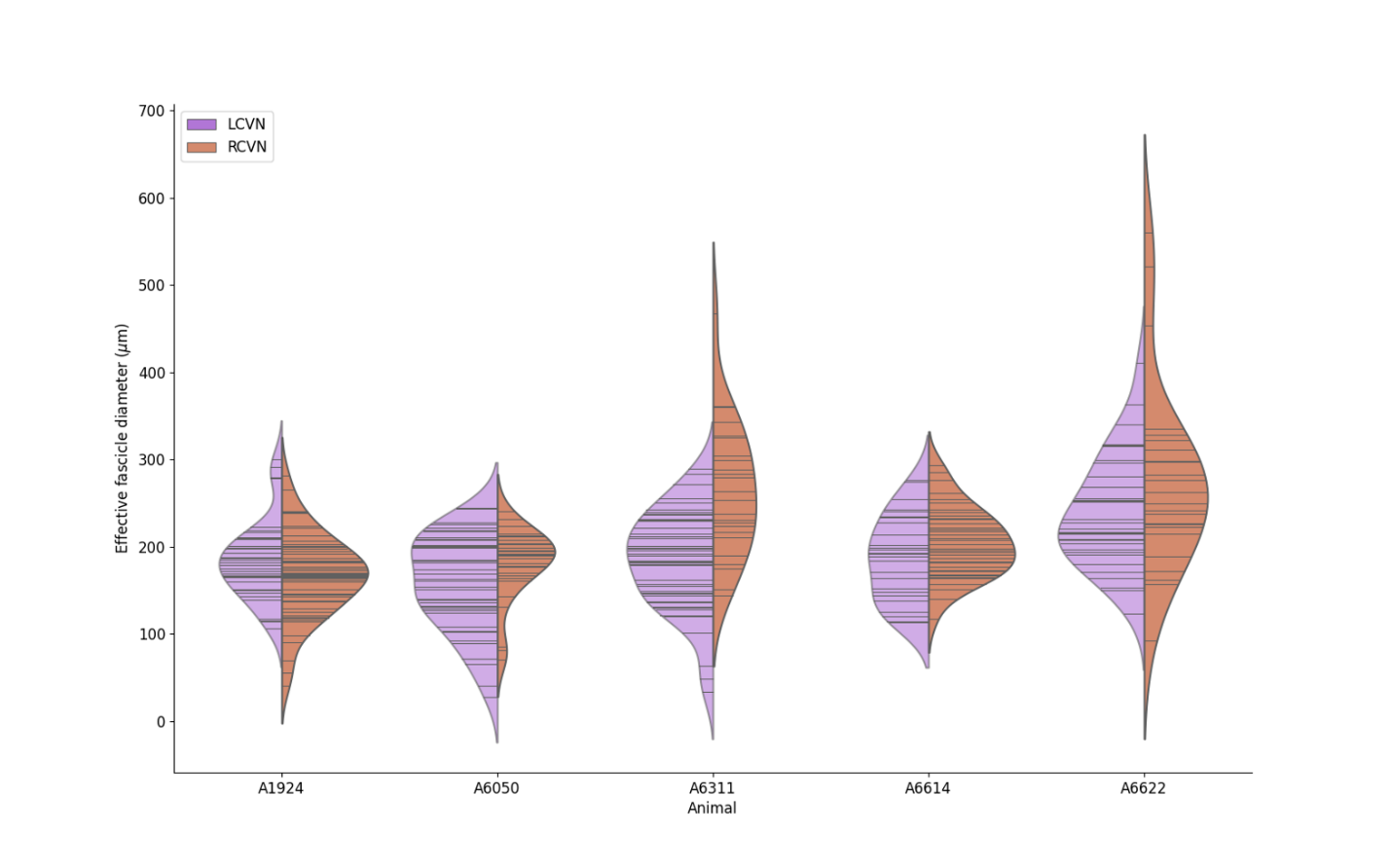

**Supplemental Figure S1:  Effective diameter of fascicles in different animals in both left and right cervical vagus nerve.** Violin plot showing the distribution of effective diameter of fascicles for each animal at the mid-cervical region.

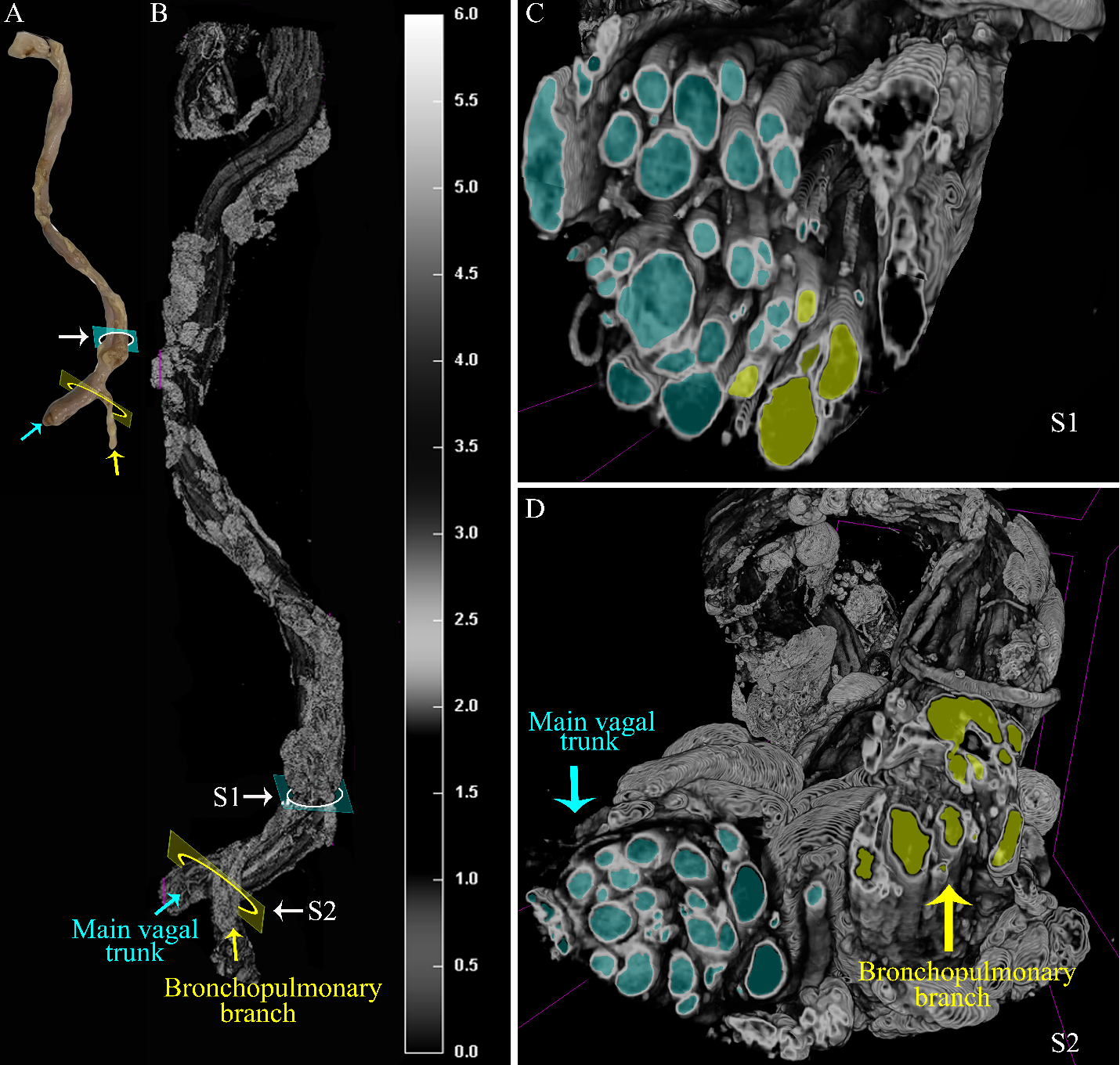

**Supplementary Fig S2**. **Micro-CT image of the Bronchopulmonary branch**. A) Representative section of the nerve showing bronchopulmonary branch of the pig right vagus nerve. B) Micro-CT image of the vagus nerve showing bronchopulmonary branch of the main vagal trunk. C) Fascicles in the bronchopulmonary branch was manually annotated in the segmented micro-CT images with yellow color and was traced back to the caudal part of the main trunk. D) Micro-CT image of the bronchopulmonary branch of a Yucatan pig, showing 15 fascicles at distance of 24 cm from the nodose ganglion.

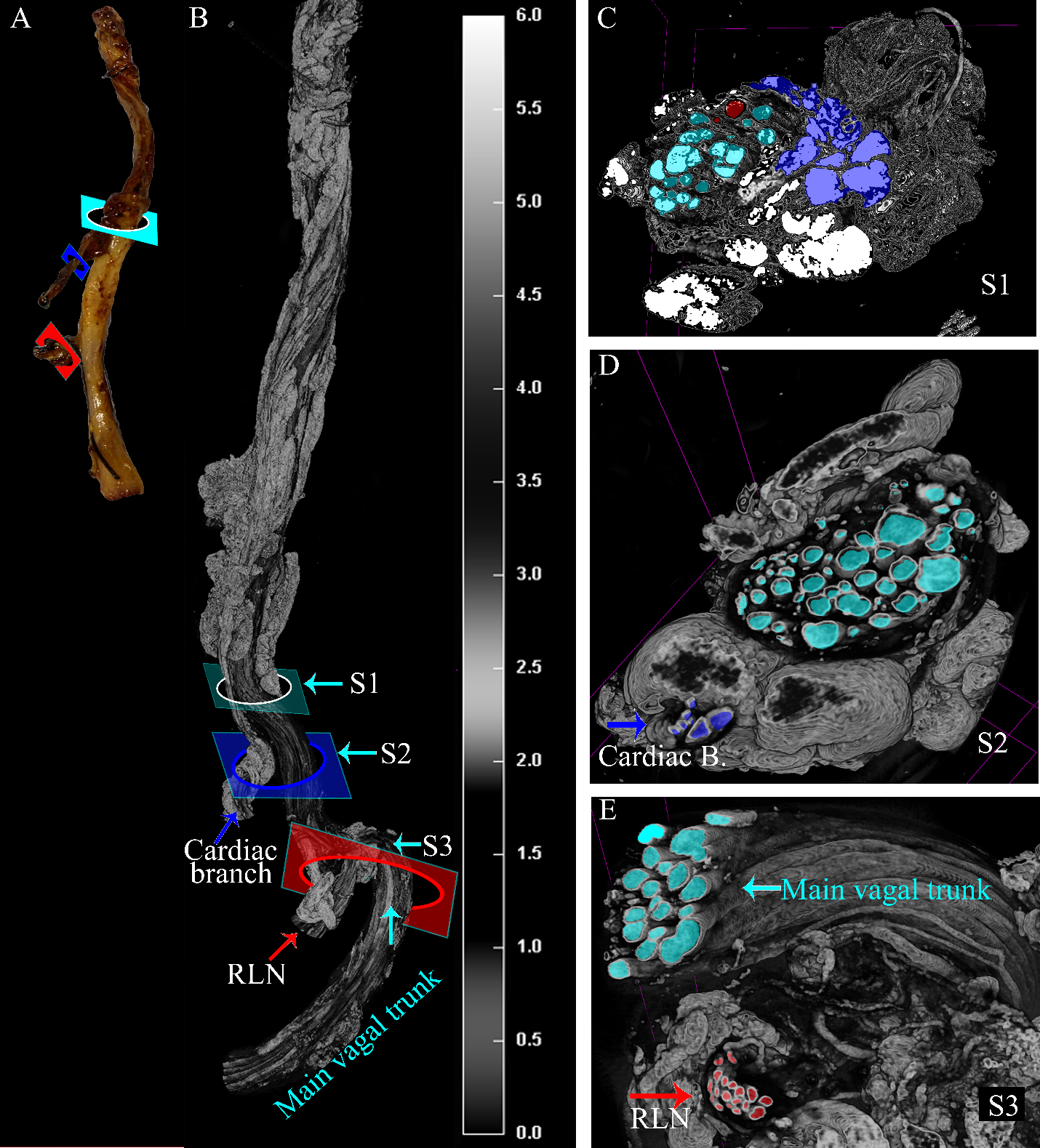

**Supplementary Fig S3.** **Cardiac and RLN branch**. A) Representative section of the nerve showing cardiac and RLN branch of the pig right vagus nerve. B) Micro-CT image of the vagus nerve showing cardiac and RLN branching from the main vagal trunk. (C-E). Fascicles corresponding to the RLN or Cardiac branch was manually annotated in the segmented micro-CT images with red or blue color respectively. Annotated fascicles of the vagal branches were traced to the main vagal trunk moving form rostral to caudal direction. C) Caudal section of the Vagus nerve showing fascicles that projected to the Cardiac branch (blue color) and the RLN ( Red color). D) Micro-CT image of the cardiac branch showing 7 fascicles at distance of 16 cm from the nodose ganglion. Micro-CT image of the Recurrent Laryngeal branch showing 18 fascicles at distance of 17 cm from the nodose ganglion.

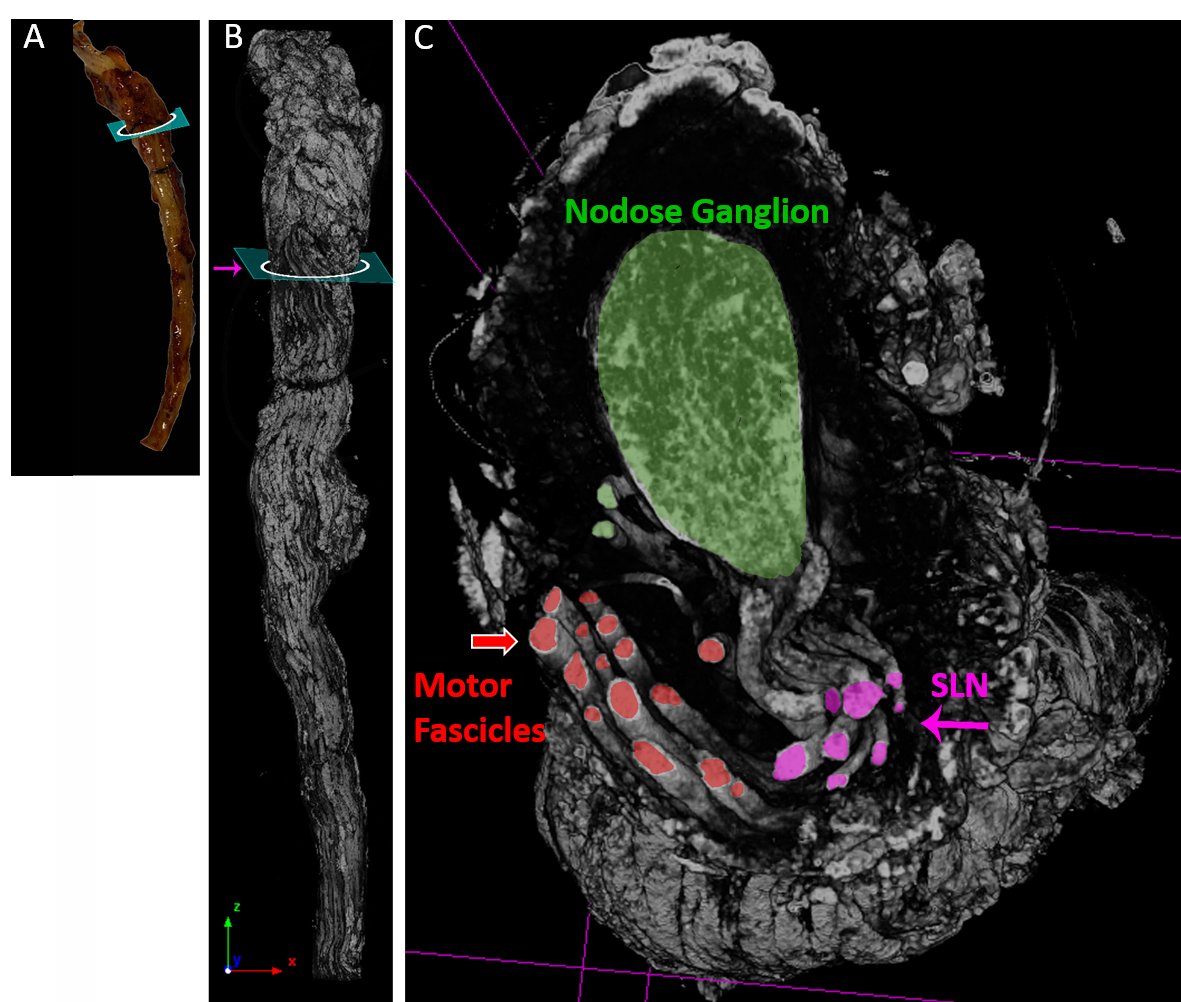

**Supplementary Fig S4. Nodose ganglion and the Superior Laryngeal branch.** A) Representative image of the right vagus that includes the nodose ganglion and the cervical region of the nerve. B) Micro-CT image of the corresponding section of the nerve shown in (A). Plane of the cut shows the region of the nerve where segmented micro-CT images was analyzed. C) Superior Laryngeal nerve extending from the nodose ganglion showing 8 fascicles was annotated in pink, motor fascicles in red color and the nodose ganglion containing the pseudo unipolar cells in green color.

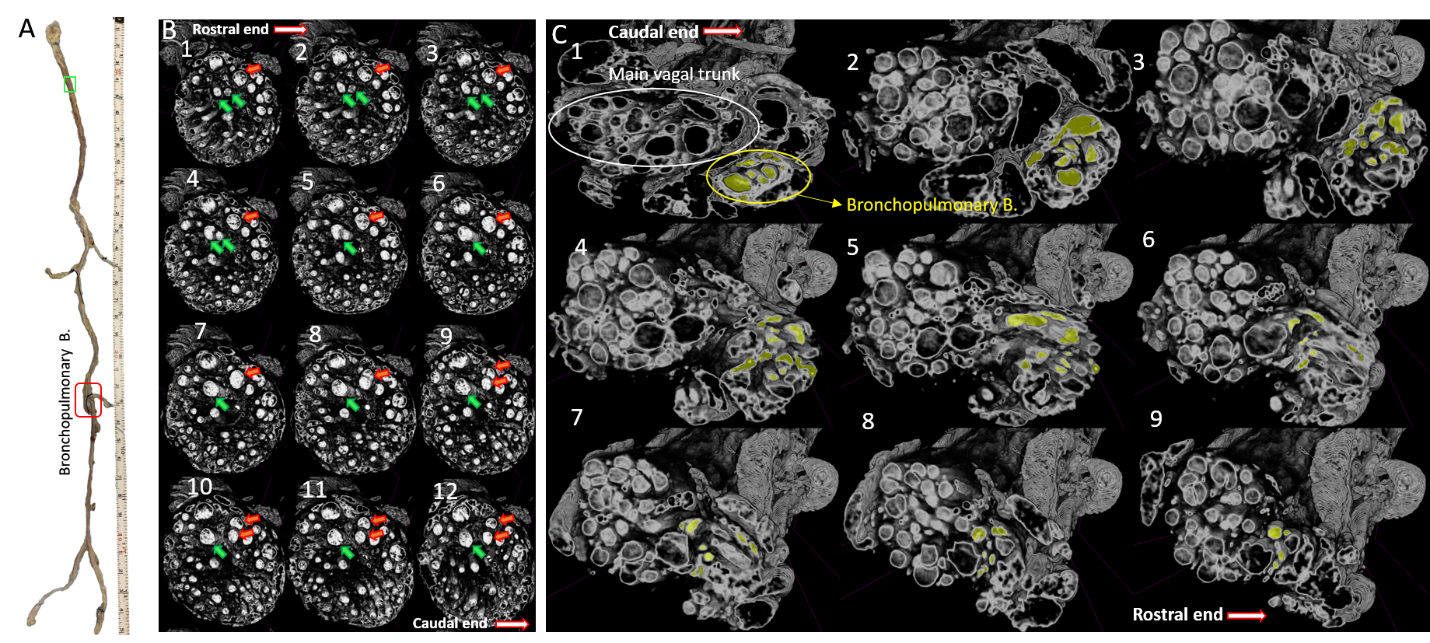

**Supplementary Fig S5. Fascicular branching and fascicular merging was observed throughout the length of the nerve**. A) Shown is a representative image of the nerve, indicating the region where the microCT images were taken at the cervical region (green box) and at the bronchopulmonary vagal branch point (red box) . B) Substantial fascicular branching and merging was observed over a small cervical region of the vagus nerve. Moving from the rostral to caudal direction, a single fascicle can be seen ( red arrow) splitting into two while the adjacent two fascicles ( green arrow), can been seen merging to form one single fascicle ( B1-B12). C) Constituent fascicle belonging to the bronchopulmonary branch was traced (yellow fascicles) to the main vagal trunk. Large number of fascicles from the bronchopulmonary branch was observed to fuse to from few number fascicles as the branch merged within the main vagal trunk (C1-C9).

a

b

c

d

e

f

g

h

i

j

k

3 cm

3.5 cm

4.5 cm

5.5 cm

6 cm

6.5 cm

7 cm

7.5 cm

8 cm

8.7 cm

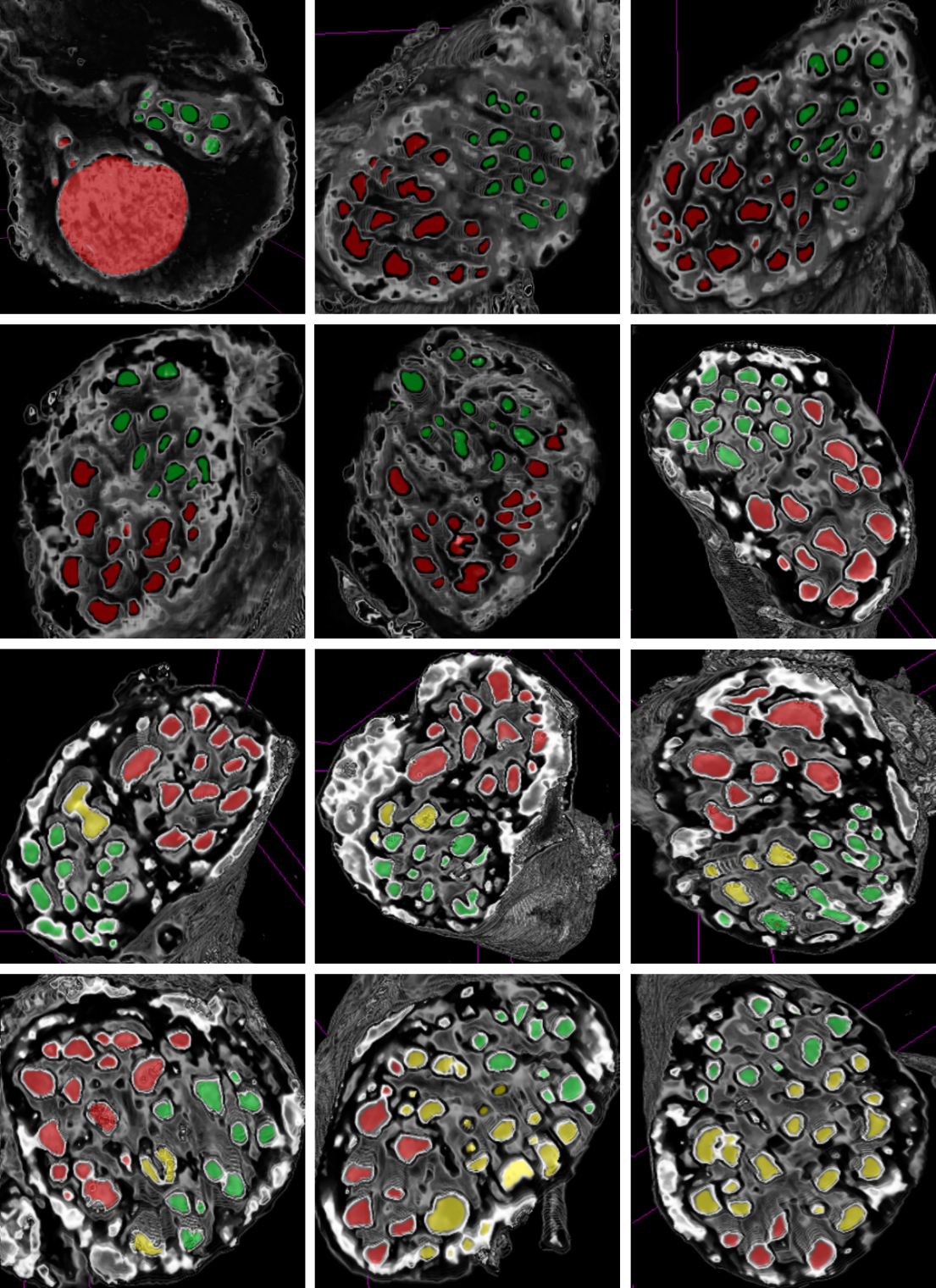

Afferent

Efferent

Mixed

l

5 cm

Nodose

**Figure S6: Micro-computed tomography imaging of an intact right vagus trunk, from Nodose ganglion to the upper thoracic level in a second animal**. Panels (a-l) show the trajectories of afferent and efferent fascicles in consecutive sections through the cervical vagus, at different levels, shown in cm from the rostral end of the sample.

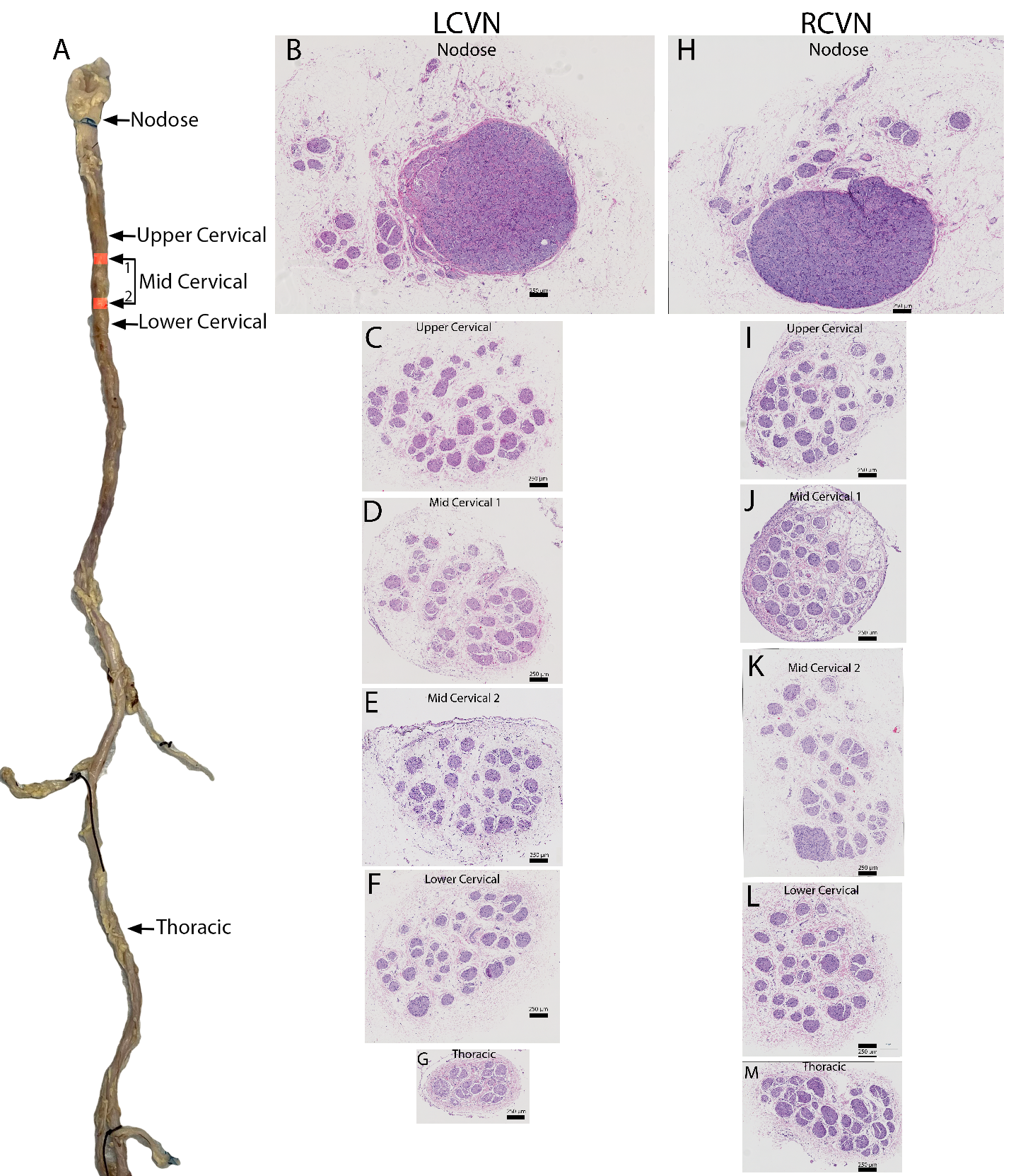
**Supplementary Fig S7. Representative H&E image of left and right vagus nerve at different levels of the nerve.** (A) Image of an intact pig vagus nerve explanted from nodose ganglion to the thoracic region. Cross sections of the nerve stained with Haematoxylin and Eosin (H&E) from both the left (B-G) and right (H-M) vagus taken at the Nodose (B,H), Upper cervical (C,I), Mid cervical (D,E,J & K), Lower cervical (F,L) and Thoracic (G,M). All scale bar 250 µm

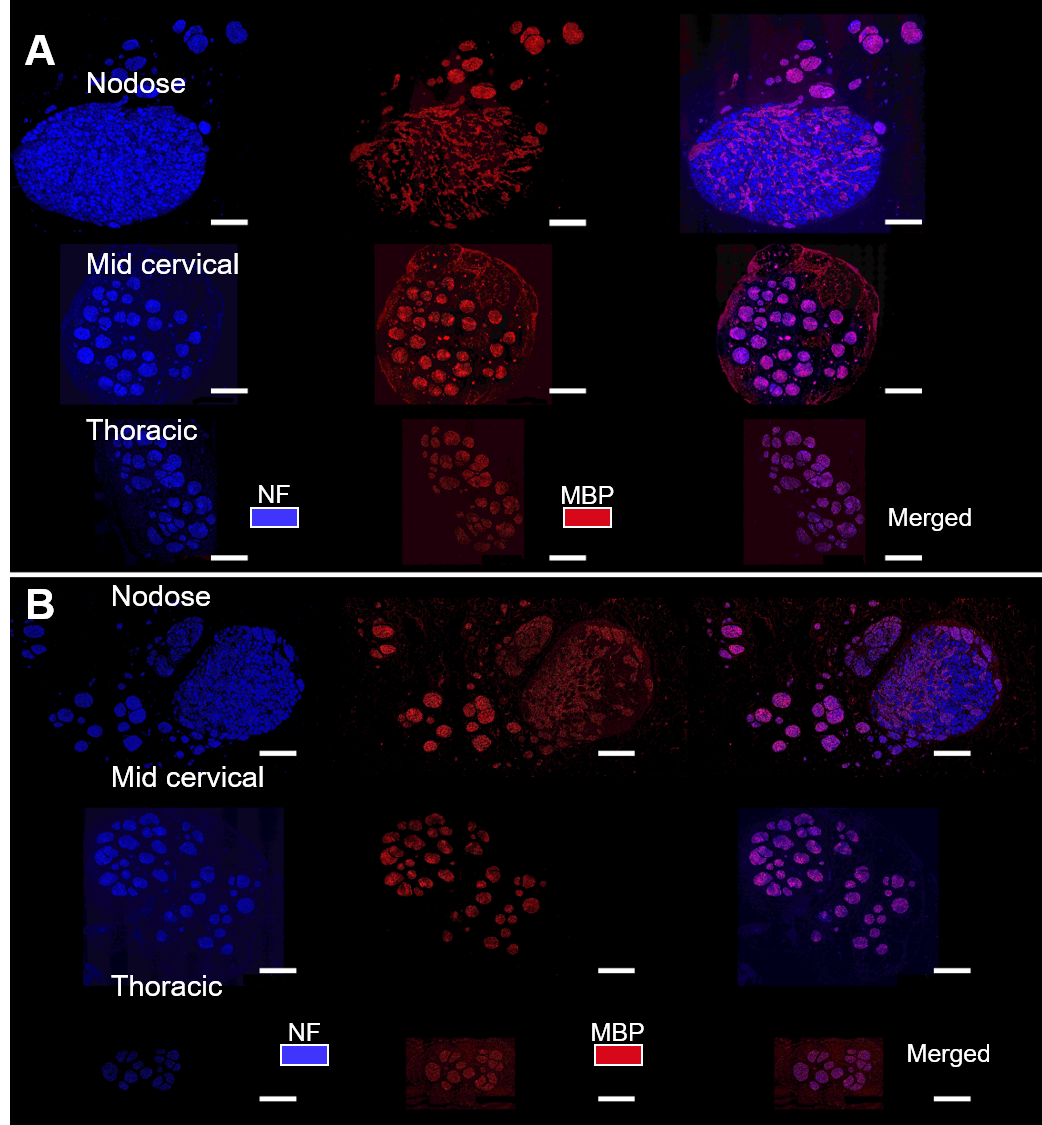

**Supplementary Fig S8. Representative IHC image of RCVN and LCVN.** Cross section of the right cervical vagus nerve (A) and left cervical vagus nerve (B) stained with Neurofilament and Myelin at the Nodose, Mid Cervical and Thoracic level. All scale bar 500 µm

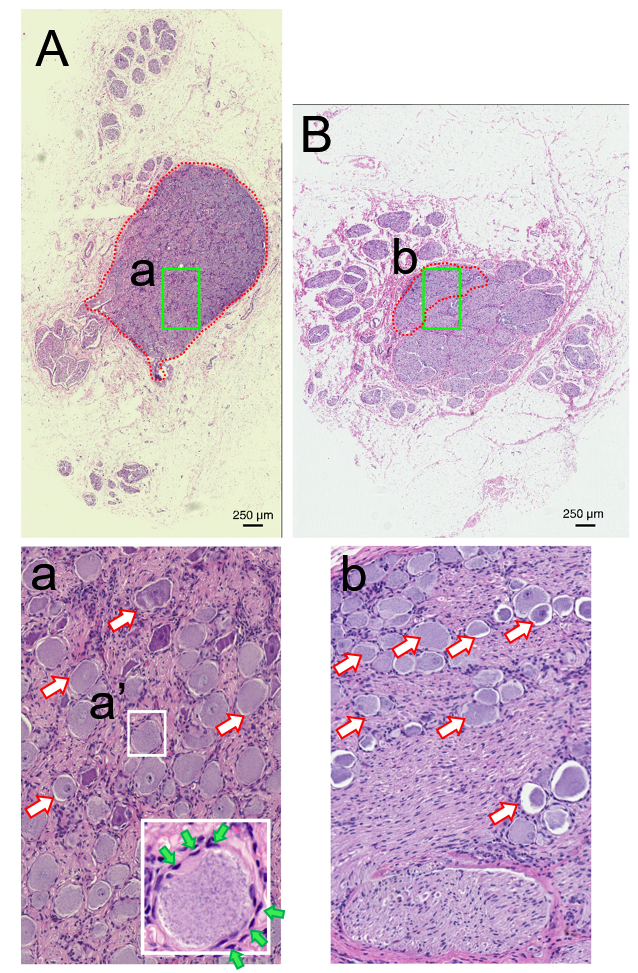

**Supplemental Figure S9: Aggregation of Sensory cell bodies in the Nodose ganglion:** H&E image of the Nodose ganglion from two different animals. Large group of pseudo unipolar cell bodies were clearly visible in ganglion of some nerves, while in other nerves the cell bodies were clustered to side of the nerve and or absent in others. (A). Pseudo unipolar cell bodies can be seen distributed evenly in the ganglion, marked by the red boundary. (B) Cell bodied can been seen clustered to one side of the ganglion and was absent in the remainder of the ganglion. Magnified images of the green insert are shown in a & b. Red arrow points to the pseudo unipolar cell bodies. Cell bodies were identified by their characteristic large shape and the surrounding satellite cells (green arrow) as shown in the insert (a’).

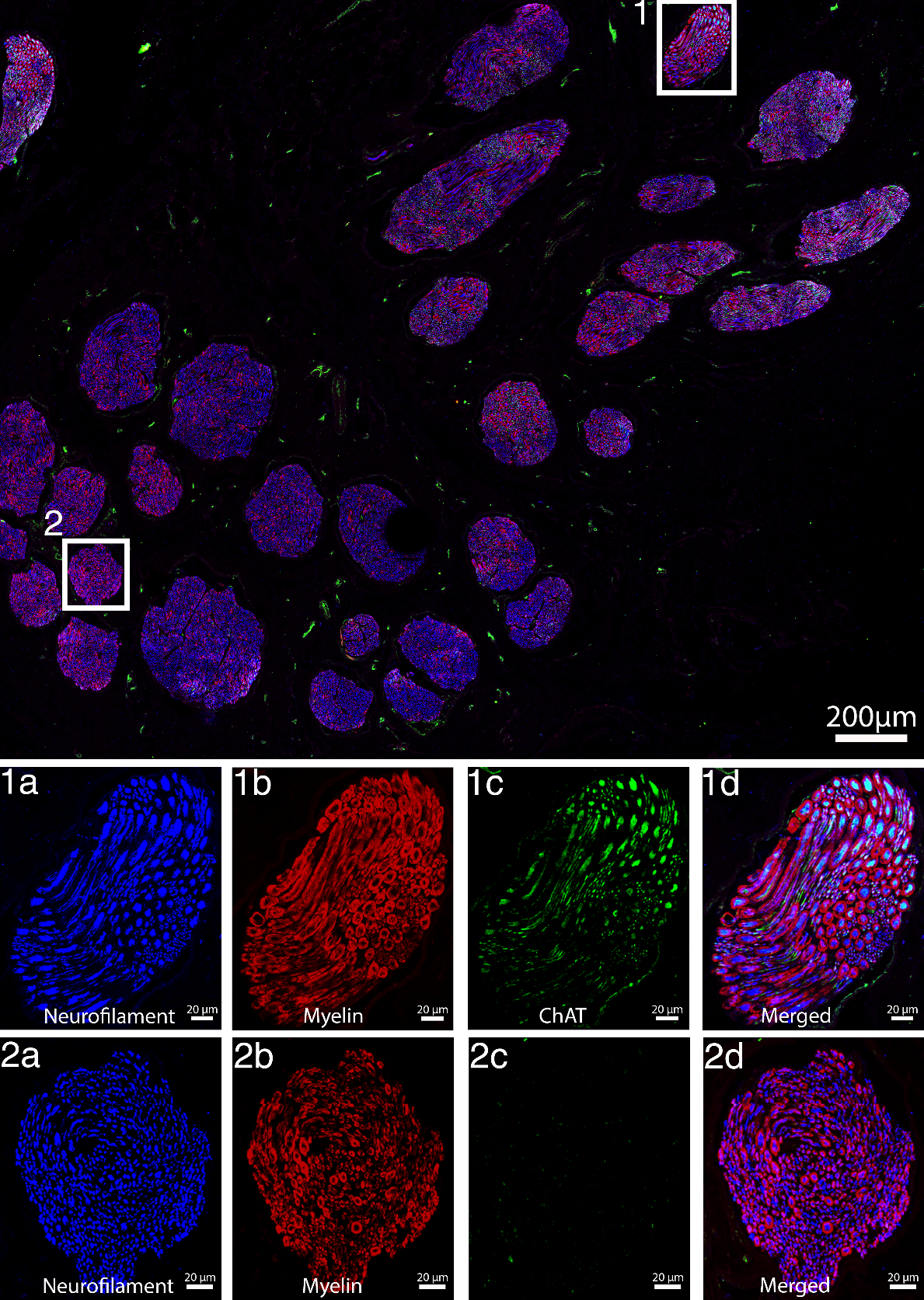

**Supplemental figure S10. Unimodal Expression Of Choline Acetyltransferase in the Cervical Vagus Nerve:** (A) Image shows cross section of the cervical vagus nerve stained with NF (green), Myelin basic protein (MBP) and ChAT (Choline acetyltransferase). Inserts (1 & 2) shows magnified image of two fascicles, taken at two different location of the nerve. Expression of the NF and MBP is uniform across all the fascicles in the nerve, however ChAT expression is confined to one segment of the nerve. Inserts 1a – 1d, shows an example image of a fascicle rich in ChAT+ fibers and insert 2a-2d, shows an example image of a fascicle poor in ChAT+ fibers.

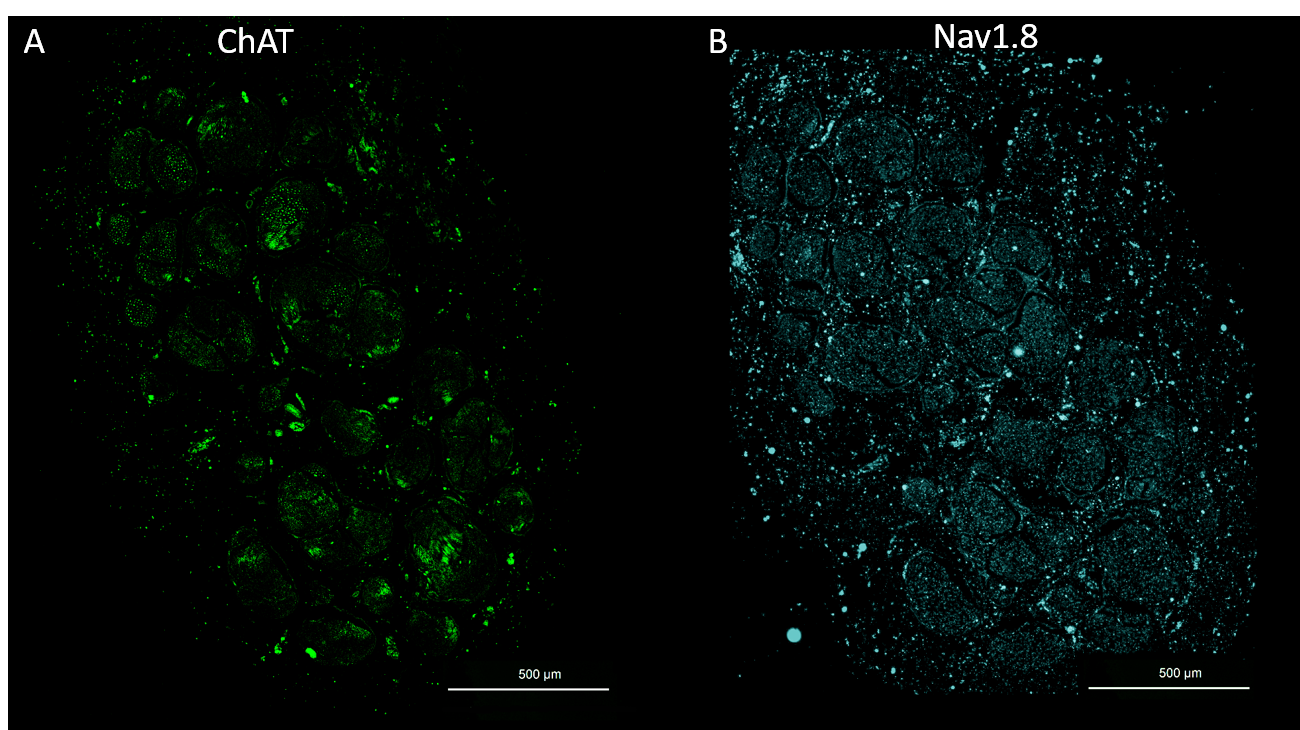

**Supplementary Figure S11.** **Bimodal distribution of the fibers is absent at the thoracic levels**. A) Immunofluorescence cross section image of the vagus nerve at the thoracic region stained with ChAT (A) and Nav1.8 (B) antibody. Expression of ChAT and Nav1.8 was found to be uniform across the nerve.

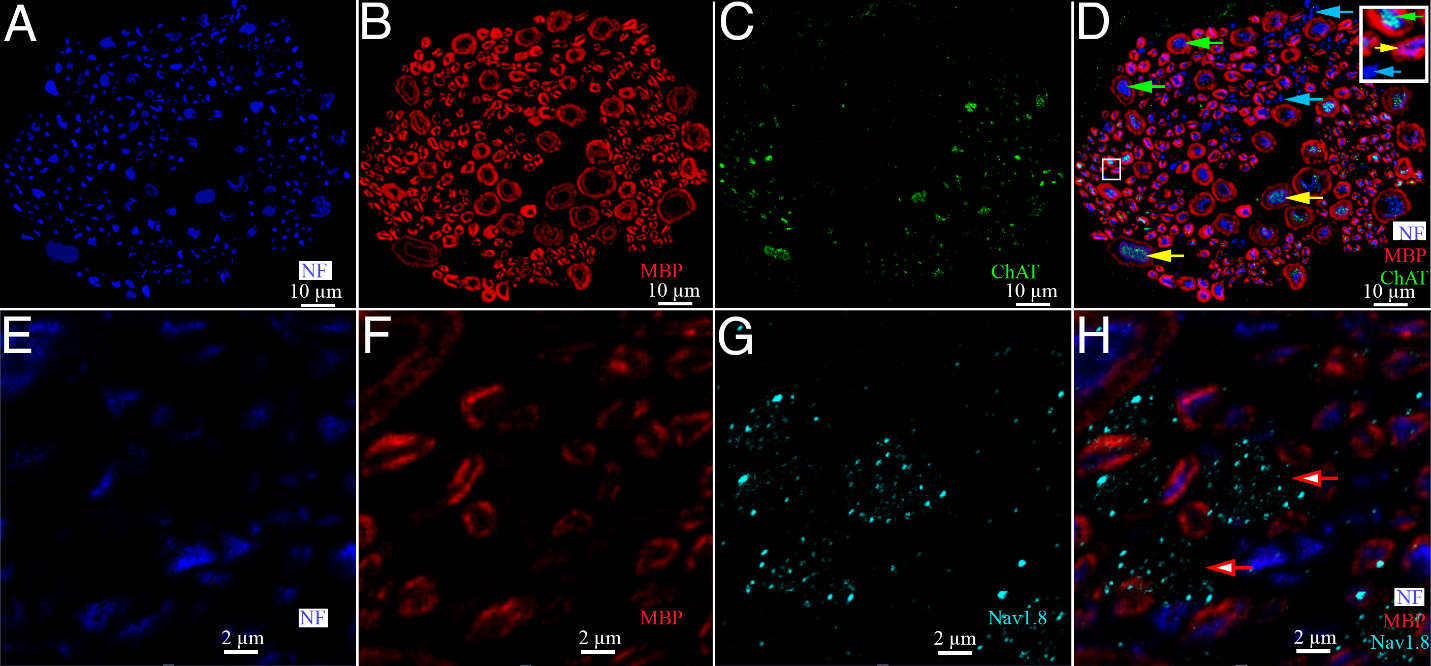

**Supplementary figure S12.** **Representative immunofluorescence image of a single fascicle showing expression of different fibers types**. Myelinated efferent fibers (MEff) that co-express Neurofilament (**A**), Myelin (**B**) and ChAT (**C**) are identified in yellow arrow in (**D**). Myelinated afferent fibers (MEff) that lack expression of ChAT are identified in green arrow in (**D**). Unmyelinated efferent fibers (UEff) that express only neurofilament are identified in blue arrow in (**D**). IHC co-localization of Neurofilament, Myelin and Nav1.8 was done in a separate section (**E-F**). Axons expressing Neurofilament (**E**), myelin (**F**) do not colocalize with Nav1.8 expression (**G**) as identified in the red arrow in (**H**).

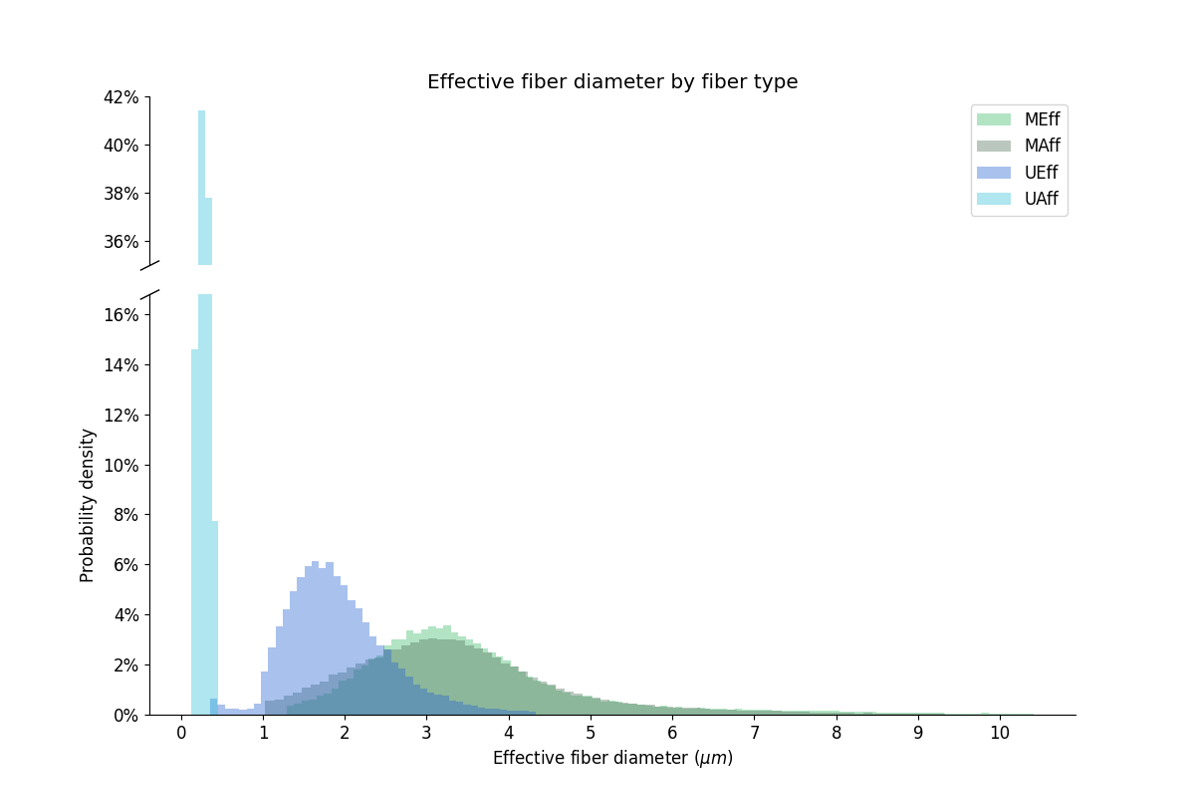

**Supplemental Figure S13. Distribution of effective diameters per fiber type.** Probability density plot of effective dimeter of MEff, MAff, UEff and UAff fibers imaged at the mid cervical region of the vagus nerve. The plot indicates distinct unimodal distributions for specific fiber types.

**Supplemental Figure S14. Distribution of effective diameters per fiber type.** (**A**) Spatial organization of fibers inside the nerve was measured in relation to the distance from the epineurium. Epinerium distances are arranged in bins of 200 um. (**A**-**a**) Illustration of bins (white lines) and enclosed fascicles (white patches) at distances 200-400 um (left) and 600-800 um (right) from the epinerium (red line). (**A, b-e)**
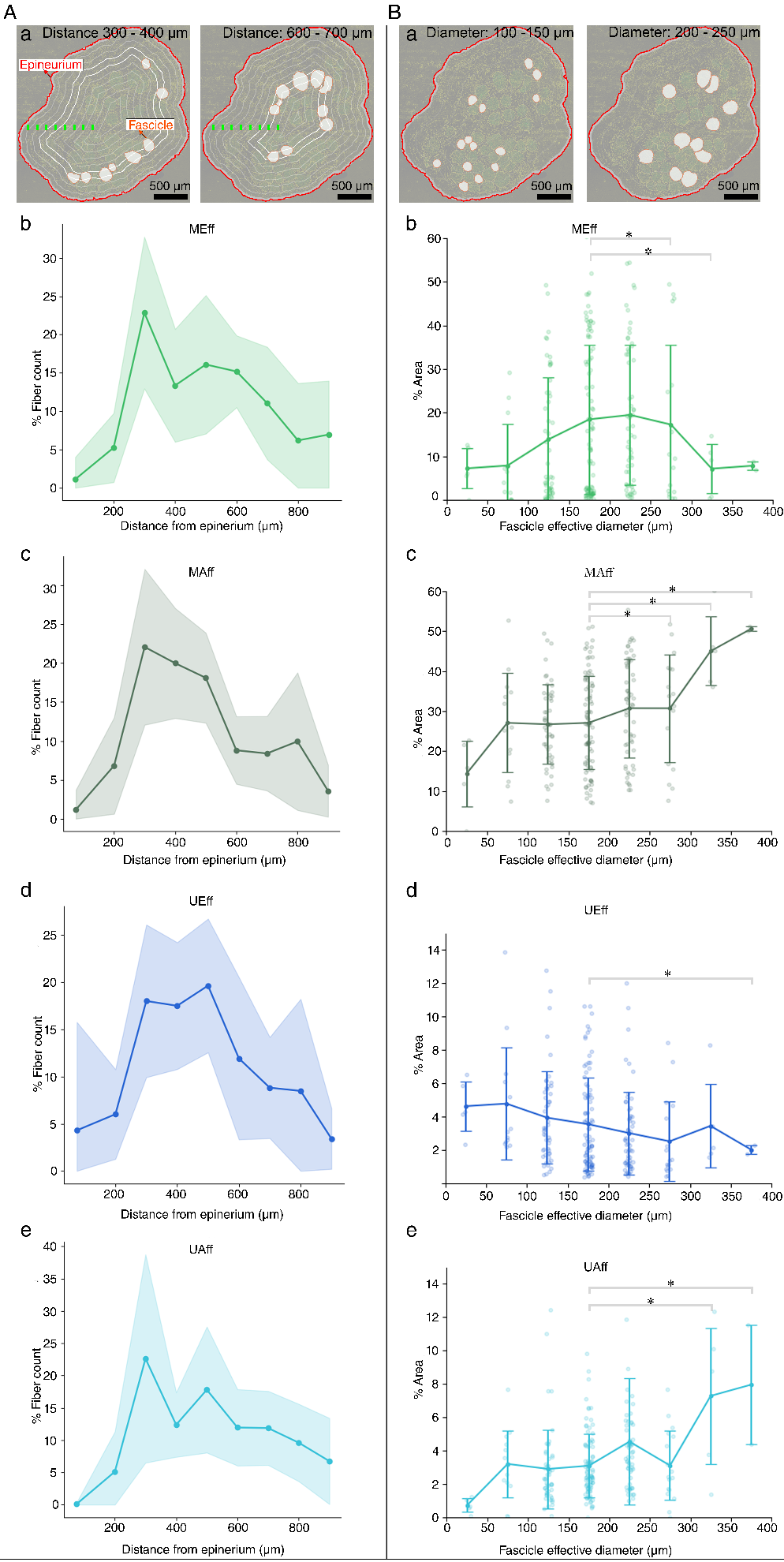
Percentage of MEff, MAff, UEff and UAff at different distances from the epinerium. The solid line shows the average percentage of fibers across nerves of different animals and the shaded region stretches one standard deviation away from the mean. (**B**) Fiber composition of the fascicle was measured in relation to the fascicle diameter by grouping fascicles into different bins according to the effective diameter. (a) Illustration of fascicles with effective diameter <200 µm(Left) and >200 µm (Right). We observe percentage area occupied by the efferent fibers (MEff and UEff) decreases with increasing fascicle diameter (**B, b & d**). Whereas percentage area occupied by the afferent fibers (MAff and UAff) increases with increasing fascicle diameter (**B, c & e**). Note* For each fascicle we captured the percentage area occupied by fiber type normalized by the fascicle area.

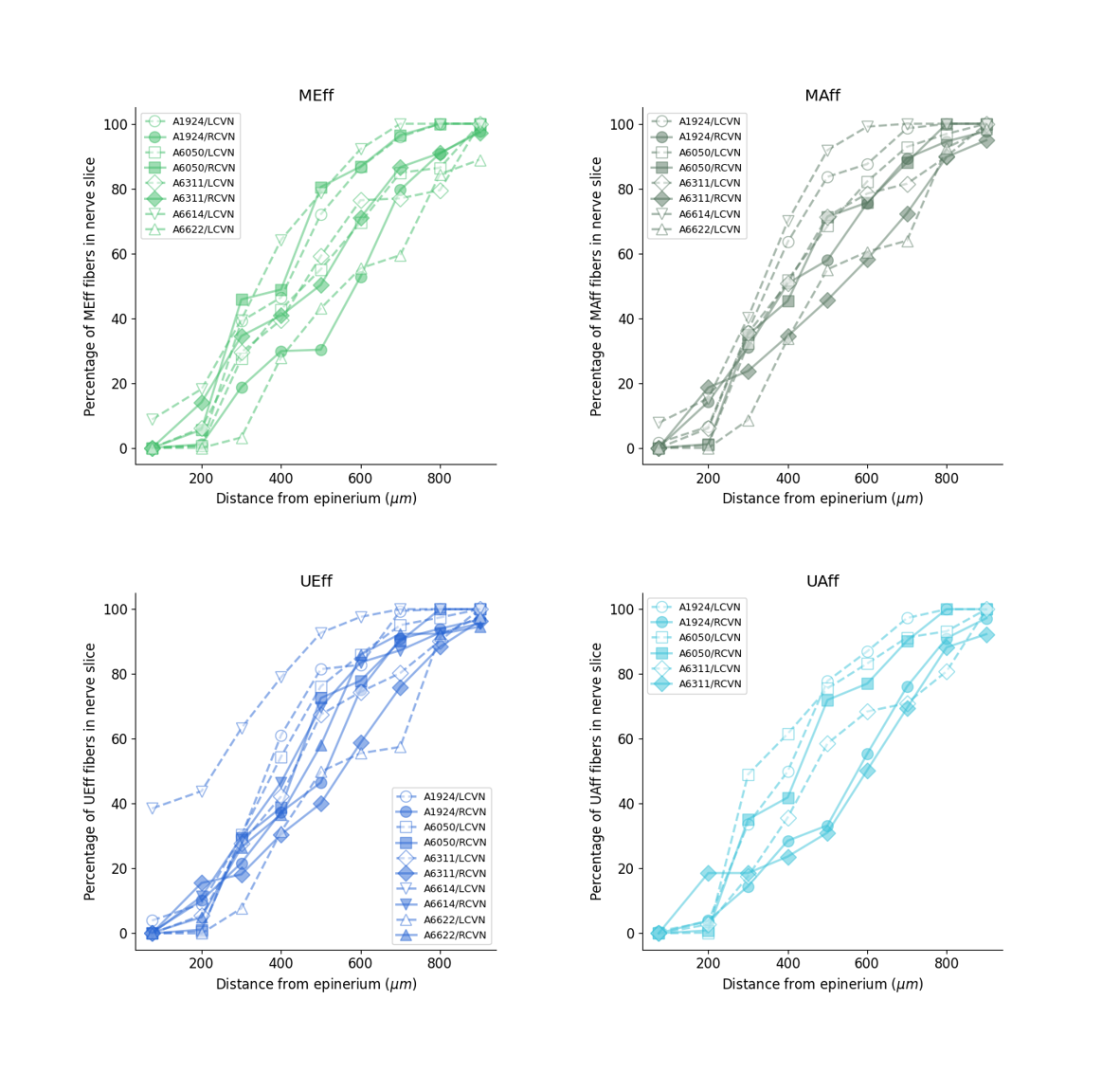

**Supplementary figure S15. Cumulative distribution of fiber types as the function of epineurium distance.** Fibers are counted at bins of 200 µm, as in S11. We show the cumulative distribution of the percentage of fibers at different epineurium distances separately for both LCVN and RCVN for each animal at mid cervical level.

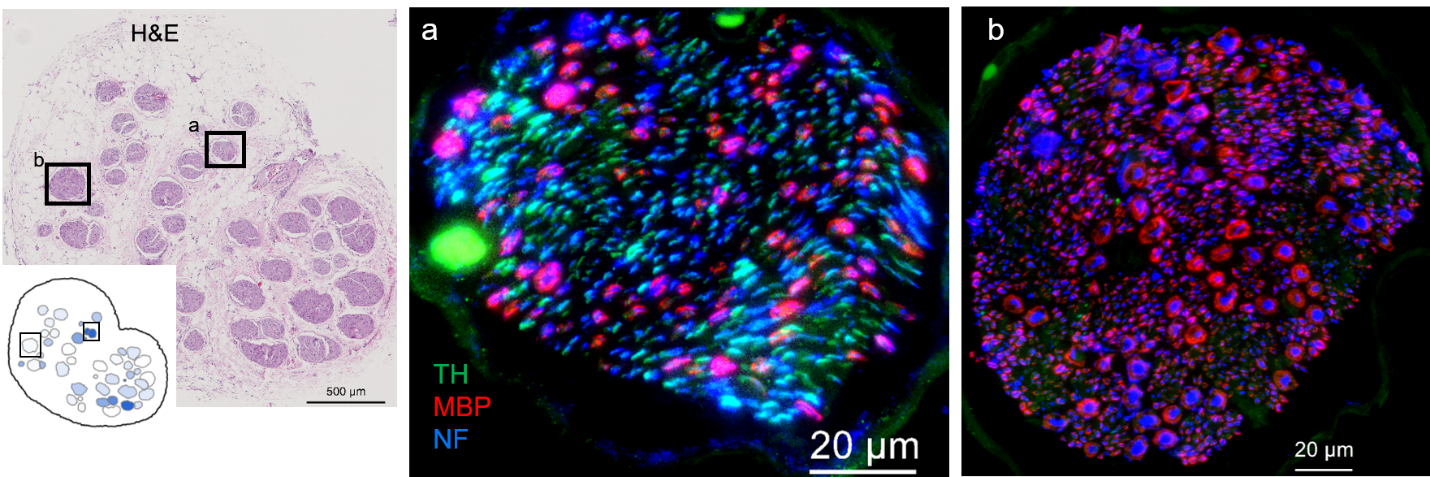

**Suppl. Figure S16. Tyrosine hydroxylase (TH)-immunoreactive fibers in the pig cervical vagus nerve**. (A) Cross section of the left cervical vagus nerve stained with Hematoxylin and Eosin (H&E). The color map shows the distribution of the UEff fibers in the same nerve at the cervical level. Darker colors denote a higher proportion of presence of a TH+ fibers in the fascicle. Panel (a & b) shows a magnified IHC image of two fascicles, taken at two different location of the nerve stained with NF, MBP and TH. Expression of the NF and MBP is uniform across all the fascicles in the nerve, however TH expression is confined to only to certain fascicles of the nerve. Panel (a) shows example image of the fascicle rich in TH+ fibers and panel (b) shows example image of the fascicle poor in TH+ fibers.

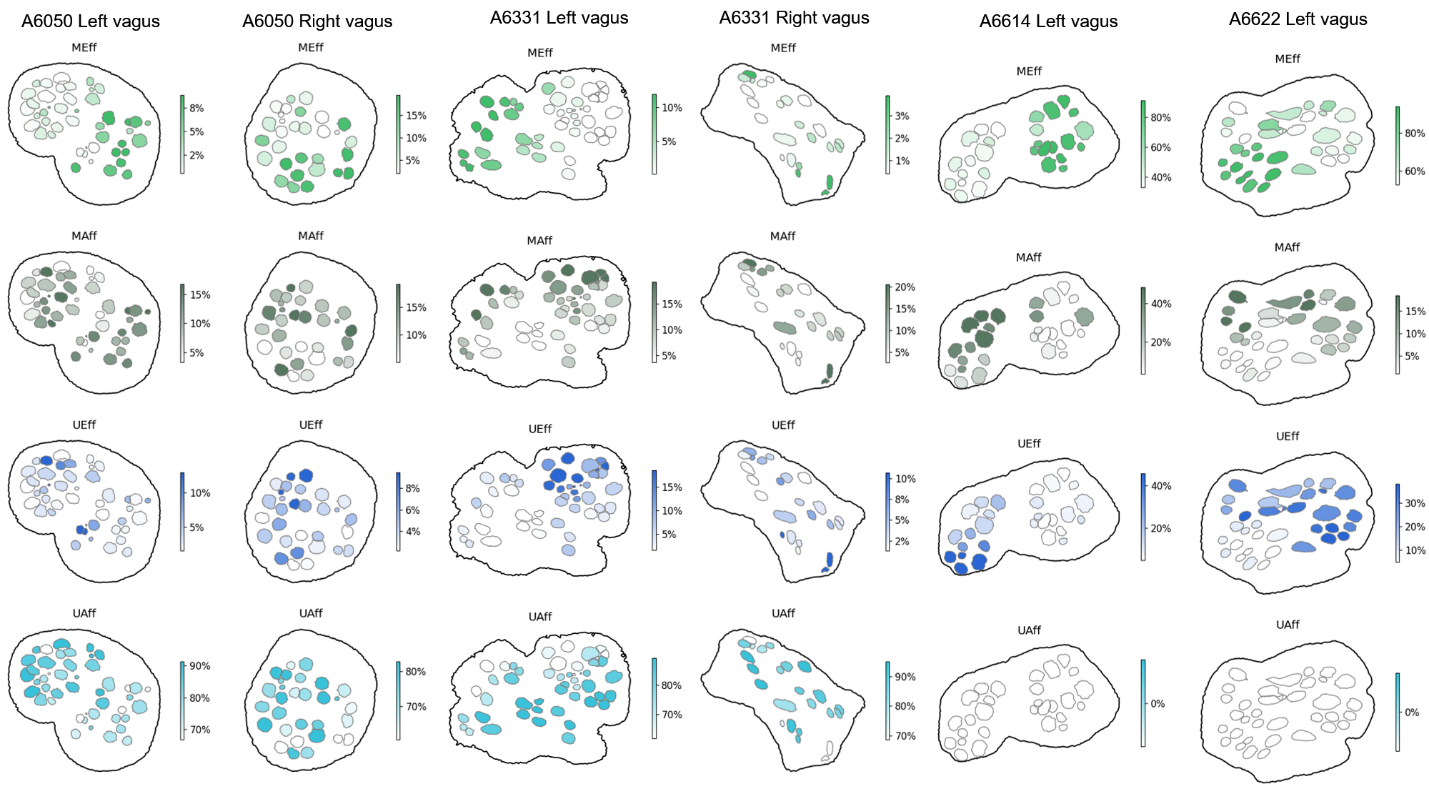
Supplemental Figure S17 : Percentage of different fiber types per fascicle at the mid cervical vagus nerve in different animals : The color map shows the distribution of the MEff, MAff, UEff and UAff fibers at the mid-cervical region of the vagus nerve in different animals. Darker colors denote a higher proportion of presence of a given fiber type. For each fascicle, percentages across fiber types amounts to 100%. Animals A6614 and A6622 were omitted from Nav1.8 staining , hence UAff fibers were not quantified for these animals.

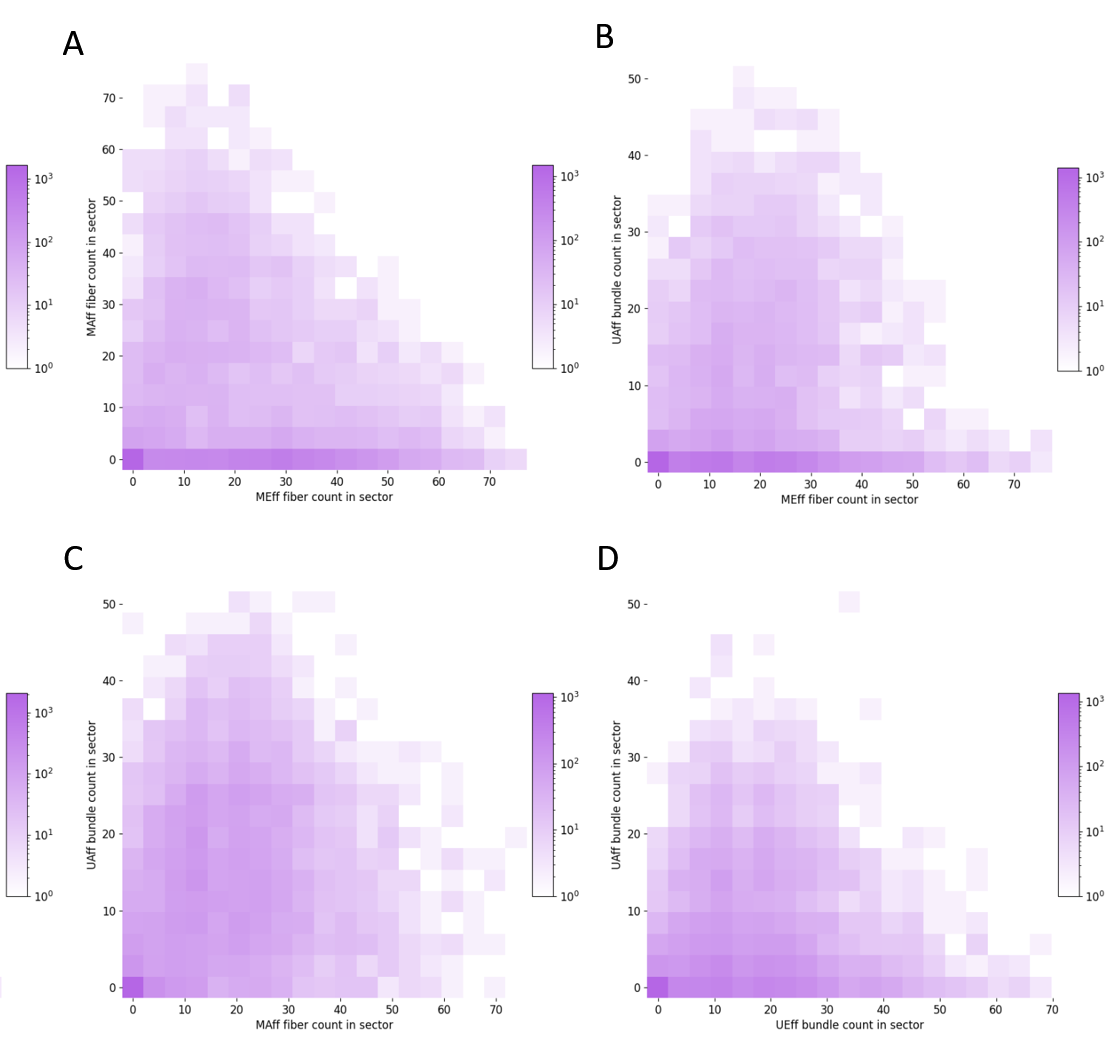

**Supp Figure S18. Spatial interaction of the fibers.** Plot shows within-fascicle spatial interaction of fiber type counts in spatial sectors of 30 um width. Fascicles were divided into a non-overlapping grids of sectors and counts of MEff, MAff, UEff and UAff inside these sectors were contrasted against each other across all fascicles at the mid-cervical level. Spatial interaction maps show the distribution of fiber type compositions when contrasting MEff with MAff counts (A), MEff with UAff counts (B) MAff with Uaff counts (C) and UEff with UAff counts (D). Note* Counts of UEff and UAff reflect bundle counts and counts of MAff and MEff reflect individual axon counts within the 30 um spatial sectors.

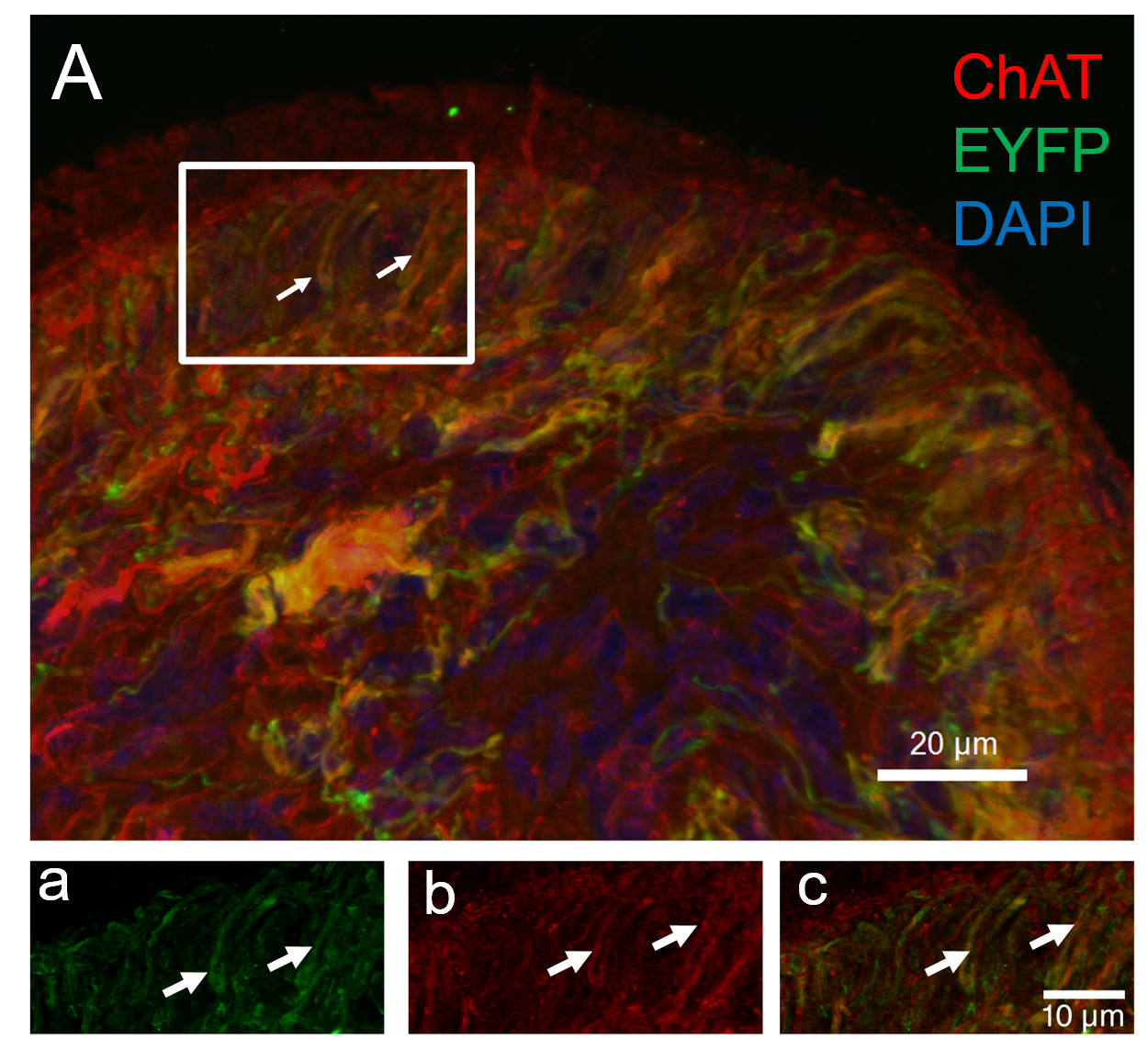

**Supplemental Figure S19**: Vagus nerve of ChAT (BAC)-eYFP transgenic mice, labelled with ChAT antibody. A) Expression of ChAT in vagus nerve of the transgenic mice colocalize with the innate expression of eYFP expressing axons. Bottom panels shows magnified image of the insert (a) eYFP, (b) ChAT and (c) merged image

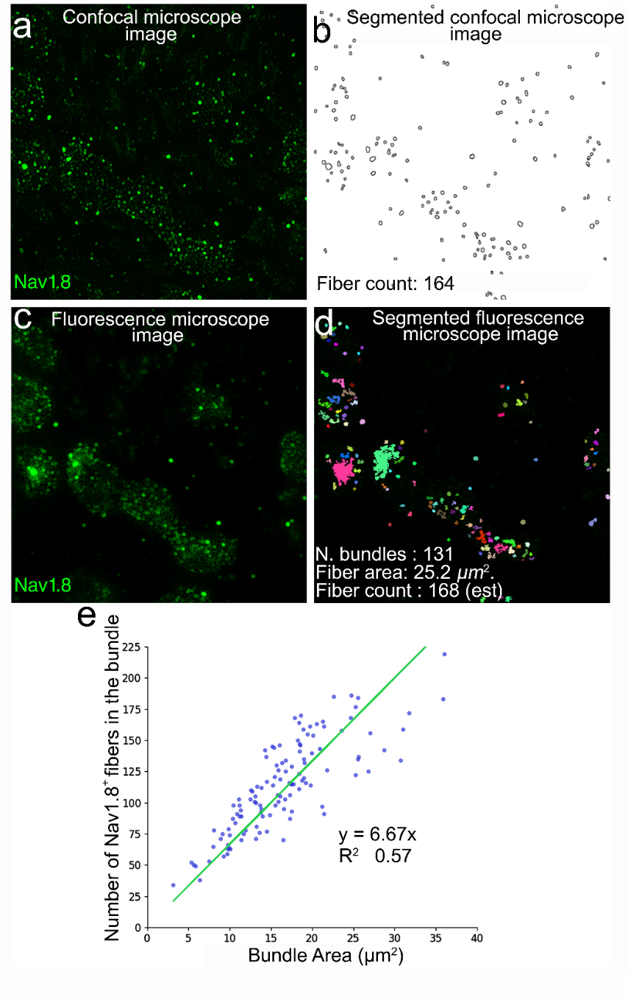

**Supplemental Figure S20. Detection and quantification of SCN10A (Nav1.8). (**a) Airyscan equipped super high resolution confocal image, showing a region of fascicle in mid cervical vagus nerve stained with Nav1.8 where Individual UAff fibers can be distinguished clearly. (b) We quantified Nav1.8 positive fibers using manual annotation. Automated cell identification process implemented in ImageJ was used to verify the counts done using manual annotations. Shown is the contour of individual fibers identified using the automated process. (c) Same region of the nerve as **(**a) imaged in a traditional wild field fluorescence microscope. (d) On this coarser, traditional fluorescence microscope image single UAff fibers cannot be distinguished, however, the area of UAff bundles can be approximated. Accordingly, we quantified the area occupied by afferent C-fibers by setting an appropriate threshold and counting the positive pixels. (e) To link the quantified bundle area to fiber counts, we imaged individual bundles in super high resolution, measured their area and manually counted fibers in each bundle. Each dot in the plot represents a bundle. We perform linear regression to correlate bundle area to the number of enclosed C-fibers.

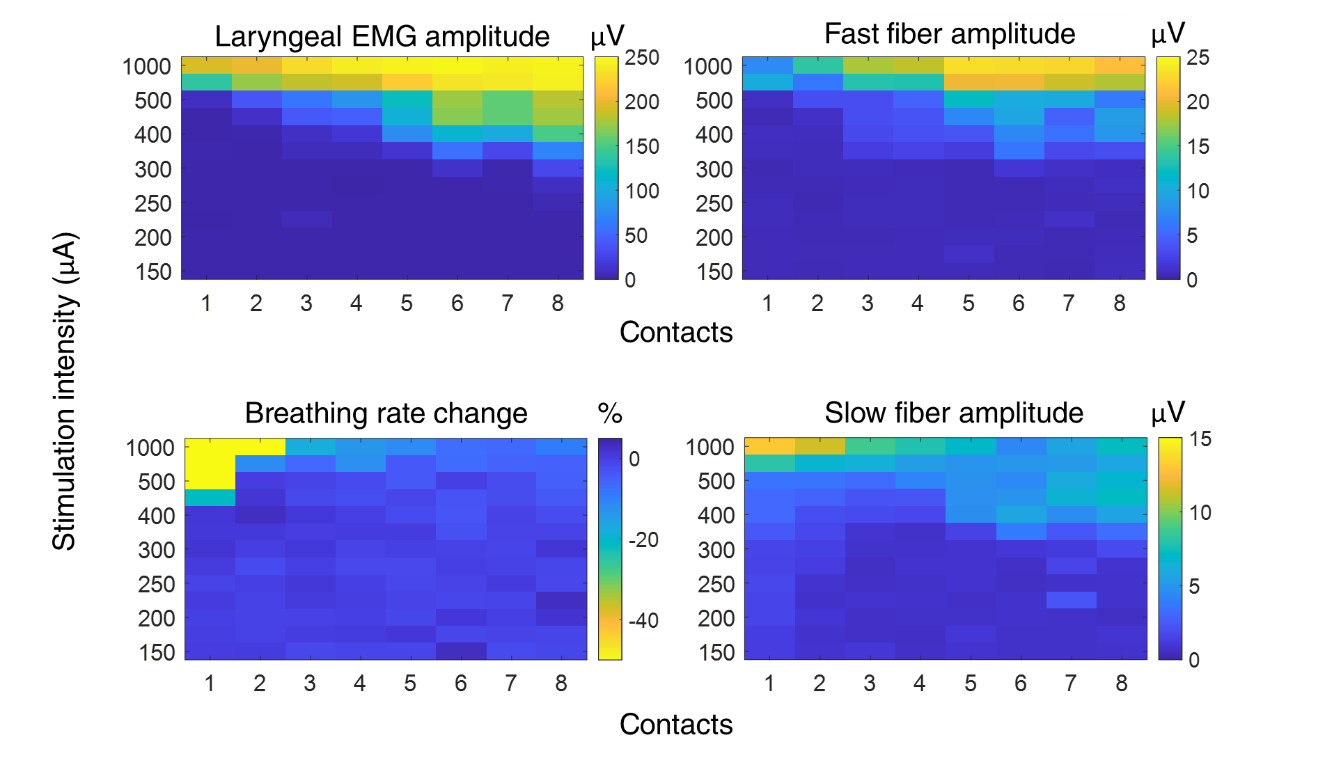

**Supplementary Figure S21. Recruitment maps of some physiological responses and the fibers mediating them, from different contacts of an epineurial nerve cuff placed on the cervical vagus of a swine.** (A) Amplitude of laryngeal EMG represented by color, at each of the 8 contacts of the electrode device (abscissa), for different stimulus intensities (ordinate). (B) Corresponding amplitude of the fast fiber response to the same stimuli. (C) Percent change in breathing rate for the same stimuli. (D) Corresponding amplitude of the slow fiber response to the same stimuli.

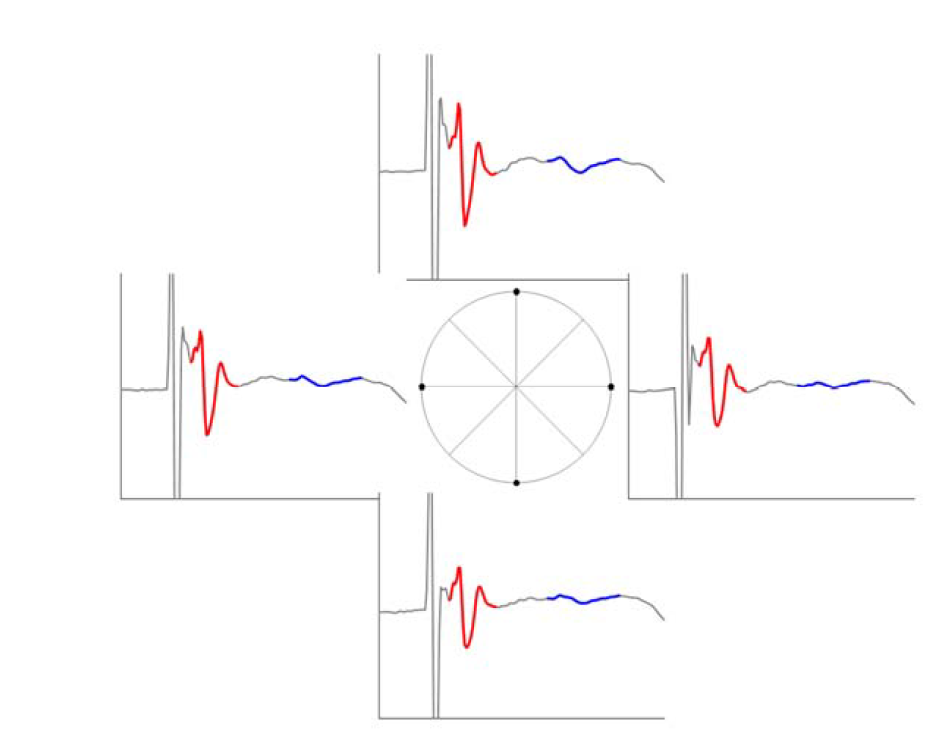

**Suppl. Figure S22**. **Stimulus evoked compound action potential (eCAP) traces through different contacts of the recording cuff electrode, in response to stimuli delivered to one of the contacts of the stimulating cuff electrode.** The overall shape and amplitude of eCAP responses is similar across all recording contacts. For that reason, the eCAP response in all analyses was calculated from the average trace across all 8 contacts of the recording cuff electrode.

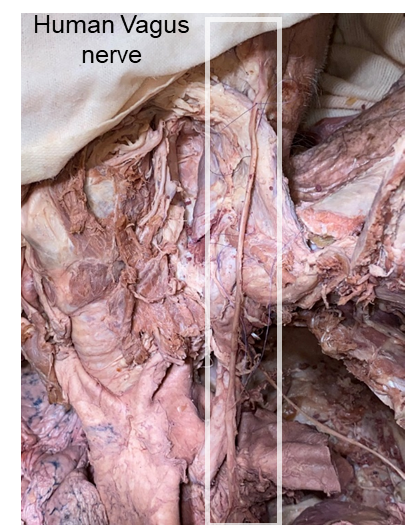

**Supplementary Figure S23**: Surgical view of human left vagus nerve at the cervical and thoracic inlet region.

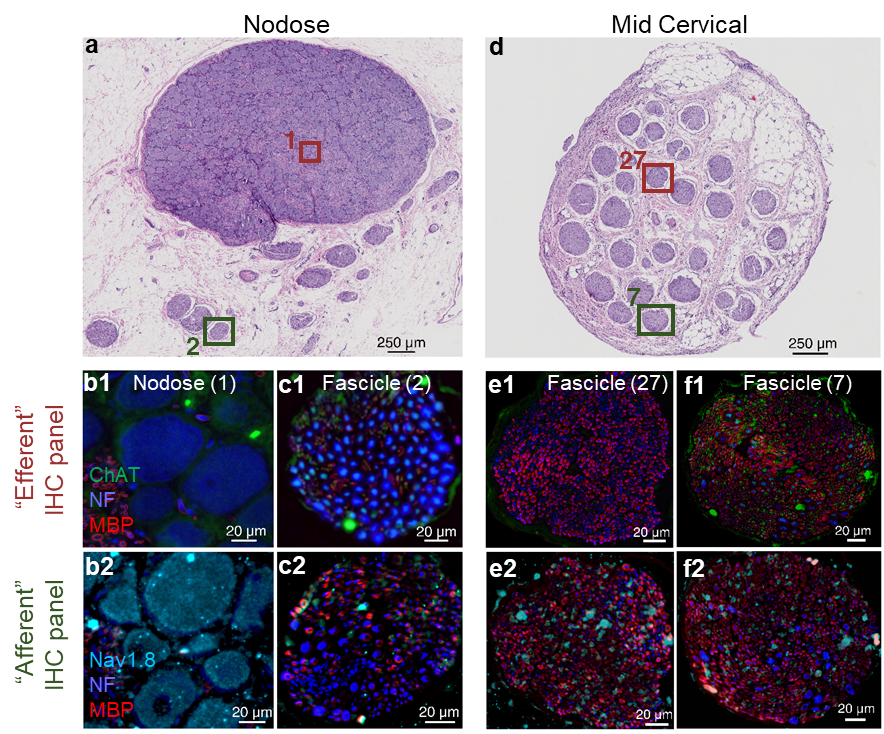

**Supplementary figure S24**: Imaging of afferent and efferent fibers in primarily afferent and primarily efferent fascicles. (a) H&E section at the level of the nodose ganglion. An area inside the ganglion (1) and a primarily efferent fascicle (2) are selected. (b1) Nodose area stained with an “efferent” panel of antibodies for immunohistochemistry (IHC), consisting of anti-neurofilament antibody (NF, blue) that labels nerve fibers, anti-myelin basic protein antibody (MBP, red) that labels myelin, and choline-acetyl-transferase (ChAT, green) that labels efferent, cholinergic, fibers. (b2) The same nodose area stained with an “afferent” IHC panel, consisting of NF, MBP and anti-Nav1.8 antibody (cyan) that labels afferent fibers. (c1) Fascicle (2) stained with an efferent IHC panel. (c2) Fascicle (2) stained with an afferent IHC panel. (d) H&E section at the mid-cervical level. A primarily afferent (27) and a primarily efferent fascicle (7) are selected. The 2 fascicles are stained with an efferent (e1 and f1, respectively) and an afferent IHC panel (e2 and f2, respectively).

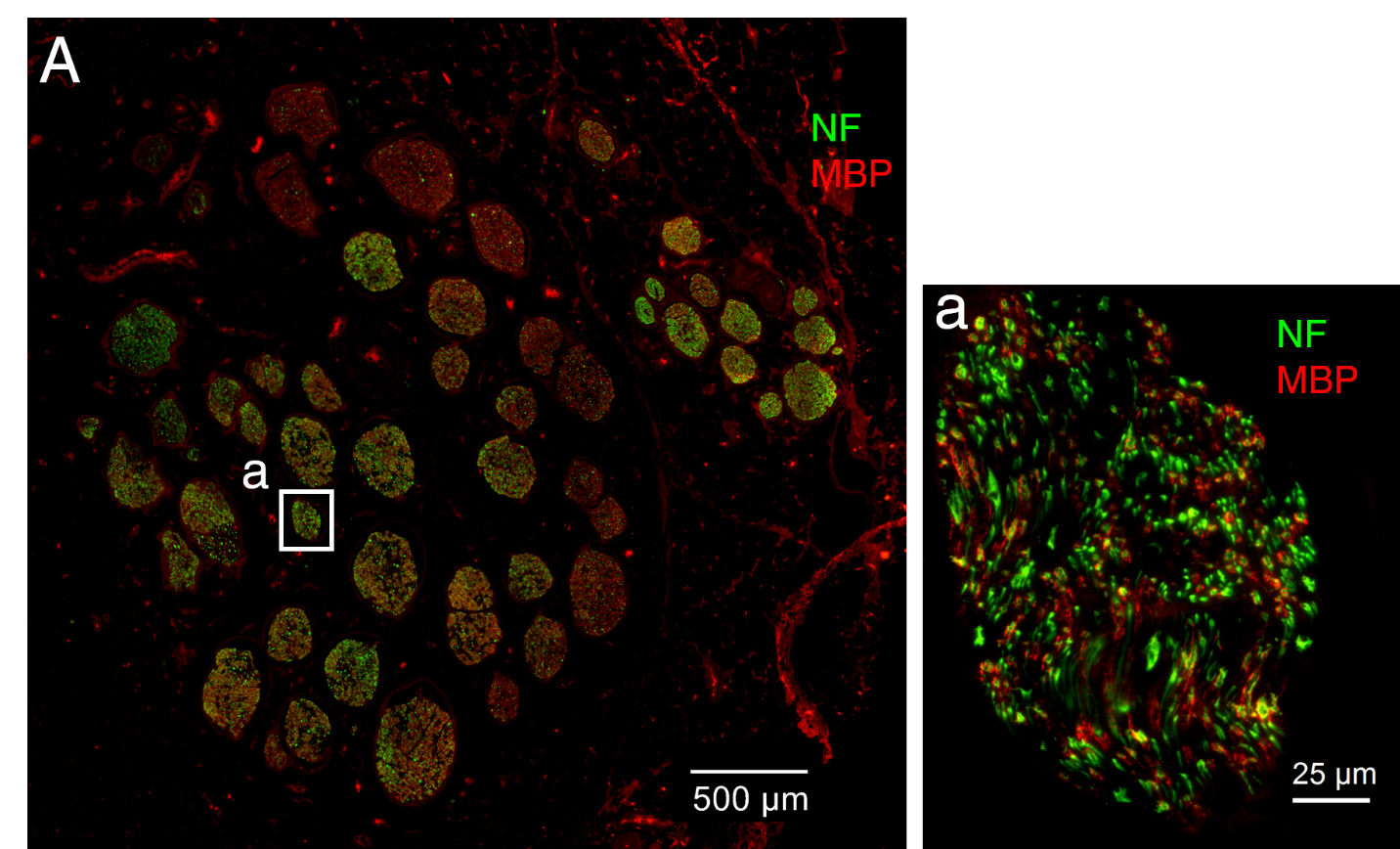

**Supplementary Figure S25: Integrity of the vagus nerve after chronic cuff placement**: (A) Cross sections of the cervical vagus nerve were taken below the chronic cuff and stained for NF and MBP (Animal k). Panel (a) shows a single fascicle imaged at 100x with expression of the NF and MBP intact in the axons and no visible damage to the nerve observed.

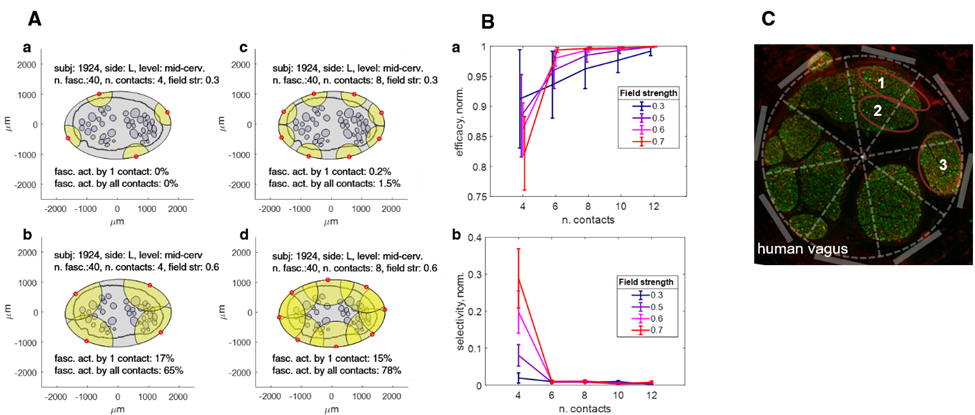

**Supplementary figure S26:** Cuff design with the 8 contacts compatible with the human vagus nerve to achieve spatially selective stimulation. Shown here is the 8-contact electrode design superimposed with the fascicular anatomy of a human cervical vagus nerve. 3 fascicles (1, 2, 3) that eventually give rise to the recurrent laryngeal nerve are highlighted.

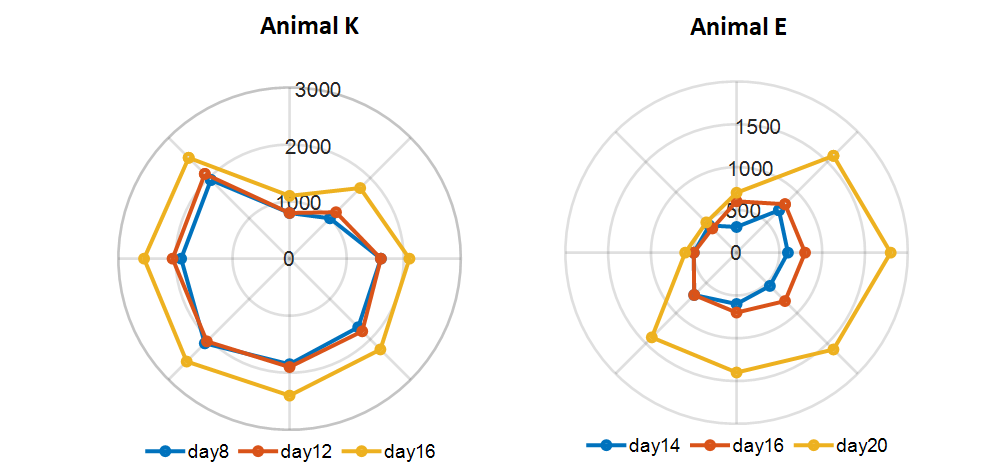

**Supplementary figure S27:** coughing threshold across days in two animals. The selectivity of the VNS is stable across days, while the coughing threshold increases with implantation days for both animal K and animal E.

| **Branch** | **Distance from nodose (cm)** | |
| --- | --- | --- |
|  | Domestic | Yucatan |
| Superior Laryngeal | N/A | N/A |
| Cardiac | 13 | 14 |
| Recurrent Laryngeal | 15 | 19 |
| Pulmonary | 24 | 24 |

**Suppl. Table 2: Summary distance from the nodose ganglion in each of the identified vagal branch**

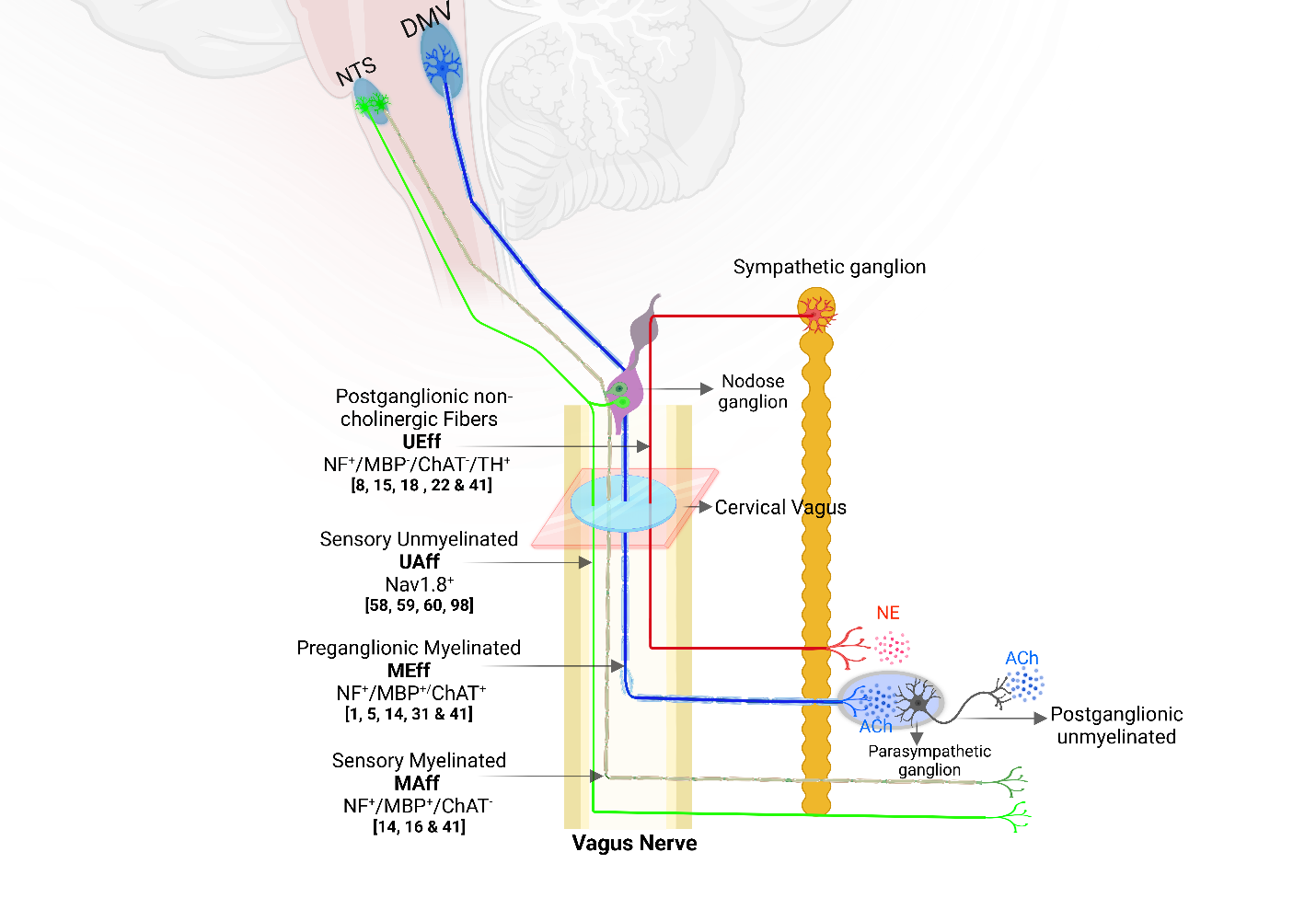

**Supplementary figure S28:** Schematic showing afferent and efferent fibers of the vagus nerve. The Preganglionic myelinated efferent fibers (blue axons) emerge from the Dorsal motor nucleus (DMV) to terminate in the parasympathetic ganglion. Sensory unmyelinated (bright green axons) and myelinated sensory (light green axons) afferent fibers terminate at the nucleus tractus solitaries (NTS) with their pseudounipolar cells bodies in the nodose ganglion. Postganglionic non-cholinergic (ChAT^-^) fibers that are also TH^+^ (Red axons) in the vagus nerve emerge from the sympathetic chain.

| **Type** | **Morphology** | **Functions** | **NF** | **MBP** | **ChAT** | **Nav1.8** | **Diam. criteria** |
| --- | --- | --- | --- | --- | --- | --- | --- |
| **Aα** | Large, myelin. | Efferent; somatic motor (laryngeal muscles) | + | + | + | - | >10 μm |
| **Αβ** | Large, myelin. | Afferent; somatic sensory (laryngeal and pharyngeal mucosa, cough reflex) | + | + | + | - | >5 μm |
| **Aδ** | Small, myelin. | Afferent; visceral sensory, nociceptors, mechanoreceptors (lung, heart, vessels, GI tract) | + | + | - | - | <3 μm |
| **B** | Small, myelin. | Efferent; visceral motor (heart, bronchi, vessels, GI tract) | + | + | + | - | <3 μm |
| **C** | Small, unmyelin. | Efferent; visceral motor (heart, bronchi, vessels, GI tract) | + | - | - | - | NA |
| **C** | Small, unmyelin. | Afferent; visceral sensory, nociceptors, chemoreceptors (lung, GI tract) | - | - | - | + | <1 μm |

**Suppl. Table 3: Detailed criteria used for classification of single fibers in distinct functional types.**

All nerve fibers, except small unmyelinated afferents, are positive for neurofilament (NF).

Myelinated fibers are positive for myelin basic protein (MBP) and can be afferent or efferent.

Myelinated efferents are positive for NF, MBP and ChAT, and can be large (Αα-type, >10 μm in diameter, typically somatic motor fibers to laryngeal muscles), or small (Β-type, <3 μm in diameter, typically parasympathetic motor fibers to visceral organs) [1,5, 15, 31 & 41].

Myelinated afferents are positive for NF, BMP but negative for ChAT, and can be large (Aβ-type, >5 μm in diameter, typically somatic sensory for upper airways) or small (Αδ-type, <3 μm in diameter, typically visceral sensory nociceptors and mechanoreceptors from the heart, vessels, lungs and GI tract) [14, 16 & 41].

Unmyelinated fibers are negative for MBP and can be either efferent or afferent.

Unmyelinated efferents are positive for NF, negative for ChAT and negative for Nav1.8. They represent non-cholinergic visceral motor fibers, some of which may be catecholaminergic (TH+), dopaminergic, or non-cholinergic-non-adrenergic. They are typically small (<3 μm) but fiber diameter is not among the criteria for classification. [8, 15, 18, 22 & 41]

Unmyelinated afferents are positive only for Nav1.8, have small diameters (<1 μm) and are arranged in clusters around nonmyelinating Schwann cells. [58, 59, 60, 98]
